# “Myosin-Driven Nuclear Movement Generates Stromules in *Arabidopsis thaliana*”

**DOI:** 10.64898/2026.10.05.756700

**Authors:** Jessica Lee Erickson, Jolina Marx, Sedef Özyürek, Simon Ortmann, Vanessa Brückner, Chaima Guizani, Alexandra Gurowietz, Carolin Alfs, Birgit Möller, Martin Hartmut Schattat

## Abstract

Plastids form stroma-filled tubular extensions, termed stromules, that are implicated in inter-organellar communication. In pathogen-challenged *Nicotiana benthamiana*, stromule formation is driven by kinesins acting along microtubules. However, whether this mechanism also operates in unstressed plants or in other species remains unknown.

Using systematic genetic analysis of all class XI myosin genes and quantitative time-lapse imaging in Arabidopsis (*Arabidopsis thaliana*), we show that basal stromule formation is predominantly myosin dependent through two distinct mechanisms. Myosin XI-I generates most stromules by driving oscillatory nuclear movement that mechanically pulls stromules from stationary plastids within the approx. 8µm wide stromule-promoting zone. Myosins XI-K, XI-1, and XI-2 additionally drive a movement-independent, nucleus-associated stromule subpopulation within the same zone, establishing two mechanistically distinct pathways. We propose that movement-coupled stromules function as dynamic tethers maintaining plastid–nucleus proximity during nuclear oscillations, while movement-independent stromules expand the plastid membrane area within the perinuclear space, increasing the potential contact surface with the nucleus and surrounding environment. Together, both populations are proposed to enhance signal and metabolite exchange at the plastid–nucleus interface.

These findings indicate that stromule biogenesis is both mechanistically and functionally context dependent, challenging the universality of the kinesin–microtubule model across plant species and physiological states.

## Introduction

Stromules are highly dynamic, stroma-filled, envelope membrane extensions that emanate from the surfaces of plastids and have been observed in plants throughout the plant kingdom (Gray et al., 2001). Although always characterized as thin protrusions (0.4 - 0.8 µm thick), stromules show a wide diversity of morphologies between tissues and species. For example, in the epidermis of *Nicotiana benthamiana* when stimulated with *Agrobacterium tumefaciens* plastids often extend kinked or branched stromules (Schattat et al., 2011; Erickson et al., 2014; Erickson et al., 2018b), while in unstressed *Arabidopsis thaliana*, epidermal plastids preferentially exhibit unbranched straight stromules (Erickson et al., 2017). Stromule length is also highly variable and can range from just a few micrometers to spanning entire cells (Kohler and Hanson, 2000; Waters et al., 2004; Erickson et al., 2014; Erickson et al., 2017).

One of the most striking features of stromules is that their formation increases under various stress conditions, with microbial responses emerging as a particularly active area of investigation. For example, in *Nicotiana benthamiana*, recognition of viral or bacterial effector proteins triggers robust immune responses in resistant plants (effector-triggered immunity, ETI), accompanied by plastid clustering and stromule formation. A subpopulation of these stromules have been found to extend toward the nucleus (Caplan et al., 2015; Kumar et al., 2018; Meier et al., 2023; Prautsch et al., 2023;). Beyond ETI, more general pathogen-associated molecular patterns (PAMPs), such as the bacterial flagellin peptide flg22, also induce transient stromule formation as part of pattern-triggered immunity (PTI). Together correlations between stromules and PTI/ETI induction have inspired two hypotheses: 1) that stromules function both in the transfer of immune-relevant signals to the nucleus (Caplan et al., 2015) and that 2) they are required for guiding plastids toward the nucleus to facilitate clustering, increasing the efficiency of immune signal transmission from plastids to nuclei (Kumar et al., 2018). This focus on stromule formation and potential function in the context of immune responses in *N. benthamiana* has neglected that the formation of stromules and nucleus-associated plastid clusters are not exclusive to plant interactions with microbes. While *N. benthamiana* pavement cells have low basal stromule frequencies and no plastid clusters prior to stress (Prautsch et al., 2023), 15-20% of plastids in unstressed *A. thaliana* upper epidermis have stromules under standard growth conditions and all cells show nucleus-associated plastid clusters (Erickson et al., 2017). Constitutive stromules and perinuclear plastid clusters in this steady state context suggest that stromules may have functions beyond immune signaling (Erickson et al., 2017). While recent years have led to a clearer understanding of how stromules might support immune responses in *N. benthamiana*, their role in cell homeostasis, as observed in *A. thaliana* upper epidermis cells, remains elusive and less well understood.

One crucial step toward understanding the biological relevance of stromules is to elucidate the mechanism(s) by which they form. In *N. benthamian*a significant progress has been made in this regard. In a previous study we established that stromules actively extend along cortical microtubules in *N. benthamiana* (Erickson et al., 2018b). Colocalization studies and speed measurements of extending stromule tips identified two distinct populations: microtubule-dependent stromules (slow-extending, 0.05-0.15 μm/s, persistent) and microtubule-independent stromules (faster, 0.24-0.28 μm/s, transient). This led us to hypothesize that kinesins, which move along microtubules, drive microtubule-dependent stromule formation, while alternative mechanisms, likely myosin-based, generate the faster, transient population. Very shortly after Kumar et al., 2018 confirmed microtubule-dependent stromule extension in the same experimental system (lower epidermis of *N. benthamiana*). Subsequently, Meier et al., 2023 identified AtKIS1, a kinesin from *A. thaliana*, as capable of inducing both stromule formation and plastid cluster formation upon heterologous overexpression in *N. benthamiana*. Since that important study, the kinesin-based mechanism is now considered the main driver of stromule formation in pathogen-challenged *N. benthamiana*. However, direct visual evidence for stromules extending along microtubules in *A. thaliana* is still lacking. Thus, whether this mechanism is conserved more broadly, including in *A. thaliana*, remains an open question.

Conversely, multiple studies implicate class XI myosin motors, in stromule formation across different tissues in *A. thaliana*, *N. benthamiana* and *N. tabacum*. Myosins are ATP-dependent motor proteins that move along actin filaments, typically faster than kinesins, and have been hypothesized to pull directly on plastid membranes during stromule formation (Gray et al., 2001, Kwok and Hanson, 2004, Natesan et al., 2009, Sattarzadeh et al., 2009, Sattarzadeh et al., 2013, Erickson et al., 2018a). In the two *Nicotiana* species, myosin ATPase inhibitors and class XI myosin silencing clearly reduced stromule frequencies (Natesan et al., 2009, Sattarzadeh et al., 2009), hinting at the myosin’s involvement in stromule formation and maintenance in hypocotyl and leaf epidermis cells. *A. thaliana* contains 17 myosins divided into classes VIII and XI (Reisen and Hanson, 2007), with class XI myosins implicated in movement of diverse organelles (Sparkes, 2011). Notably, truncations of myosin XI-2 and XI-F localize to chloroplasts and stromules in *N. benthamiana* (Natesan et al., 2009; Sattarzadeh et al., 2009) making myosins likely contributors to stromule formation. However, despite several studies investigating the connection between stromules and myosins, the role of particular myosins in stromule formation remains uncharacterized.

To understand which myosin actively supports stromule formation, we took a genetic approach in this study, screening *A. thaliana* mutants deficient in members of the myosin XI family to identify those responsible for stromule extension in unstressed *A. thaliana* upper epidermis cells. Subsequent detailed quantitative analysis of time-lapse experiments in different mutant backgrounds and inhibitor treatments revealed that two microtubule-independent mechanisms drive the majority of steady-state stromule formation in this tissue and species, establishing a new mechanistic framework for understanding plastid dynamics beyond the immune contexts.

## Results

### Myosin XI-I is required to maintain basal stromule levels in upper epidermis of *A. thaliana*

In order to investigate which of the class XI myosins contributes to stromule formation, we generated transgenic marker lines expressing a plastid-targeted eGFP (FNRtp:eGFP; Schattat et al., 2011) to label stromules. Wild-type *A. thaliana* (Columbia ecotype) and T-DNA insertion mutants in all XI-type myosins (Supplemental Information 1) were transformed and at least three independent transgenic lines were isolated for the analysis of basal stromule frequency (percentage of plastids showing one or more stromules = SF%). We focused our analysis on the upper epidermal tissue of rosette leaves, where plastid position and stromules have been well characterized before (Schattat and Klosgen, 2011; Erickson et al., 2017). Since myosins often function redundantly (Prokhnevsky et al., 2008), we also included previously characterized triple (*xi-k, xi-1, xi-2 = 3ko*) and quadruple (*xi-k, xi-1, xi-2, xi-i* = *4ko*) knockout lines that lack the most highly expressed myosins in vegetative tissues (Peremyslov et al., 2010). The majority of myosin mutants showed mean SF% between 15-24%, which is not significantly different from wild type (mean SF% of 20.8%; Figure 1A and Supplemental Information 2). The exception was *xi-i*, which showed a more substantial and clearly significant drop in SF% to 10% (p < 0.001), suggesting that myosin XI-I has a contributing role in stromule formation. The *3ko* (*xi-1*, *xi-2* and *xi-k*) mutant showed no significant difference in average SF% compared to wild type (p = 0.509; Supplemental Information 2), despite having severe defects in organelle movement and cytoskeletal structure (Peremyslov et al., 2010). Notably, individual *3ko* plants exhibited high variability. Interestingly, the addition of *xi-i* to the *3ko* mutant resulting in the *4ko* mutant, led SF% to drop dramatically to 2.7% (Supplemental Information 2). Taken together, only when XI-I was absent, either alone or in the *4ko* mutant (together with *xi-1*, *xi-2* and *xi-k*), did we observe drastically reduced stromule frequencies, highlighting the specific importance of myosin XI-I in stromule formation. This result was surprising, because although XI myosins have been implicated in stromule formation and morphology in general, myosin XI-I has never been considered important for stromule formation. In contrast myosin XI-I has an established role in nuclear movement in root tissues, with *xi-i* mutants exhibiting slow-moving, round nuclei instead of the faster-moving, spindle-shaped nuclei seen in wild type (Tamura et al., 2013). XI-I localizes specifically to the nuclear envelope through its interaction with WPP DOMAIN-INTERACTING TAIL_ANCHORED PROTEIN 1 (WIT1) and WPP DOMAIN-INTERACTING TAIL_ANCHORED PROTEIN 2 (WIT2), two integral outer nuclear membrane proteins (Tamura et al., 2013). Importantly, a double mutant combining *wit1 and wit2* (henceforth referred to as *wit12*), shows the same nuclear phenotype as *xi-i* mutants due to XI-I mis-localization (Tamura et al., 2013), making *wit12* an ideal tool to test whether XI-I’s effect on stromules is linked to impaired nuclear movement as opposed to other potential XI-I functions. However, as XI-I and WIT1, WIT2 function has only been 183 shown in roots, we first aimed to confirm the role of these proteins in nuclear movement in upper leaf epidermal cells, the focus tissue of this study.

**Figure 1.**
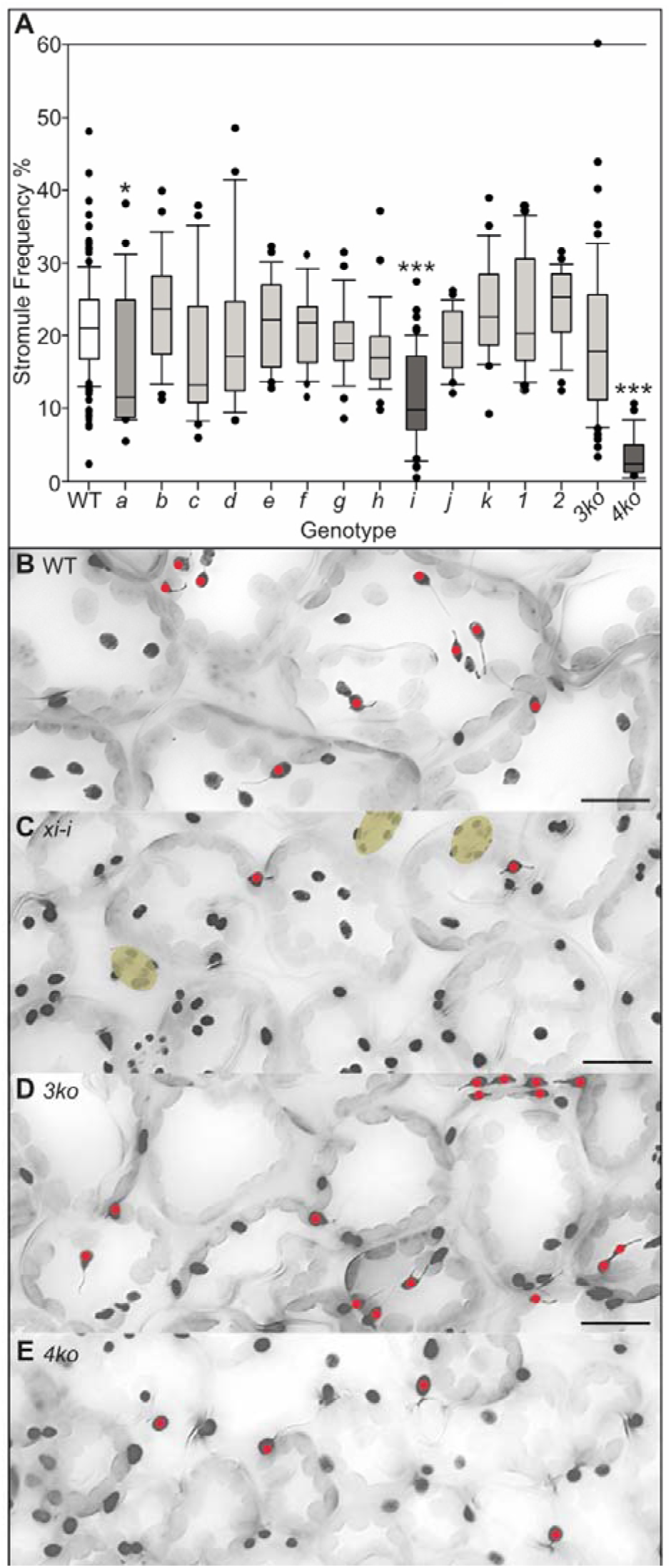
Stromule frequency is reduced in mutants lacking myosin *xi-i*. **A** Box plot of stromule frequencies in the upper epidermis of *A. thaliana* Columbia-0 (WT-white), compared to myosin xi mutants *a-k*, *1*, and *2* (light and dark gray bars), a triple mutant, harboring *xi-k*, *xi-1* and *xi-2* (*3ko*) and a quadruple mutant harboring *xi-k*, *xi-1*, *xi-2* and *xi-i* (*4ko*). All genotypes were transformed with *FNRtp:eGFP* to label plastid stroma (at least three independent transgenic lines were analyzed for each genotype, for details see Supplemental Information 1 and 2). Raw data was arcsine transformed for statistical testing and data were back-transformed for graphical representation. Boxes represent first to third quartiles, while the median is marked by a horizontal line and whiskers show the distribution of remaining data points. A Kruskal-Wallis One Way Analysis on Ranks and Dunn’s post-hoc test compared WT and mutants, significant differences are indicated with asterisks (‘*’ p<0.05, ‘***’ p<0.001). **B**-**E** Examples of epifluorescence images analyzed in **A**. **B** shows a wild-type plant, **C** a *myosin xi-i* mutant, **D** a *3ko* and **E** a *4ko* plant. For ease of visualization images were converted to gray scale and inverted, so plastids and stromules are visible in dark gray. Stomata are indicated in beige and were not part of the quantification. Plastids with stromules are marked with red dots. Scale bars are 20μm.

### *wit1_wit2*, *xi-i* and *4ko* mutants have round nuclei and reduced nuclear movement in the leaf upper epidermis

To confirm that the characterized XI-I, WIT1 and WIT2 functions in nucleus movement and morphology were not specific to roots we transformed wild type, *3ko, 4ko*, *wit12*, and *xi-i* mutants with a plastid-nucleus marker construct (*pLSU4::p35S:FNRtp:eGFP:T35S*_*pAtUBQ10:H2B:mCherry:Tnos*; Erickson et al., 2017) to visualize both organelles simultaneously. Two to three independent transgenic lines were selected for each genotype for analysis (Supplemental Information 3). Microscopic examination showed that nuclei in *4ko*, *xi-i* and *wit12* transgenic mutant plants were consistently rounder than in wild type and *3ko* plants (Figure 2 A-E), indicating the importance of the functional WIT1-WIT2-XI-I complex for nucleus shape, confirming observations in roots (Tamura et al., 2013). For a more global assessment the shape of multiple nuclei was measured as circularity index with the help of Fiji (Schindelin et al., 2012), where a value of 1.0 indicates a perfect circle while lower values indicate shapes that diverge from this optimum (Supplemental Information 4). Small nuclei were round in all lines, including the wildtype. In cells with larger nuclei differences between the genotypes were clearly visible, with elongated nuclei in wild type and 3ko plants (Figure 2 A-E), and round nuclei in mutants missing XI-I (*xi-i*, *4ko*) or when XI-I is mis-localized (*wit12*; Supplemental Information 4). Quantitative analysis of circularity dependent on nucleus size confirmed these observations and additionally revealed that in contrast to other mutants, *3ko* nuclei were significantly elongated compared to wildtype nuclei in large cells (Supplemental Information 4 and 5).

**Figure 2.**
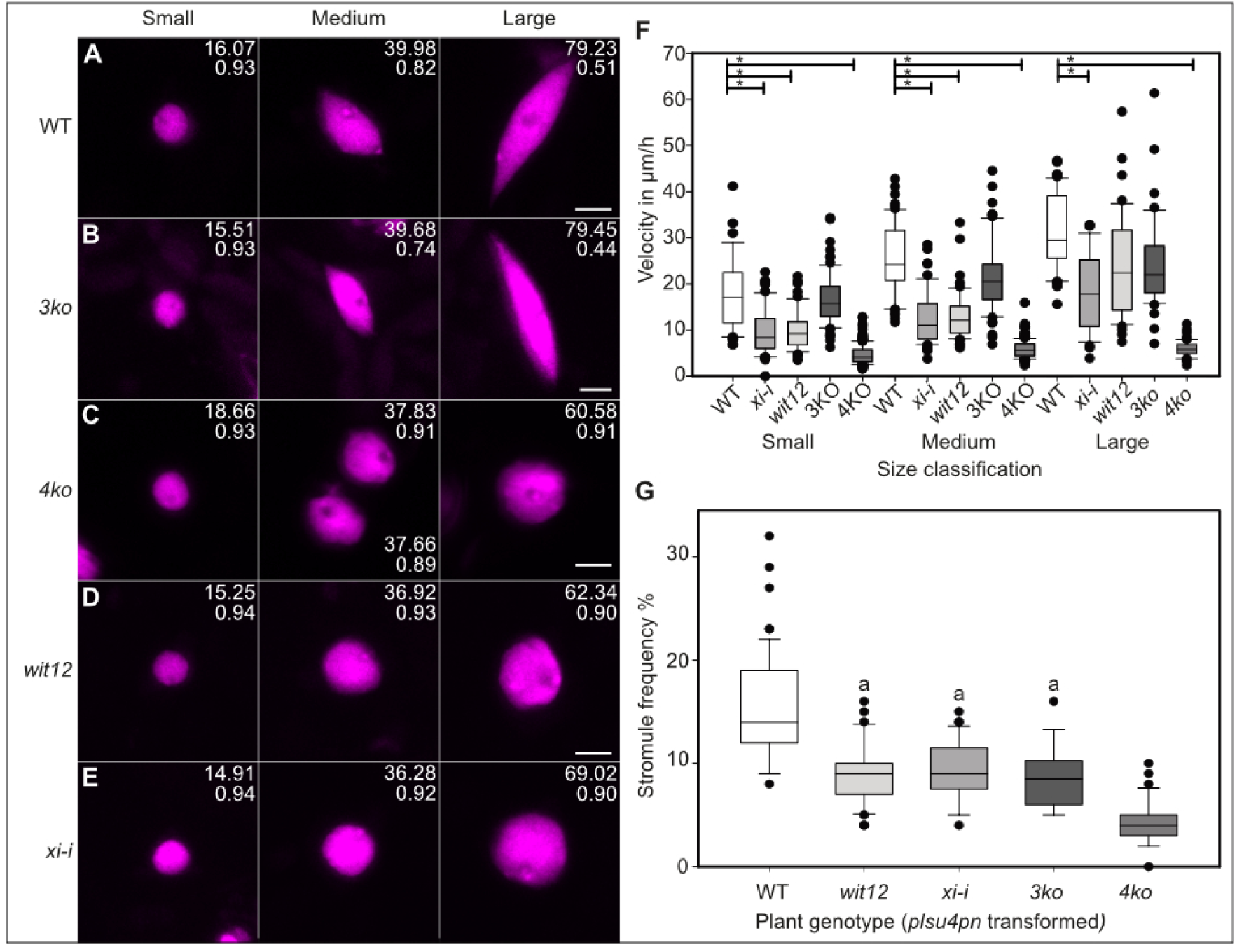
Stromule frequency in leaf upper epidermis is reduced in mutants with reduced nucleus movement. A-E) Representative confocal images (flattened z-stacks) of plants expressing H2B-mCherry to label the nucleus (visible in magenta) in A wild type Col-0, B *3ko*, **C** *4ko*, **D** *wit1_wit2* (*wit12*) and **E** *xi-i* plants from small, medium and large cell size classes. Cell size and circularity index of the nucleus depicted are shown at the top right corner. Scale bars 5 μm and are shown at the far bottom right for each genotype. **F** Nucleus velocity measurements collected from 90 min movies of plant lines depicted in **A**-**E**. Significant differences from wild type within each size class are marked with asterisks (minimum of p<0.05; Supplemental Information 8). **G** Stromule frequencies collected from epifluorescence images of plant lines depicted in **A**-**E** (3 independent *plsu4pn*, plastid nucleus organelle construct, transformants ≥9 plants). Raw data was arcsine transformed for statistical tests and data was back-transformed for the here depicted box plot (Supplemental Information 9). Treatments that are not significantly different are marked with the same letter (a). Boxes in **F** and **G** represent 50% of the data, the median is marks as a black line, whiskers represent 90% confidence intervals.

To complement the nucleus shape analysis, nuclear velocity was quantified in leaf epidermis pavement cells by tracking nuclei through CLSM time-lapse 4D datasets spanning approximately 90 minutes (30–35 frames per experiment; Video 3, 4 and 5; Supplemental Information 6). To account for potential limitation that cell size could place on nucleus movement, cells were grouped into different size classes: small (<1000 µm²), medium (1000-3000 µm²), and large (>3000 µm²) as by Erickson et al. (2017). In these experiments not all cell outlines were visible, therefore cell size was estimated based on nucleus size measurements, as these two parameters are closely correlated in *A. thaliana* upper epidermis cells of all the lines tested; Supplemental Information 7). Subsequent analysis of nuclear movement revealed that *xi-i* and *4ko* mutants showed significantly reduced nuclear speeds relative to wild type across all cell sizes, while *wit12* showed reduced speeds in small and medium cells (Figure 2F). This difference was not as pronounced in large cells (p>0.05). The *3ko* mutant also showed a slight reduction in speed, though non-significant. It is noteworthy that the *4ko* mutant exhibited the most dramatic reduction in nucleus speed, reducing nucleus movement to almost zero (Figure 2F; Supplemental information 8; Video 3, 4 and 5). Together with the nucleus shape measurements, these results demonstrate that XI-I, WIT1 and WIT2 control nucleus movement in leaf epidermis cells just as they do in root hairs (Tamura et al., 2013; Video 3, 4 and 5).

### Stromule frequency is lower in mutants with impaired nucleus movement

Having established that XI-I, WIT1 and WIT2 control nuclear movement in leaf epidermis pavement cells, we next analyzed overall SF% in mutants with slower nuclei (Figure 2G). *wit12* double mutant lines showed significantly reduced SF% (10%) compared to wild type (14%), matching the reduction seen in *xi-i* mutants (10%) and *3ko* mutants (10%), whereas *4ko* mutants exhibited the most dramatic reduction (5%) (Figure 2G). The *3ko* mutant SF% was reduced in these lines compared to the initial screen, where the statistical test did not show a significant difference (Figure 1A). However, as mentioned, in the initial screen *3ko* mutants had highly variable stromule frequencies. Decreased variability observed in this later analysis was correlated with improved control over growth conditions (compare Figure 1A and Figure 2G). Notably, the stromule frequency of *wit12* mirrored that of *xi-i*, consistent with the role of WIT1 and WIT2 as nuclear envelope adaptors linking the nucleus to myosin XI-I.

SF% values correlated with nuclear movement speeds across all genotypes: mutants with intermediate nuclear speeds (*xi-i*, *wit12*, *3ko*) showed intermediate stromule frequencies, while the 4ko mutant exhibited both the slowest nuclear movement and the lowest stromule frequency. This correlation suggests that myosin-driven nuclear movement directly promotes stromule formation, and aligns with our previously published work in this tissue (Erickson et al., 2017), in which time-lapse imaging revealed that nuclei actively move within upper epidermis cells and that stromules preferentially form from plastids within 8 µm of the nucleus, a region we termed the stromule promoting zone (SPZ; Erickson et al., 2017; Video 1). In light of this earlier observation and our stromule quantifications across the four mutant backgrounds, we hypothesized that plastid membranes are anchored near the nucleus and that nuclear movement mechanically pulls stromules from otherwise stationary plastid bodies, providing a myosin-based XI-I mechanistic explanation for the enrichment of stromules near the nucleus, resulting in the formation of the SPZ. To test this hypothesis, we performed a series of quantitative analyses to directly address the link between nuclear movement and stromule formation.

### The stromule promoting zone is altered in *xi-i, wit1_wit2* and *4ko* plants

Our initial measurements of stromule frequency were taken over entire cells and plastid position was not considered. So although it was clear that stromule frequencies are correlated with nucleus movement in the upper epidermis, it was not yet clear if the SPZ was specifically disrupted. If nucleus movement drives the formation of stromules among peri-nuclear plasids, we expected a reduced SPZ in nucleus movement mutants. To test for the effect of mutations on the SPZ specifically, stromule frequency was analyzed as a function of plastid distance from the nucleus.

In order to analyze the SPZ, four key parameters have to be measured: cell outlines, cell size, plastid counts (with and without stromules) as well as the distance between each plastid and the nucleus (Figure 3A and B). This analysis involves the examination of hundreds of cells across three independent transgenic lines for each mutant and the wild type. To efficiently process this volume of data, we developed an automated image analysis pipeline combining an enhanced version of the MTB Cell Counter Plugin (Franke et al., 2015; Supplemental Methods 1).

**Figure 3:**
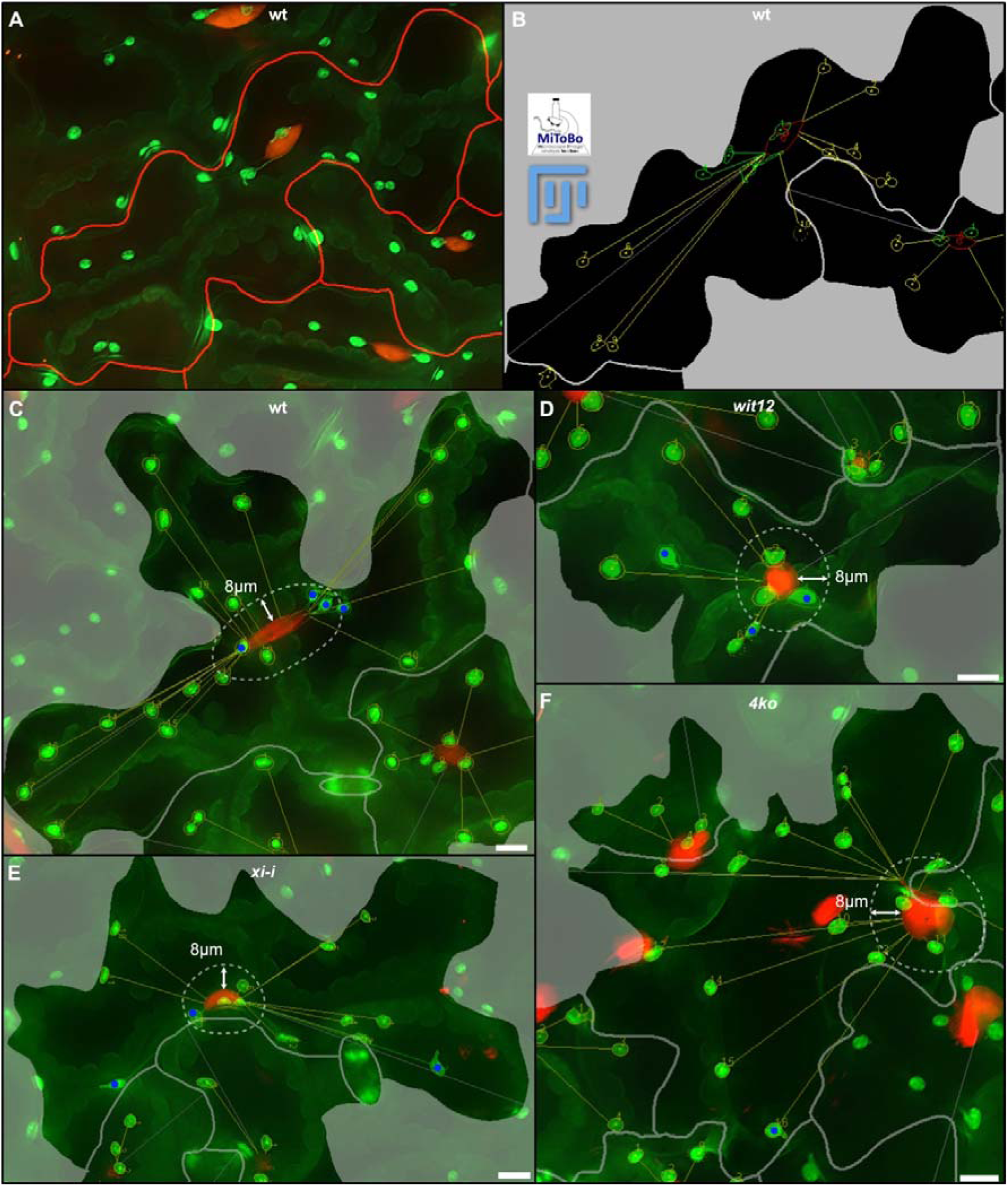
A custom image analysis tool supports collection of quantitative data for stromule promoting zone measurements. **A** Shows a section of a stacked microscope image of an upper epidermis cell of a *A. thaliana* wildtype line expressing FNRtp:eGFP (green) as well as H2B:mcherry (red). Cell outlines are marked by a red overlay line. **B** graphical output of the MiToBo runner plugin of the same image section following the detection of nuclei (red outline), plastids without (yellow outline) and plastids with stromules (green outline). The graphical and numerical output includes cell size, number of plastids with and without stromules, the distance of each plastid to the respective nucleus and the maximal possible distance from the nucleus (thin grey straight line) in each cell. **C - F** Sample images of the overlay of the runner output and stacked microscopic images for the different genetic backgrounds with indicated 8 µm zone (dashed line) and labeled stromule harboring plastids (blue dots). Scale bar = 10 µm.

To analyze the impact of mutations on the SPZ, we grew three independent transgenic plastid-nucleus marker lines each of *xi-i, wit12*, and *4ko* mutants alongside wild-type controls. We imaged the upper epidermis of 2-6 plants per line (Supplemental Information 10). The *3ko* mutant was not included in this analysis as it showed the weakest impact on nucleus velocity (Figure 2F) and the least consistent impact on stromule frequency.

Prior to analyzing the SPZ, it was necessary to establish whether the nuclear movement mutations (*4ko*, *xi-i*, and *wit12*) also affect how plastids are distributed relative to the nucleus. Therefore, plastid distribution was analyzed across genotypes by plotting histograms of plastid numbers in 4 µm-wide distance bins for small (<1000 µm²), medium (1000–3000 µm²), and large (>3000 µm²) cells (Supplemental Information 11). The distribution patterns were remarkably similar across all genotypes, with the highest proportion of plastids consistently found within the SPZ. Only *4ko* mutants showed slightly fewer plastids in this zone in small and medium cells (Supplemental Information 12), suggesting that loss of all four myosins may have a minor effect on perinuclear plastid positioning. The overall conservation of plastid distribution patterns across genotypes suggests that differences in the SPZ would reflect genuine differences in stromule formation propensity rather than a redistribution of the plastid population. Additionally, also demonstrates that the enrichment of plastids near the nucleus that is consistently observed in this tissue (Erickson et al., 2017) does not depend on the myosin XI–WIT1/WIT2 nuclear movement machinery.

Having confirmed that plastid distribution is similar in the different genetic backgrounds, SF% within the SPZ was then analyzed. Given that nuclear movement varies with cell size (Figure 2F), we analyzed the SPZ separately for each cell size category. As seen before (Erickson et al., 2017) we observed in wildtype plant lines a clear 8µm SPZ in both large and medium cells (Figure 4A). Large cells showed higher SF% within this zone (4 µm zone: 38%; 8 µm zone: 24%) compared to medium cells (4 µm zone: 21%; 8 µm zone: 17%). When moving beyond the SPZ, SF% drops dramatically. Small cells, which have the least nuclear movement (Figure 2F), also showed no distinct SPZ, with SF% consistently below 10%. *xi-i* and *wit12* mutants showed a dramatic changes to the SPZ stromule frequency within 4µm of the nucleus (*xi-i*: 17%; *wit12*: 20%) in large cells and in medium cells, the SPZ disappeared entirely. Small cells, as was the case in wild-type, showed no promoting zone. The *4ko* mutant showed the most drastic phenotype, with no SPZ in any cell size. However, stromule frequency beyond the 8µm zone remained similar to the wildtype and other genotypes (Figure 4D; for numerical values see Supplemental Information 13 and 14). Thus, in mutants with reduced nucleus speed the SPZ collapses.

**Figure 4:**
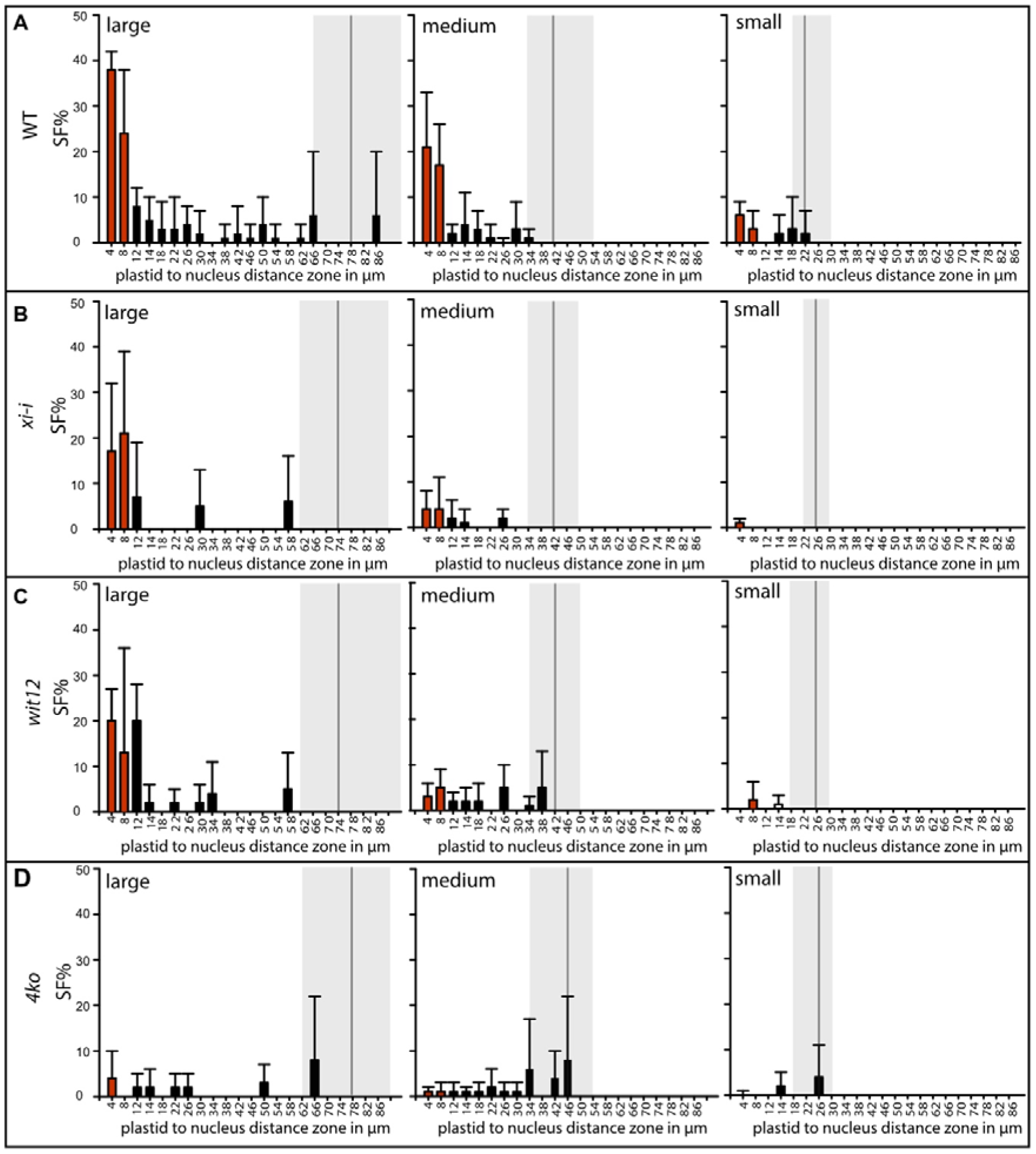
Stromule frequency analysis reveals altered stromule promoting zones in nucleus movement mutants. A-D) Average stromule frequency (SF%; error bars = standard deviation) across distance zones from the nucleus (4 µm intervals, labeled at midpoint) for large (>3000 µm²), medium (1000-3000 µm²), and small (<1000 µm²) cells (red bars = stromule promoting zone). A) Wild type shows clear stromule enrichment within 8 µm of the nucleus in large and medium cells. B) *xi-i* single mutant exhibits reduced stromule frequency in the 8 µm zone. C) *wit12* double mutant shows similar reduction to *xi-i*. D) *4ko* mutant (*xi-k/xi-1/xi-2/xi-i*) displays severely reduced stromule frequency across all distances and cell sizes. Gray shaded areas with black vertical line represent the median and the lower as well as upper quartile of maximum possible distances from nucleus to cell wall for the respective cell size class.

To test the degree of correlation between nuclear movement and stromule formation, we plotted nuclear speed against SF% among plastids 4 µm, 8 µm or >8 µm from the nucleus across genotypes (Figure 5A-C). Linear regression analysis revealed strong correlation between both parameters among plastids in the 4 µm zone, followed by the 8 µm zone, while the correlation beyond 8 µm was negligible. This analysis confirms that SF% within the promoting zone directly correlates with nuclear movement speed, while stromules outside this zone are unaffected by nuclear movement. Together with the SPZ measurements, this establishes a close correlation of nucleus speed and stromule numbers in the SPZ.

**Figure 5:**
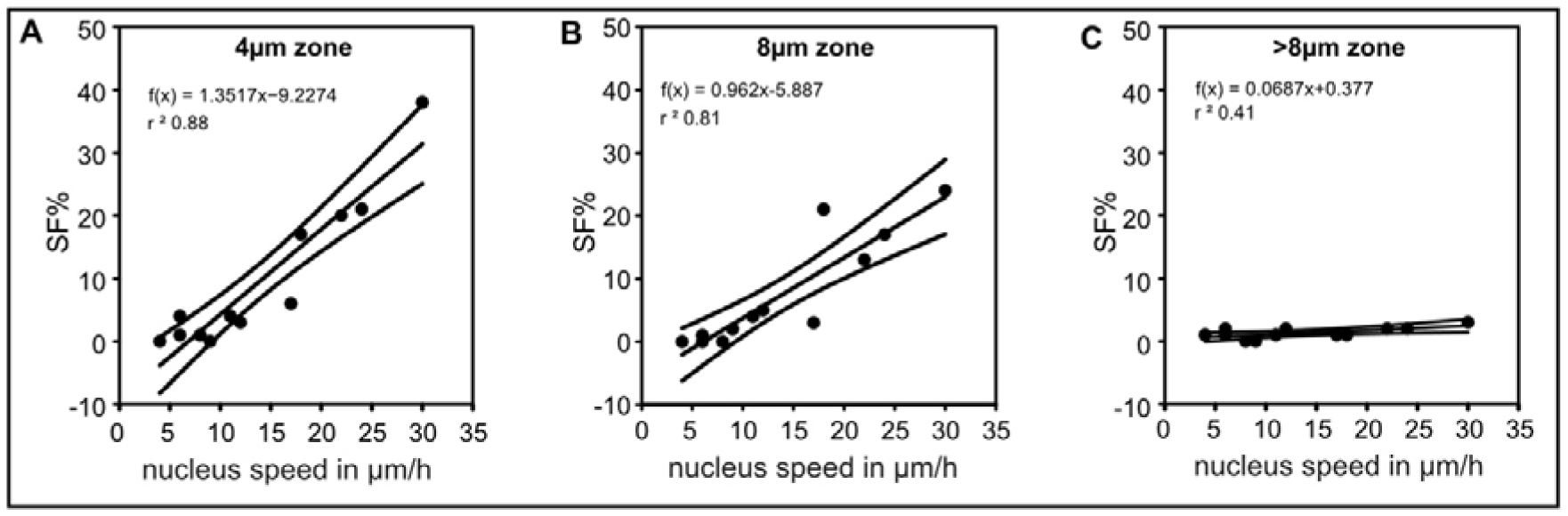
In the stromule promoting zone stromule frequency (SF%) and nucleus speed follow a positive linear correlation. A – C Scatter plot of SF% and nucleus speed within the stromule promoting zone (A and B) and outside the stromule promoting zone (C). C represents all plastids further than 8 µm away from the nucleus. straight line = linear regression line; curved lines = 95% confidence intervals; r^2^ = correlation coefficient; f(x) = linear regression equation. Scatter plot consider all cell sizes (large, medium and small) as well as genetic backgrounds (wt, *xi-i*, *wit1_wit2* and *4ko*). For plots separated by cell size see Supplemental Information 16.

### The majority of stromule forming events at the nucleus coincides with its movement

Our findings revealed a strong correlation between nuclear velocity and stromule frequency within the SPZ, we sought to obtain direct evidence linking stromule formation in this zone to nuclear movement. To this end, we employed confocal laser scanning microscopy to capture time-lapse sequences of the upper epidermis of wildtype lines over approximately 90 minutes, enabling continuous observation of nuclear and plastid behavior (representative sequence shown in Video 1).

In these recordings, individual nuclei exhibited oscillatory movement around a central position, while nearby plastids remained largely stationary (also see Figure 6A, B and Video 2). Notably, when nuclei moved away from their associated plastids, these plastids frequently extended stromules in synchrony with nuclear displacement, stretching in the same direction and over a comparable distance but membranes did not always appear to be in direct contact (Figure 6B, Video 2). These recordings supported the hypothesis that the stromule formation near the nucleus is likely a direct mechanical consequence of nuclear movement (Figure 6A, B; Video 2) and show that the attachment of stromule tips is likely not directly at the nucleus surface. Most but not all stromule formation events were synchronized with nuclear movement. For example, plastids distant from the nucleus (outside the SPZ) displayed stromule formation that occurred independently of nuclear displacement (Figure 6C, Video 2). On the other hand, such asynchronous stromule formation was occasionally observed even in plastids positioned near the nucleus (within the SPZ), suggesting that multiple stromule-inducing mechanisms operate forming stromules outside and within the SPZ (Figure 6D, Video 2).

**Figure 6:**
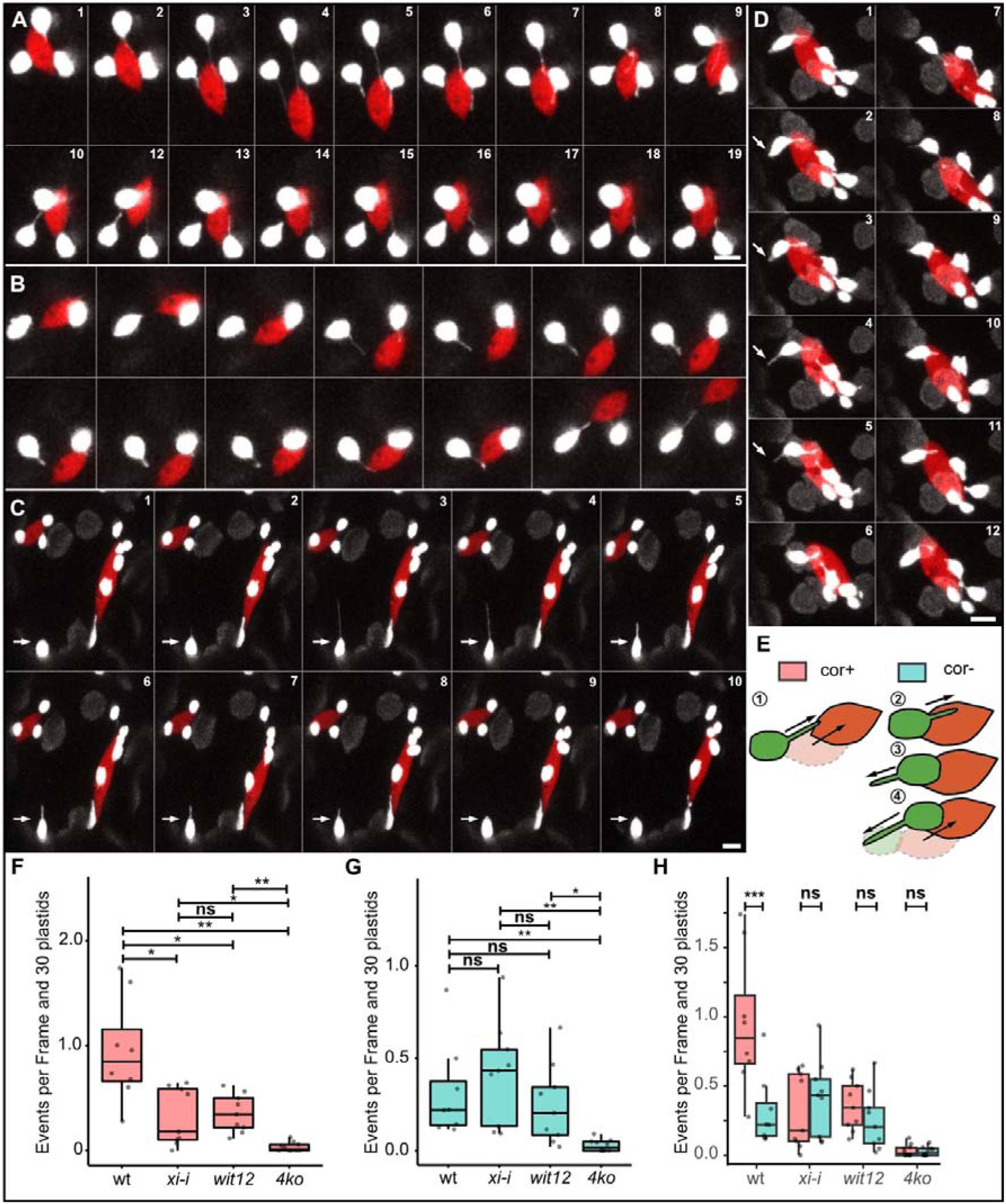
Stromule formation near the nucleus predominantly coincides with nuclear movement. (A–B) Time-lapse sequence illustrating synchronized stromule (white) formation and nuclear (red) movement. (A) stromule anchored possibly at the nucleus; (B) stromule anchored close to the nucleus (visible gap between stromule tip and nucleus); (C) plastid located distant from the nucleus forming a stromule independently of nuclear movement; (D) Example of asynchronous stromule formation in a nucleus-associated plastid; (E1–E4) Schematic overview of stromule formation categories. cor+ (E1): stromule formation coinciding with nuclear movement in both direction and distance. cor– events: asynchronous stromule formation while the nucleus remains stationary, either extending toward (E2) or away from (E3) the nucleus, or occurring during nuclear movement without matching direction or distance, including cases where plastids move together with the nucleus (E4). (F–H) Quantitative analysis of stromule formation events per frame normalized to 30 plastids across wild type (wt), *xi-i*, *wit1_wit2*, and *4ko* (F–G) Quantitative comparison of stromule formation categories across genotypes; (F) cor+ events; (G) cor–; (H) Comparison of cor+ and cor– events within each genetic background. For (F) and (G), between-genotype differences were assessed using the Kruskal-Wallis test followed by pairwise Wilcoxon rank-sum tests with Holm correction (non-parametric tests were used as ANOVA assumptions were not met; see Supplemental Information 17). For (H), within-genotype differences between cor+ and cor– were assessed using a paired t-test, as the paired differences did not deviate from normality (Shapiro-Wilk test; see Supplemental Information 17). Statistical significance is indicated by: ns = no difference; *p < 0.05; **p < 0.01; ***p < 0.001. Scale bars = 5 µm (A–D).

To more directly assess the extent to which nuclear movement contributes to stromule formation within the SPZ, we quantified the proportion of stromule formation events coinciding with and without nuclear displacement. Stromule events were categorized as *cor+* (correlation positive) when stromule formation matched both the direction and distance of nuclear movement (Figure 6E1), or as *cor–* (correlation negative) when one or both of these parameters did not align. The *cor–* category included stromules formed while the nucleus remained stationary, either extending toward (Figure 6E2) or away from it (Figure 6E3), as well as cases where stromule formation occurred during nuclear movement but without matching direction or distance, sometimes including plastids that moved together with the nucleus (Figure 6E4).

Across the genetic backgrounds of wildtype and the key mutants (*xi-i*, *wit12* and *4ko*), a total of approximately 57 hours of imaging time were analyzed, encompassing 1075 nucleus-associated plastids and documenting 727 stromule formation events. Stromule formation events per frame and 30 plastids, are presented in Figures 6F–H (see also Supplemental Information 17). In wild-type plants, *cor+* events clearly dominated, occurring almost 4 times as frequent (*cor+* wt = 0.85 vs. *cor-* wt = 0.22), as *cor–* events and showing a statistically significant difference (Figure 6H; Supplemental Information 17). In contrast, in *xi-i* and *wit12* mutants—both impaired in nuclear movement to the same degree, the frequency of *cor+* events dropped sharply to the same level as *cor–* events. In the *4ko* mutant (lacking *xi-1*, *xi-2*, *xi-k* and *xi-i*), both *cor+* and *cor–* events were nearly abolished together with nucleus movement (see Figure 2F) and occurred at similarly low frequencies, showing no difference (Figure 6H).

This detailed video analysis supports the hypothesis that nuclear movement is a major driver of stromule formation within the SPZ, with myosin XI-I playing a key role. Further, approximately one-quarter of stromule formation events occurred independently of nuclear movement and did not rely on XI-I activity, indicating the presence of a separate, nucleus-movement-independent mechanism. The near-complete loss of both event types in the *4ko* mutant suggests that the remaining stromule formation observed in *xi-i* and *wit12* depends on one or more of the myosins absent in the *4ko* mutant, XI-1, XI-2, and XI-K. Further, this reveals that myosins are required for both nucleus movement-dependent and independent stromule formation.

### Nuclear movement explains the stromule promoting zone’s reach

The quantification of type-specific stromule formation frequency demonstrated the dominant importance of nuclear movement for stromule formation within the SPZ. However, this correlation alone does not explain the extent of the zone, which we consistently measured to be approximately 4 to 8 µm in size. A possible explanation for this reach may lie in the oscillatory movement patterns of the nucleus, clearly visible in our time-lapse recordings (Video 1; and illustrated in Figure 7A). In these sequences, stromules are initiated as the nucleus moves away from nearby plastids, elongate during movement, and retract as the nucleus reverses direction, suggesting that the amplitude of nuclear oscillation defines the typical stromule length and, consequently, the effective range of the SPZ.

**Figure 7:**
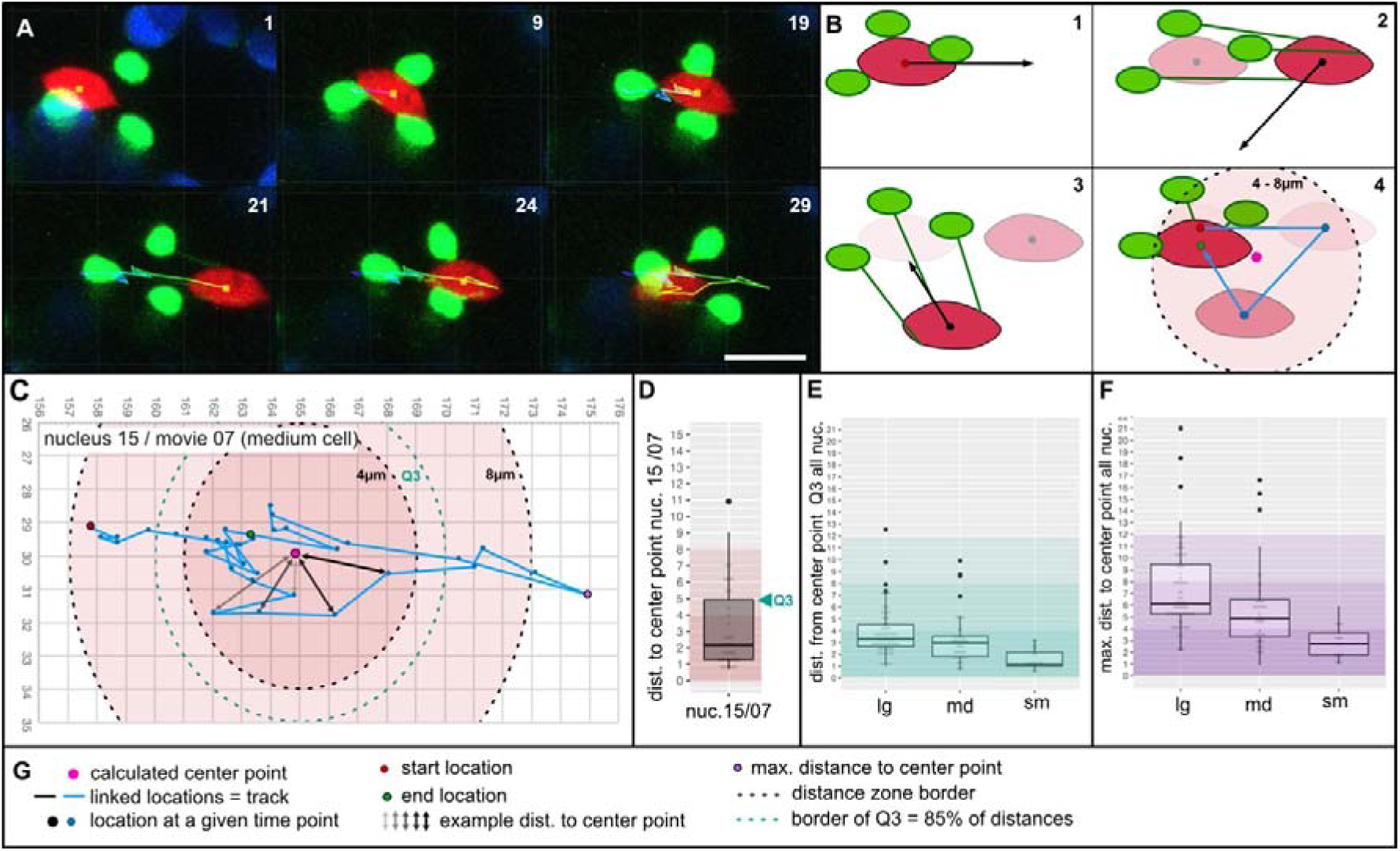
Nuclei describe an oscillating movement pattern, which correlates in its amplitude with the measured stromule promoting zone. A) snap shots of a time series, capturing the oscillating movement pattern of a nucleus; colored line represents the track of nucleus center point; number indicate frame number; scale bar = 10 µm; B) Schematic, depicting nucleus movement, stromule attachment to nucleus, calculated center point of nucleus track and resulting stromule promoting zone; C) sample track of a tracked nucleus with calculated center point, Q3 distance to center point, 4µm as well as 8µm zone boundary (measured from nucleus track center point); D) sample box plot for nucleus 15 movie 07 showing the measured distances of the nucleus for each time frame from the nucleus track center point (nucleus track shown in C); Q3 represents the 85 percentile of these measurements and means that in 85% of all time points the nucleus is closer to the center point than this value; E) box plots capturing the distribution of all Q3 values of nuclei tracks in large (lg), medium (md) and small (sm) cells; F) box plots capturing the distribution of the maximum distance from the center point of all large (lg), medium (md) and small (sm) cells; D – F) color gradients mark the 4 µm, 8 µm and >8 µm distance border.

If this mechanism applies across the entire cell population, the amplitude of the oscillating nucleus, that is, the distance the nucleus travels away from its central position before changing direction, should correspond to the width of the SPZ (Figure 7B). To test this, we determined whether wild-type nuclei exhibited oscillatory movement and whether their oscillation amplitudes matched the previously defined SPZ. For each tracked nucleus, we (1) calculated its central position of movement, (2) measured the distance from this center point over time, and (3) assessed whether these distances were within the previously defined SPZ boundaries (4 µm in small cells and 8 µm in medium and large cells).

Figure 7C presents a representative example, nucleus 15 from movie_07, situated in a medium-sized cell. We found that across 28 out of 30 tracked points along the trajectory of this specific nucleus fell within 8 µm of the center of oscillation, matching the region defined as the SPZ. The same can be also seen when distances from the oscillation center point are plotted as a boxplot (Figure 7D). In order to quantitatively compare as many nuclei as possible we followed the same strategy for 130 nuclei (Supplemental Information 18), calculating the distance from the oscillation center point across all time frames and visualized these as individual box plots. (Supplemental Information 19). We quantified nuclear behavior using two metrics: the radius containing 85% of each nucleus’s tracked positions (referred to as Q3 (3^rd^ quartile); a measure of the region where nuclei spent most of their time) and the maximum distance traveled from the calculated center point (Figure 7D). Nuclei were grouped by cell size category, and the Q3 (Figure 7E) and the maximum travel distance (Figure 7F) for each category was plotted. The plots reveal that almost all nuclei (96%) spent at least 85% (Q3) of their time within the SPZ boundary (8 µm for medium and large cells, 4 µm for small cells). Only 5 of 130 nuclei (2 in large cells, 3 in medium cells) spent more time outside this zone (Figure 7E). Even when examining maximum distances, most nuclei never left the SPZ during the entire observation period (large cells: 71%; medium cells: 83%; small cells: 100%; for all plots see Supplemental Information 19 and 20).

These findings led us to conclude that the nucleus’s oscillation amplitude is indeed representative of the reach of the SPZ, providing further evidence that nuclear movement is the driving force of stromule formation and even defines the SPZ.

### Nucleus-associated actin provides the anchor for stromule formation

Now that we had established that myosin driven oscillating nuclear movement occurs within an 8µm radius, driving the majority stromule formation within the SPZ and by that defining it. One key aspect of this system remained unaddressed: what serves as the anchor point for stromule tips at the nucleus? Understanding this anchoring mechanism is crucial, as identification of the specific anchoring structures would enable the generation of mutants lacking these components, potentially eliminating SPZ stromules and allowing investigation of their physiological significance.

Previous work in *N. benthamiana* lower epidermis suggested roles for both actin and microtubules in stromule-mediated plastid-nucleus interactions, with microtubules directing stromule growth and actin providing anchor points at stromule tips and along stromules, particularly at the nucleus (Kumar et al., 2018). To determine which cytoskeletal elements might serve as anchoring structures in Arabidopsis, we characterized the three-dimensional arrangement of nuclei (DAPI stained), plastids (Chlorophyll autofluorescence), and cytoskeletal (GFP::MAP4 = microtubules or eGFP::ABD2::eGFP = actin) elements in *A. thaliana* upper epidermis using super-resolution confocal imaging (Lightning Processing provided by the LasX Software from Leica, Figure 8A-D). We observed that microtubules did not form a cage around nuclei but were primarily located in the cell cortex (Figure 8C). Consequently, microtubule-nucleus contacts were only visible where nuclei were pressed against the cell periphery (Figure 8D; Video 6 and 7). Only on rare occasions did individual filaments diverge from this constellation (Video 8). We observed no microtubules at plastid-nucleus interfaces, suggesting they do not facilitate nucleus anchoring of plastids (Figure 8D).

**Figure 8:**
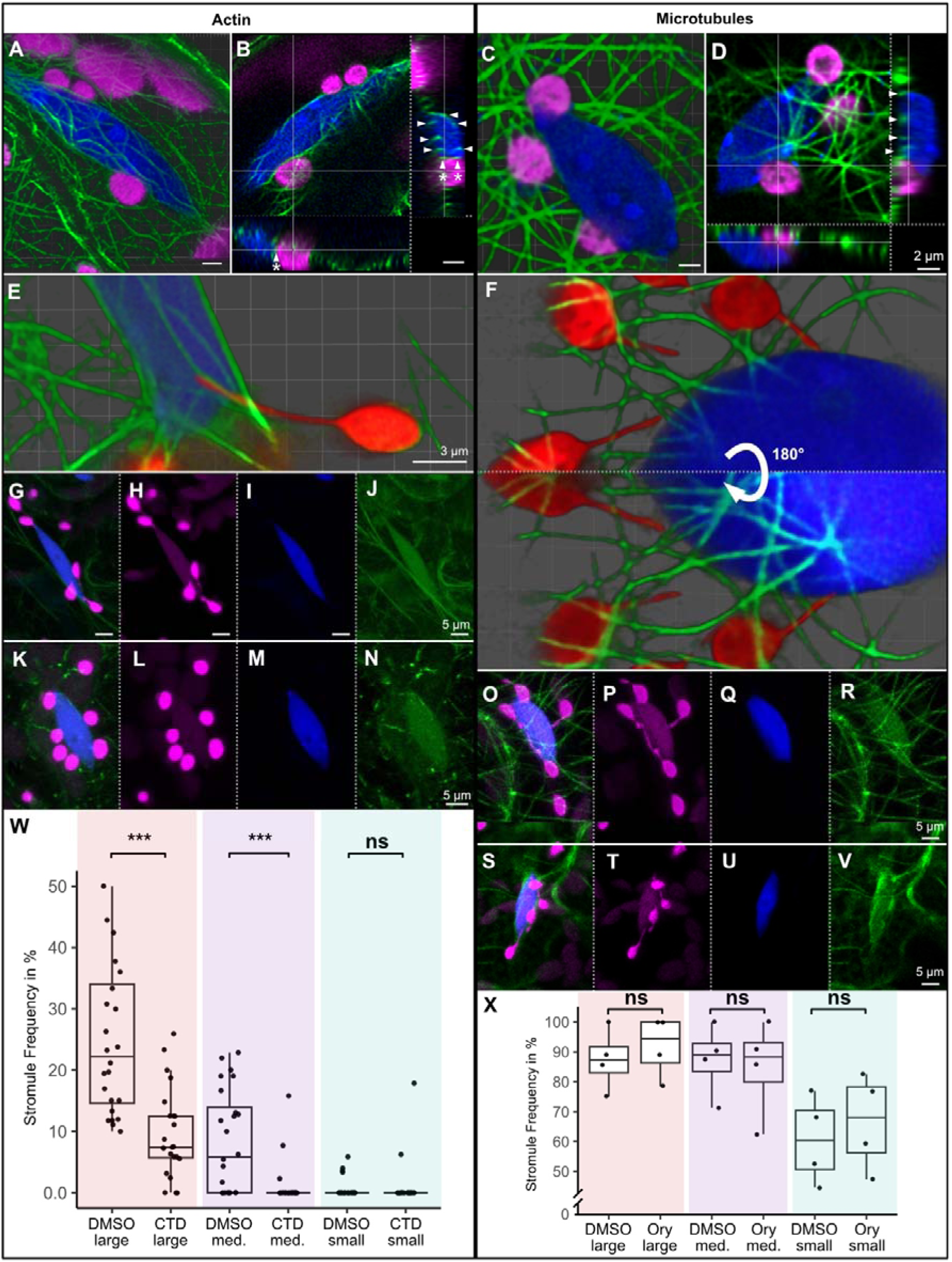
Nucleus-associated actin, not microtubules, anchors stromule tips at the nuclear periphery. A–B) Three-dimensional rendering (A) and orthogonal view (B) showing actin filaments completely encaging the nucleus (arrowheads) and forming contact points with plastids (arrowheads marked with asterisks). C–D) Equivalent visualizations of microtubule-labeled plant material show that microtubules contact the nucleus only on the cell wall-facing side (C,D; arrowheads). Green = actin (A,B) or microtubules (C,D), magenta = chlorophyll, blue = DAPI. Scale bars = 2 µm. E) Three-dimensional rendering of a deconvolved (Lightning) confocal z-stack showing close association between a stromule tip and nuclear periphery actin. F) Three-dimensional rendering of a deconvolved (Lightning) confocal z-stack showing stromule tips in close proximity to, but not in contact with, microtubules. 3D reconstruction is shown from two viewing angles (0° and after 180° rotation). Green = actin (E) or microtubules (F), blue = nucleus, red = plastid stroma. Scale bar = 3 µm. G–N) Maximum intensity projections showing the impact of 2-hour cytochalasin D treatment on actin and stromules. DMSO control (G–J) maintains intact actin filaments and SPZ stromules, while cytochalasin D treatment (K–N) disrupts actin filaments and causes stromule retraction. Green = actin, blue = nucleus, magenta = plastid stroma. Scale bar = 5 µm. O–V) Maximum intensity projections showing the impact of 2-hour oryzalin treatment on microtubules and stromules. DMSO control (O–R) maintains intact microtubules and SPZ stromules, while oryzalin treatment (S–V) disrupts microtubules but leaves stromule dynamics unaffected. Green = microtubules, blue = nucleus, magenta = plastid stroma. Scale bar = 5 µm. W) Stromule frequency in the 4 µm stromule promoting zone following DMSO control or cytochalasin D (CTD) treatment, across cell size classes. Significant reduction in stromule frequency occurs in large and medium cells (p < 0.001), while small cells show minimal stromule frequency regardless of treatment. X) Stromule frequency in the 4 µm stromule promoting zone following DMSO control or oryzalin (Ory) treatment, across cell size classes. No significant difference between treatment and control was observed in any cell size class. Box plots show quartiles and median; whiskers indicate 10th/90th percentiles.

For clear visualization of actin filaments, including smaller structures, we briefly treated samples with CCCP (Carbonyl cyanide m-chlorophenyl hydrazone) to halt actin remodeling by depleting cellular ATP (Felle and Bentrup, 1977). In contrast and consistent with previous studies (Iwabuchi et al., 2010), we observed a dense network of actin filaments surrounding nuclei (Figure 8A). Crucially, we frequently detected actin filaments at plastid-nucleus contact interfaces (Figure 8B), providing potential anchor points for nucleus interactions (Video 9 to 11). Together, these observations identify actin as the most likely anchor point for stromule tips to moving nuclei.

To obtain more direct evidence for actin- or microtubule-mediated stromule-nucleus interaction, we established triple-marker transgenic lines simultaneously visualizing actin or microtubules, nuclei, and the plastid stroma. As fluorescent proteins we used EGFP for actin or microtubules, mKOK for nuclei and E2-Crimson for the plastid stroma, fused with well-characterized targeting sequences (Grebenok et al., 1997; Marc et al., 1998; Kohler et al., 1997; Wang et al., 2008; for details on those constructs see Supplemental Methods 3). In these transgenic lines, actin and stromule tips were frequently found in close proximity at or near the nuclear surface (Figure 8E; Supplemental Information 21 and 22; Video 12 to 16). In contrast, stromule tips came into clear contact with microtubules near the nucleus in only a single instance (Figure 8F; Supplemental Information 23; Video 17 and 18). Together, these observations pointed to actin rather than microtubules as the more likely anchor for stromule tips at the nucleus.

To test this directly, we treated plants with the actin-depolymerizing drug cytochalasin D (4 μM) or the microtubule-depolymerizing drug oryzalin (100 μM) as well as respective solvent controls (0.2 % DMSO for oryzalin, 0.1 % DMSO Cytochalasin D), reasoning that disruption of the relevant anchoring element should release stromules from the nucleus, causing their retraction and reducing stromule frequency within the SPZ. Time-lapse imaging tracked this process (cytochalasin D: Video 19 = DMSO control and 20 = cytochalasin D treatment; oryzalin: Video 21 = DMSO control and 22 = oryzalin treatment). In DMSO control conditions, both the actin and microtubule networks remained intact throughout the observation period. After 2-hour drug treatments, most actin filaments or microtubules were disrupted following chemical treatments, though some small cytoskeleton fragments remained in both cases (cytochalasin D: Figure 8G–N; oryzalin: Figure 8O–V). The two treatments produced strikingly different outcomes. In cytochalasin D-treated cells, stromule retraction within the SPZ was frequently observed in time-lapse recordings (Video 19 and 20), and quantification revealed a clear reduction in SPZ stromule frequency across all three cell size classes (Figure 8W, Supplemental Information 25). In contrast, oryzalin treatment had no detectable effect on stromule frequency in the SPZ in any cell size class (Figure 8X, Supplemental Information 24), and stromule dynamics, both nucleus-dependent extension and retraction and nucleus-independent events, persisted throughout the treatment period (Video 21 and 22).

One could argue that the reduction in stromule frequency following cytochalasin D might simply reflect inhibited nuclear movement rather than loss of anchoring points. However, halting nucleus movement by actin depolymerization with no impact on stromule anchoring would presumably leave the SPZ unchanged, frozen in time. Instead, the observed collapse of SPZ stromules indicates that actin maintains stromule attachment at the nucleus.

Taken together, the three-dimensional imaging of cytoskeletal-plastid-nucleus arrangements, the colocalization of stromule tips with nuclear-associated actin, and the contrasting effects of cytochalasin D and oryzalin provide strong evidence that nucleus-associated actin, and not microtubules, serves as the anchor for stromules within the SPZ.

## Discussion

### Myosins are important contributors to stromule elongation

In recent years, the prevalent model of stromule formation was formed based on observations in immunity triggered *N. benthamiana* plants and subsequent research has revolved around this idea. Leading to a focus on the involvement of microtubules and kinesin motor proteins as the primary driving force for stromule formation (Caplan et al., 2015; Kumar et al., 2018, Meier et al., 2023). Within the frame of this model actin has been suggested to play a role in stromule stability rather than playing an active part in their formation, anchoring stromules along their length and at the periphery of the nucleus (Kumar et al., 2018). Surprisingly, our investigations reveal that the mechanisms of stromule formation stemming from work in *N. benthamiana* epidermal cells under stress does not explain stromule formation observed near to the nucleus in epidermis cells in *A. thaliana* under non-stressed conditions. We found that under these conditions stromule formation in the SPZ is likely governed not by kinesins, but instead by myosin motors. Our analysis reveals that their extension occurs via two distinct mechanisms. The majority of perinuclear stromules are generated indirectly, by myosins driving nucleus movement, which exerts mechanical force on the plastid envelope through actin attachment points at the nuclear periphery, pulling out stromules as the nucleus undergoes oscillatory movement. In addition, a subpopulation of approximately 25% of perinuclear stromule formation events occurs independently of nuclear movement, yet still depends on myosin activity, specifically that of XI-K, XI-1, and/or XI-2, pointing to a parallel, potentially direct actin-myosin mechanism at the nuclear periphery. Therefore, in addition to the kinesin-based models of stromule extension, this work establishes two additional stromule-forming mechanisms which are myosin-mediated. This demonstrates the importance of considering species specific, tissue specific and environment specific stromule-forming mechanisms and functions. Additionally, this raises the immediate question of whether kinesins play any role in stromule formation in *A. thaliana*.

### What is the role of kinesins in stromule formation in *A. thaliana*?

The evidence for kinesin/microtubule-dependent stromule formation rests mainly on two complementary lines of evidence from *N. benthamiana*: 1) the clear induction of stromules upon heterologous overexpression of kinesin AtKIS1 (Meier et al., 2023), and 2) time-lapse imaging co-visualising stromules and microtubules combined with pharmacological inhibitor treatments (Erickson et al., 2018b; Kumar et al., 2018), together providing strong evidence for a kinesin/microtubule-based mechanism in this species. Similarly detailed experimental evidence in *A. thaliana* is entirely absent. Stromule frequency has only been examined in plants challenged with ETI-inducing *Pseudomonas syringae* strains, where comparisons were made between *A. thaliana* wildtype and T-DNA insertion lines with disrupted AtKIS1 (*Atkis1-1* and *Atkis1-2*) (Meier et al., 2023). In these experiments both mutant lines exhibited lower stromule frequencies than wildtype, but whether this is a result of less AtKIS1-mediated stromule extension or an indirect consequence of knocking out the kinesin is unclear. Critically, baseline stromule frequency data for the *Atkis1* mutants under non-inoculated control conditions was not investigated (Meier et al., 2023), meaning we have little insight into the steady-state conditions of the SPZ.

Intriguingly, our three-channel movies co-visualisating stromules, microtubules, and nuclei revealed a single instance in which a stromule seemed to be initiated to form along a microtubule near the nucleus (Video 17, circle). This hints that microtubule–plastid contact and extension may actually occur in *A. thaliana* leaf epidermis, but as shown by our inhibitor treatments might be not of great importance to the SPZ in non-stressed conditions. Importantly such a microtubule-based mechanism is not excluded by our myosin-driven SPZ model. While stromules in close proximity to the nucleus are largely a consequence of myosin activity, stromule formation outside this zone also remains unexplained. Our data suggest these distal stromules do not rely on XI-I, XI-1, XI-2, or XI-K, leaving the possible involvement of AtKIS1, other kinesins, or additional myosins as possibilities. Thus, the direct involvement of kinesins in stromule formation in *A. thaliana* remains still unclear. Clarifying the role of microtubules-based movement for stromule formation in *A. thaliana* requires the same rigorous experimental approaches applied in *N. benthamiana*, co-visualisation of stromules and microtubules in time-lapse imaging combined with pharmacological inhibitor treatments and systematic mutant analysis, ideally conducted under both steady-state and ETI-induced conditions.

### Could myosin-driven nucleus movement also be important for *N. benthamiana*?

Myosins have long been implicated in stromule formation in *Nicotiana* species: actin and myosin inhibitor treatments reduce stromule frequencies in tobacco hypocotyls (Kwok and Hanson, 2003), and RNAi-mediated silencing of myosin XI in *N. benthamiana* reduces stromules in leaf epidermal cells (Natesan et al., 2009; Sattarzadeh et al., 2009), although the precise mechanism was never elucidated. Our discovery of myosin-driven nuclear movement as a dominant mechanism of stromule formation in *A. thaliana* therefore raises an important reciprocal question: might an equivalent mechanism be operating, and have been overlooked, in *N. benthamiana*? Nucleus-associated stromule formation during immune responses has primarily been described in terms of microtubule- and kinesin-based mechanisms (Erickson et al., 2018b; Kumar et al., 2018; Meier et al., 2023), with actin generally considered to anchor and stabilize stromule tips at the nucleus rather than to drive their formation. Time-lapse studies have independently shown stromule extension along microtubules, anchoring of stromules to perinuclear actin, and chloroplast movement toward a stromule tip anchored to actin near the nucleus. However, these independent observations have not yet been captured together in a single continuous recording. In particular, the proposed first step, a stromule tip growing along a microtubule from a plastid toward a distant nucleus and then anchoring to nuclear actin has not yet been directly documented. Consequently, the claim that plastids reach toward the nucleus via stromules is, to date, an interpretive synthesis of separate observations rather than a directly visualized sequence. This raises the possibility that these perinuclear actin-anchored stromule tips arise not from stromules growing toward the nucleus, but from the nucleus moving away from the plastid, as proposed in our model for *A. thaliana*. Crucially, the two proposed scenarios would be virtually indistinguishable in static images or during short time-lapse movies. Only by tracking nuclear dynamics over hours, the timescale required to capture the full oscillatory behavior of the nucleus as demonstrated here for *A. thaliana*, could one determine whether the nucleus in *N. benthamiana* is sought out by stromules or whether it actively drives stromule formation. The framework established in this study provides the experimental foundation needed to rigorously test this question in *N. benthamiana* and by extension, to determine whether the current model of kinesin-directed plastid recruitment during ETI responses is complete.

### XI-1, XI-2, and XI-K contribute to nucleus movement

Although nucleus movement was not the primary focus of this study, our analysis of myosin mutants revealed unexpected complexity in the myosin machinery underlying nuclear mobility in the upper epidermis. Historically, nucleus movement in plants has been primarily associated with myosin XI-I, which directly interacts with the nuclear envelope through the WIT1/WIT2 adaptor complex (Tamura et al., 2013). However, our study reveals that XI-I alone is not sufficient to account for the full extent of nuclear movement in the upper epidermis. Knockout of XI-I alone did not completely halt nuclear movement, whereas the *4ko* mutant (lacking XI-I, XI-1, XI-2, and XI-K) showed near-complete nuclear immobilization. Consistently, the 3ko mutant lacking XI-1, XI-2, and XI-K also displayed a significant reduction in nuclear movement, demonstrating that at least one or more of these three myosins contributes to nuclear mobility independently of XI-I. Since we did not assess the individual or pairwise contributions of XI-1, XI-2, and XI-K, we cannot determine which of these myosins is responsible or whether their contributions are redundant.

Since XI-K, XI-1, and XI-2 are not known to interact with the nuclear envelope directly, their influence on nuclear movement is likely indirect, mediated through effects on perinuclear actin organization and dynamics. Previous work showed that a xi-k/xi-2 double mutant exhibited disrupted actin organization, with randomly distributed actin bundles replacing the thick, longitudinally oriented filament arrays that normally guide rapid organelle dynamics (Ueda et al., 2010). Disorganization of actin tracks is expected to impair the directed movement of any cargo, including the nucleus, that depends on them, even without a direct myosin-nuclear envelope interaction. In our wildtype actin marker lines, we observed comparable thick actin bundles surrounding nuclei, with nuclei appearing to move along these structures (Video 19), suggesting that XI-K, XI-1, and XI-2 activity along these filament arrays may displace the nucleus independently of the direct XI-I/WIT1/WIT2 tethering mechanism. However, we note that this proposed mechanism remains speculative, as we did not directly image actin organization in the *3ko, 4ko* or *xi-i* mutant backgrounds.

### Genetic evidence for direct myosin-driven stromule formation

Beyond their contribution to nuclear movement, at least one or more of XI-K, XI-1, and XI-2 also drive a distinct subpopulation of stromule formation events near the nucleus that occur independently of nuclear displacement, pointing to a second actin-myosin mechanism of stromule formation. The notion that myosins might drive stromule formation through an actin-based mechanism was proposed in early work from the Gray and Hanson labs, where it was established that actin inhibitors reduce stromule formation and movement across multiple species and tissues (Gray et al., 2001; Kwok and Hanson, 2003; Kwok and Hanson, 2004). Natesan et al. (2009) subsequently provided more direct evidence by showing that RNAi-mediated knockdown of myosin XI in *N. benthamiana* reduced stromule formation, and that expression of the C-terminal cargo-binding domain of the same myosin localized to the chloroplast envelope and abolished stromules entirely, consistent with a dominant-negative effect. Independently, Sattarzadeh et al. (2009) showed that a specific sub-domain of the myosin XI-F tail localized to the chloroplast surface and along stromules, implicating a further myosin isoform in direct plastid–myosin interaction. However, the relationship between myosin localization and functional importance is not straightforward: XI-F was not among the myosins whose loss affected SPZ stromule formation in our genetic screen, nor was its removal required to abolish SPZ stromules in the *4ko* background. This illustrates that physical association with the plastid envelope is not sufficient to predict functional contribution and that a systematic dissection of individual myosin roles across isoforms, tissues, and cellular contexts remains an important goal for the field. Together, these earlier studies established a conceptual framework implicating myosin–actin interactions in stromule formation although the specific myosins responsible or the mechanism by which they acted could not be identified until now.

This nucleus-movement-independent subpopulation, representing approximately 25% of perinuclear stromule formation events, was unaffected by loss of XI-I or WIT1/WIT2, but was abolished in the *4ko* mutant, genetically identifying XI-K, XI-1, and/or XI-2 as drivers of this mechanism. That this subpopulation escaped detection in our primary mutant screen is likely explained by two non-mutually exclusive factors: nucleus-movement-independent events represent only approximately 25% of total perinuclear stromule formation, making them easy to overlook in single time-point analyses; and cooperative or redundant activity among XI-K, XI-1, and XI-2 means the effect might only become apparent when two or all three are removed simultaneously, as in the transition from the *xi-i* to the *4ko* mutant. Thus, while our data establish that one or more of XI-K, XI-1, and XI-2 are required for nucleus-movement-independent stromule formation, they do not resolve which myosin is individually responsible or whether they act cooperatively, a question that will require systematic analysis of single and double mutant combinations in the future. An additional layer of complexity is introduced by recent evidence that membrane contact sites between plastids and the ER contribute to plastid positioning and stromule dynamics near the nucleus (Kunjumon et al., 2024), raising the question of whether such ER-plastid interactions act in concert with, or independently of, the actin-based anchoring mechanisms described here.

### What is the biological purpose of nucleus driven stromule formation?

Yet underlying all the mechanistic questions addressed above is a more fundamental one: what biological purpose is served by this elaborate system of stromule mediated dynamic plastid-nucleus association, and why does it depend on a continuously oscillating nucleus rather than a stationary one? Our observations, while not designed to directly test function, nevertheless provide a coherent mechanistic basis for a functional model of this interaction.

Plastid clustering near the nucleus has long been proposed to facilitate inter-organellar communication, yet the continuous oscillatory movement of the nucleus poses a fundamental challenge to maintaining stable proximity with plastids in leaf epidermal pavement cells. Based on our observations, we propose a tethering model (Figure 9) in which a subset of plastids is associated with actin filaments encircling the nucleus while simultaneously anchored to peripheral actin filaments that constrain their position within the cell. This dual anchorage means that as the nucleus moves, it pulls its associated actin anchor points away from the stationary plastids, generating mechanical force that extrudes stromules from the plastid envelope at the attachment sites. Stromules formed in this manner may function like an extendable tether: the stromule tip remains attached to the moving nucleus while the plastid body stays stationary, and the finite capacity of the plastid membrane to sustain tubular extension may impose a physical constraint on nuclear displacement, potentially contributing to the oscillatory return behavior, a possibility that remains to be directly tested, but is evidenced by stromule thinning during stromule extension (Erickson et al., 2018a). In this way, *cor+* stromules, those forming in synchrony with nuclear displacement, maintain plastid-nucleus proximity despite continuous nuclear movement, a function supported by observations in which plastids that lose their stromule-nucleus connection are left behind as the nucleus moves on (Video 1).

**Figure 9:**
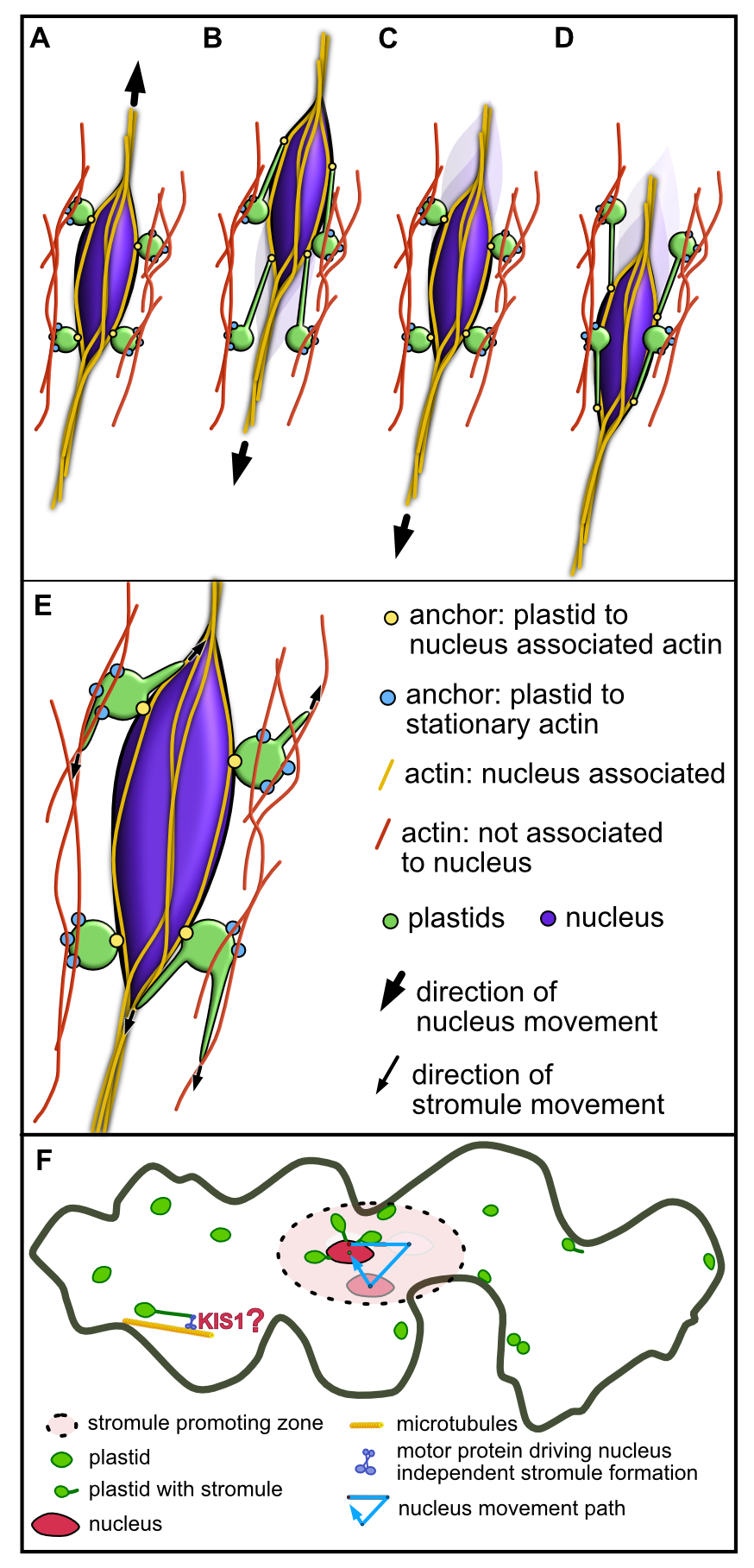
Model for the two distinct myosin dependent stromule formation mechanisms acting in the stromule promoting zone of upper epidermis cells in *A. thaliana*. A-D) Model for nucleus movement myosin-driven stromule formation: The nucleus (purple) is suspended within a cage of actin filaments (yellow) and undergoes myosin-driven oscillatory movement, primarily mediated by myosin XI-I with contributions from one or more of myosins XI-K, XI-1, and XI-2 (myosins not depicted). A subpopulation of plastids (green) positioned near the nucleus is simultaneously anchored to both the stationary cortical actin (red lines) cytoskeleton (blue dots) and to the mobile nucleus-associated actin cage (yellow dots). This dual anchoring system, combined with nucleus movement, generates mechanical force that pulls on the plastid envelope membrane, resulting in stromule formation, when the nucleus is moving (A-D). E) Model for nucleus movement independent myosin driven stromule formation: Stromules (protrusions extending from the green plastids) are generated through interactions with nucleus-associated actin (yellow) or actin filaments located in close proximity to the nucleus (red), likely mediated directly or indirectly by one or more of the myosins XI-1, XI-2, and XI-K (not depicted). The resulting increase in plastid surface area-to-volume ratio is proposed to enhance exchange between plastids and the immediate perinuclear environment. F) Both mechanisms described above operate within the stromule-promoting zone (pink shaded area) but cannot explain stromule formation outside this region. Therefore, at least one additional stromule formation mechanism likely exists outside the stromule-promoting zone. This mechanism remains to be identified but is proposed to involve kinesin motors (e.g., AtKIS1) and/or myosins acting independently of nuclear movement.

However, not all perinuclear stromule formation is synchronised with nuclear movement. The distinct spatial behaviour of the remaining cor− stromules is consistent with a different functional logic: rather than maintaining proximity during movement, we propose that these stromules may serve to amplify the contact surface between the plastid and the nucleus or perinuclear cytoplasm, maximizing the interface available for molecular exchange (Erickson et al., 2024). We therefore propose that SPZ stromules in the upper epidermis of *A. thaliana* serve two functionally distinct but complementary roles that correspond to these two temporal classes of stromule formation. Cor+ stromules function as dynamic tethers, ensuring sustained plastid-nucleus proximity across the full amplitude of nuclear oscillation. We propose that cor− stromules may function as contact surface amplifiers, extending the plastid membrane into the nuclear periphery to maximise the interface available for signal or metabolite exchange during nuclear rest. Whether signals such as H2O2 travel from the plastid to the nucleus through the stromule lumen itself, or simply diffuse through the cytoplasm across the short distance maintained by both tethering and surface contact, has not been experimentally distinguished (Caplan et al., 2015), and our model does not depend on resolving this question. In either case, maintaining proximity and maximising interactive surface are the structural prerequisites for efficient signal transfer, and the two stromule populations identified here provide a mechanistic basis for how both might be achieved in a cell with a continuously moving nucleus.

### Why is the nucleus constantly moving?

While nucleus positioning relative to the periclinal walls in leaf epidermis of *A. thaliana* cells is known to be responsive to blue light illumination, the oscillatory movement is distinct from directed photo relocation responses and persists independently of the phot2 blue light receptor (Higa et al., 2014). Interestingly, despite this detailed characterization of nucleus behavior in response to different light qualities, including light-dark positioning (Iwabuchi et al., 2010) and plastid-guided nucleus avoidance under strong light (Higa et al., 2014), the potential physiological consequences and benefits of oscillatory nuclear behavior for epidermis cells remain largely unexplored. The most concrete interpretation comes from studies of plastid-guided nuclear movement, which is hypothesized to serve as a protective measure against strong light and UV damage (Higa et al., 2014; Iwabuchi et al., 2010).

However, this interpretation primarily addresses directional short-term movement rather than the continuous oscillatory behavior we observe. The independence from blue light points toward a broader light-independent cellular origin. Indeed, similar oscillatory nucleus behavior can be found in root hair cells (Tamura et al., 2013) and other non-green tissues. In these cells, a general nuclear oscillation may serve to position the nucleus at the center of rather large mature cells and the pronounced mobility might originate from ongoing actin remodeling necessary for cytoplasmic streaming maintenance.

## Conclusion

By taking a systematic genetic approach in unstressed *A. thaliana* upper epidermis, this study reveals that the prevailing kinesin-microtubule model of stromule formation is not supported in this tissue and context. Instead, myosin XI-I drives nucleus oscillation within an actin cage, mechanically pulling stromules from anchored, stationary plastids and providing a mechanistic explanation for both the enrichment of stromules near the nucleus and the precise extent of the SPZ. Yet nucleus-driven extension is not the whole story: a distinct subpopulation of approximately 25% of perinuclear stromule formation events forms independently of nuclear movement, abolished only when XI-K, XI-1, and/or XI-2 are additionally removed, pointing to a direct actin-myosin mechanism operating in parallel. Strikingly, these two mechanistically distinct populations appear to serve two complementary biological functions: movement-coupled stromules act as extendable tethers maintaining plastid-nucleus proximity during nuclear oscillation, while movement-independent stromules are proposed to amplify the interactive interface between the plastid and the perinuclear region. Together they likely ensure that plastid-nuclear association is both maintained and optimized during nucleus oscillation. Future work will have to test if limited stromule length indeed constrains nucleus movement and thereby controls nucleus oscillation movement.

More broadly, this work challenges the field to treat stromule biology as context-dependent rather than governed by a single conserved mechanism, and invites a fresh look at *N. benthamiana*, where the role of nuclear movement in generating perinuclear stromules has never been examined and where the prevailing kinesin model may tell only part of the story.

## Methods

### Plant Material

*A. thaliana* myosin XI mutant lines used in this study are detailed in Supplemental Information 1 (provided by Valerian V. Dolja). The *wit1_wit2* double mutant was obtained from Zhao and co-workers (Zhao et al., 2008).

For cytoskeleton visualization, we utilized transgenic *A. thaliana* lines expressing p35S::GFP::MAP4 for microtubule labeling (Marc et al., 1998) and p35S::eGFP::ABD2::eGFP for actin filament visualization (Wang et al., 2008).

### Plant Growth conditions

*A. thaliana* plants cultivated in soil were grown in controlled environment chambers with short day conditions consisting of 8 hours light and 16 hours dark. Growth chambers were maintained at 21°C with a cool down to 17°C during the night and 60% relative humidity. Illumination was provided by T8 fluorescent bulbs with a 50:50 ratio of warm and cool light bulbs, delivering a light intensity of 100 - 120 μE m−2s−1 at soil level. For flower induction and seed production, plants were transferred to long day conditions (16 hours light/8 hours dark) with daytime temperatures of 24°C while maintaining all other growth parameters.

For growth in sterile conditions *A. thaliana* was sewn on AT-medium containing 5 mM KNO3, 2.5 mM KH2PO4, 2 mM MgSO4, 2 mM Ca(NO3)2, and 50 μM Fe(EDTA) as macronutrients. Micronutrients are supplemented at final concentrations of 70 μM H3BO3, 14 μM MnCl2, 0.5 μM CuSO4, 1 μM ZnSO4, 0.2 μM NaMoO4, 10 μM NaCl, and 0.01 μM CoCl2. The pH of the medium was adjusted to 5.8 with KOH prior to autoclaving. For solid medium, 0.7% (w/v) agar was added before autoclaving. Plates were incubated at 80 μE m−2s−1 at constant 21°C at 12h/12h day night cycle.

### Bacterial growth

Agrobacterium tumefaciens strain GV3101(pMP90), used for A. thaliana transformation, was grown in YEB medium containing sucrose supplemented with rifampicin (50 μg/mL) and gentamicin (20 μg/mL) plus the respective antibiotic needed for the T-DNA vector at 28°C. For electroporation, cells were made competent using a multi-step washing procedure with ice-cold sterile water and 10% glycerol. Transformation was performed using a Gene Pulser apparatus (Bio-Rad) set to 1.8 kV in 1 mm electroporation cuvettes. Following electroporation, cells were immediately recovered in 1 mL YEB medium and incubated with shaking (200 rpm) at 28°C for 1-2 hours. Transformed cells were selected on YEB plates containing appropriate antibiotics based on the transformed plasmid resistance in addition to rifampicin and gentamicin. E. coli strain Top 10 was used for cloning purposes and was grown in LB liquid or on solid medium supplemented with spectinomycin (50 µg/ml), carbenicillin (100 µg/ml) or kanamycin (50 µg/ml) depending on the vector. Chemically competent cells were transformed with a standard heat shock protocol.

### Plant Transformation and Selection

Transgenes where introduced into plant lines by using the direct floral dip method as described by (Davis et al., 2009). Transgenic plants were selected on AT-medium plates supplemented with 40 mg/L hygromycin for plsu4pn (plastid nucleus marker; (Erickson et al., 2017)) and pGreenII0129::So1-55aaFNR::eGFP (single plastid marker; (Schattat et al., 2011)). The nucleus-plastid-actin triple marker transgenics were selected on AT-medium containing 100 µg/ml kanamycin. All selection media also contained 100 µg/ml cefotaxime to suppress *A. tumefaciens* growth.

### Cloning

For visualizing plastids and nuclei, two established fluorescent protein constructs were used plsu4pn (Erickson et al., 2017) and pGreenII0129:FNReGFP (Schattat et al., 2011) (for more details see also Supplemental Methods 3). For this study a triple marker was cloned simultaneously, visualizing actin, plastids, and nuclei. For this the MoClo cloning system was used as described in (Werner et al., 2012; Engler and Marillonnet, 2014). For details on the composition see Supplemental Methods 3.

### Inhibitor treatments and DNA staining

Treatments with carbonyl cyanide m-chlorophenyl hydrazone (CCCP; Sigma-Aldrich C2759, Sigma-Aldrich Chemie GmbH, Taufkirchen, Germany) were used to investigate the 3D organization of actin at the nucleus. Leaves were treated with a 200 μM solution of CCCP (diluted in water) and directly prepared for imaging. This led to an immediate and reversible arrest of all active actin movement.

To investigate the role of actin and microtubules in plastid-nucleus interactions, leaves were treated with the actin-depolymerizing drug Cytochalasin D (CTD; Sigma-Aldrich C8273, 1 mg, Sigma-Aldrich Chemie GmbH, Taufkirchen, Germany) or the microtubule-depolymerizing drug oryzalin (Duchefa O1318, Duchefa Biochemie B.V., Haarlem, The Netherlands). CTD was prepared as a 50 mM stock in DMSO and diluted to a 4 μM working concentration (final DMSO concentration = 0.1%); oryzalin was prepared as a 50 mM stock in DMSO and diluted in tap water to a 100 μM working concentration (final DMSO concentration = 0.2%). Matched control solutions contained the corresponding DMSO concentration alone (0.1% for CTD controls, 0.2% for oryzalin controls).

Both inhibitors were applied using two complementary imaging strategies. For quantitative assessment of stromule frequency, leaf discs were incubated in treatment or control solution for 2 h, after which single time-point z-stacks were captured. For analysis of dynamics, leaf discs were vacuum-infiltrated with treatment or control solution and mounted in a custom silicone imaging chamber on a glass slide, covered with a coverslip, and imaged continuously to follow nucleus-plastid-stromule dynamics alongside actin or microtubule stability, respectively.

For labeling nuclei in plants without fluorescent nucleus marker genes, DAPI (4’,6-Diamidino-2-Phenylindole-Dihydrochloride; Sigma-Aldrich D9542, Sigma-Aldrich Chemie GmbH, Taufkirchen, Germany) was used. A 100 μM DAPI stock (in water) was diluted with tap water to a 6 μM working concentration, and tissue was incubated in solution for 10 minutes. Samples were washed for 1 minute in tap water prior to imaging.

### Imaging and Image Analysis

All experiments were conducted in the upper-epidermis of fully expanded mature leaves of 6- to 10-week-old plants. In total three different microscope setups were employed one epifluorescence and two different confocal laser scanning microscopes (for details see Supplemental Methods 2).

Myosin Mutant Screening - For screening myosin mutants, we employed the epifluorescence setup with the 38HE filter cube. For each leaf, 3 z-stacks were recorded. Images underwent extended depth of field processing (Schattat and Klosgen, 2009). Stromule counting was performed using MTB cell counter (Franke et al., 2015).

Cell and Nucleus Size Correlation - For cell and nucleus size correlation studies, the epifluorescence microscope setup with 43HE filter and standard brightfield using the 20x lens was employed. Both channels underwent extended depth of field processing (Schattat and Klosgen, 2009). Cell boundaries were labeled manually using LSM Image Browser software (Zeiss) in extended depth of field images, and the resulting outlines were analyzed together with nuclear sizes using Fiji (Schindelin et al., 2012) via thresholding and analyze particle functions. Cells not fully in the field of view were excluded from analysis.

Nuclear Speed and Shape Analysis - The Zeiss LSM 780 was used to observe nucleus shape and position. Nuclear movement tracking and speed measurements in z-stack projections were performed using Imaris 9.6.1, while nucleus shape and size were measured in the first frame of these z-stacks using Fiji (Schindelin et al., 2012). For time-lapse imaging, specimens were vacuum-infiltrated and mounted in a custom chamber created using silicon grease applied in a c-shape pattern on a glass slide. Leaf sections were mounted in tap water in the center of the silicon barrier and covered with a high-precision coverslip (No. 1.5H, 170 ± 5 μm). Excess water was removed using paper towel to enable gas exchange while maintaining hydration. Time-lapse sequences were acquired at 1,024 × 1,024 pixel resolution with 1.58 μs pixel dwell time, line averaging of 2, and 3-minute intervals between frames. Individual movies typically spanned 1.5-2 hours. To minimize photodamage while ensuring adequate imaging quality, these parameters were optimized in preliminary experiments, with chlorophyll autofluorescence monitored throughout imaging as an indicator of tissue health.

Stromule Frequency and Promoting Zone Measurements - These measurements were conducted using the epifluorescence microscope equipped with mCherry and GFP filter cubes and a 40x lens to record z-stacks of upper epidermis in eGFP, mCherry fluorescence, and transmitted light. Following extended depth of field calculations (Schattat and Klosgen, 2009), cell boundaries were manually labeled using LSM Image Browser software (Zeiss) in the bright field images. We employed an extended version of the MTB Cell Counter with expanded functionality to simultaneously detect both nuclei and plastids (Figure 3). Stromule and non-stromule plastids were subsequently categorized manually. Quantitative analysis of cell, nuclei, and plastid sizes and distances for stromule promoting zone analysis was performed using the new MiToBo operator “PlastidNucleiDistanceAnalyzer2D” as part of the MiToBo Fiji plugin (Möller et al., 2016). This operator assigns all plastids and nuclei to their respective cells based on center of mass positions, then measures the distance of each plastid to the nucleus by determining the shortest distance between any two contour pixels of the objects. Results include the number of plastids, their individual distances from the nucleus surface, nuclear size, cell size, and the maximum possible distance from the nucleus surface to the cell wall. The extended version of MTB Cell Counter and the MiToBo operator are available through MiToBo’s update site in Fiji (https://imagej.net/list-of-update-sites/). Data analysis was performed using standard R packages.

Cytoskeleton Imaging - For imaging nucleus-cytoskeleton-plastid constellations, the Leica Stellaris 8 setup was used with multitrack scanning of DAPI and GFP fluorescence. For actin imaging, cells were mounted in CCCP-containing water.

Actin Depolymerization Imaging - Imaging of *A. thaliana* triple marker (plastids/actin/nuclei) transgenic lines followed the same specimen preparation procedure described for nucleus speed measurements. The Leica Stellaris 8 setup was used for imaging, with stacks captured approximately 4 minutes and 17 seconds apart during time-lapse experiments.

## Supplemental Data

The following Supplemental Data are provided:

## Supplemental Information (single PDF file)

Supplemental Methods 1 — SPZ measurements: image analysis pipeline description

Supplemental Methods 2 — Microscopy imaging setups

Supplemental Methods 3 — Plasmid and construct details

Supplemental Information 1 — Lines used for myosin mutant screen

Supplemental Information 2 — Statistical analysis of SF in myosin XI mutants

Supplemental Information 3 — Transgenic lines used for nucleus roundness analysis

Supplemental Information 4 — Nucleus roundness in mutants: figures

Supplemental Information 5 — Nucleus roundness in mutants: statistical tests

Supplemental Information 6 — Time-lapse movies used for nucleus velocity data

Supplemental Information 7 — Correlation of cell and nucleus size in genotypes

Supplemental Information 8 — Statistical analysis of nucleus speed in pn marker lines

Supplemental Information 9 — Statistical analysis of SF in plastid-nucleus marker lines

Supplemental Information 10 — Transgenic lines used for SPZ measurements

Supplemental Information 11 — Plastid distribution relative to the nucleus: figures

Supplemental Information 12 — Plastid distribution relative to the nucleus: stats

Supplemental Information 13 — SPZ measurements: SF by distance and genotype

Supplemental Information 14 — SPZ measurements: stats

Supplemental Information 15 — Nucleus speed vs. SF correlation analysis: data

Supplemental Information 16 — Nucleus speed vs. SF correlation analysis: graphs

Supplemental Information 17 — Stromule formation events in time-lapse recordings

Supplemental Information 18 — Nucleus tracks with calculated oscillation center points

Supplemental Information 19 — Distribution of nucleus distances to track center point

Supplemental Information 20 — Nucleus distance distributions: figures with tracks

Supplemental Information 21 — Actin filaments at stromule-nucleus contact sites Fig1

Supplemental Information 22 — Actin filaments at stromule-nucleus contact sites Fig2

Supplemental Information 23 — Microtubule proximity to stromule tips at the nucleus

Supplemental Information 24 — Impact of oryzalin on SF in the SPZ

Supplemental Information 25 — Impact of cytochalasin D on SF in the SPZ

## Video files (22 files, Video 01–22)

Video 1 — Time-lapse of nuclear oscillation and stromule dynamics in wild-type Video 2 — Stromule formation coincides with nuclear movement in wild-type Video 3 — Comparison of nuclear movement and stromule dynamics: WT vs. *xi-i*

Video 4 — Comparison of nuclear movement and stromule dynamics: WT vs. *wit1/2* Video 5 — Comparison of nuclear movement and stromule dynamics: WT vs. *4ko* Video 6 — Three-dimensional reconstruction (DAPI/microtubules) example 1

Video 7 — Three-dimensional reconstruction (DAPI/microtubules) example 2 Video 8 — Three-dimensional reconstruction (DAPI/microtubules) rare instance Video 9 — Three-dimensional reconstruction (DAPI/actin) example 1

Video 10 — Three-dimensional reconstruction (DAPI/actin) example 2 Video 11 — Three-dimensional reconstruction (DAPI/actin) example 3 Video 12 — Triple-marker reconstruction (actin/plastids/nucleus): example 1 Video 13 — Triple-marker reconstruction (actin/plastids/nucleus): example 2 Video 14 — Triple-marker reconstruction (actin/plastids/nucleus): example 3 Video 15 — Triple-marker reconstruction (actin/plastids/nucleus): example 4 Video 16 — Triple-marker reconstruction (actin/plastids/nucleus): example 5

Video 17 — Triple-marker reconstruction (microtubules/plastids/nucleus): example 1 Video 18 — Triple-marker reconstruction (microtubules/plastids/nucleus): example 2

Video 19 — Time-lapse: DMSO control (CTD)

Video 20 — Time-lapse: impact of cytochalasin D on actin, nucleus and stromules Video 21 — Time-lapse: DMSO control (Oryzalin)

Video 22 — Time-lapse: impact of oryzalin on microtubules, nucleus and stromules

## Accession Numbers

No new sequence data was generated for this manuscript.

## Author Contributions

JLE and MHS designed the research with MHS supervising the project. JLE performed the myosin mutant screen, genotyping, nucleus shape analysis, and plastid marker transformations; SÖ performed time-lapse microscopy, analysis, and generated triple marker lines; CG and VB performed and analyzed inhibitor treatments; JM contributed to time-lapse data collection and analysis; SO analyzed stromule forming events in time lapse experiments; AG and CA performed screening and stromule frequency analysis of transgenic lines; BM developed the MTB Cell Counter Plugin and MiToBo operator “PlastidNucleiDistanceAnalyzer2D”. MHS and SÖ conducted actin and cytoskeleton experimental work. MHS, JLE, SÖ, and JM developed microscopy protocols. JLE and MHS performed statistical analyses. JLE and MHS analyzed the data, prepared figures and videos, and wrote the paper.

## Acknowledgments

We thank Valerian V. Dolja (Oregon State University) for providing myosin mutant seeds and advice on initial study design and genotyping. We acknowledge Helena Schnuch for volunteering in preliminary work setting up the myosin mutant screen and Viktoria Burgdorf for help in analyzing images for stromule promoting zone measurements. We thank Bianca Rosinsky and Siegfried Platzer for technical assistance with horticulture. We thank Vivian Linke for supporting proof reading and consistency checks. We note that large language models (specifically Claude AI and ChatGPT) were used to perform standard editorial tasks such as help with grammar, spelling, and ensuring consistency in references to figures, videos, and supplemental material. No image and video content was created or data analysis was performed by these or other AI tools.

