## Supplementary Information for "“Myosin-Driven Nuclear Movement Generates Stromules in *Arabidopsis thaliana*”"

### 1 Supplemental Information

#### 1.1 Supplemental Methods

##### 1.1.1 Supplemental Methods 1 - Stromule promoting zone measurements

**Step 1: Detection of nuclei and plastids** Stromule promoting zone (SPZ) measurements were carried out using \*pn\* transgenic lines expressing FNRtp:eGFP (plastids) and H2B:mCherry (chromatin/nuclei). Imaging was performed on an epifluorescence microscope equipped with eGFP and mCherry filter cubes and a 40× objective, recording z-stacks of the upper epidermis in the eGFP, mCherry, and transmitted light (DIC) channels. All channels were exported as individual image files and sorted by channel type (DIC, eGFP, mCherry).

Cell outlines were generated by importing the DIC images into LSM Image Browser (Zeiss) and using the overlay function to produce red cell contour lines against a black background, which were then exported as separate images (see example in Figure 3A main manuscript). Extended depth-of-field images were subsequently calculated for both the eGFP (plastid) and mCherry (chromatin/nucleus) channels using the Combine Z approach described in Schattat and Klösgen (2009), yielding a single in-focus composite image per channel.

The resulting outline image and both extended depth-of-field images were combined into a stack and imported into the MTB Cell Counter plugin (Franke et al., 2015) in Fiji. Within this tool, plastids and nuclei were detected semi-automatically, and plastids bearing stromules were assigned a distinct counter type in a semi-manual step, enabling separate tallies of total plastid numbers and stromule-bearing plastids. Detection errors were corrected manually. The graphical output of this detection step, showing nuclei (red outline), plastids without stromules (yellow outline), and plastids with stromules (green outline), is illustrated in Figure 3B of the main manuscript. The detection output was then passed to the MiToBo runner and the operator "PlastidNucleiDistanceAnalyzer2D", which automatically assigns plastids and nuclei to their respective cells based on centre-of-mass positions and calculates the shortest distance between the contour pixels of each plastid and its associated nucleus, alongside cell size and the maximum possible nucleus-to-cell-wall distance. The overlay of the runner output on the stacked microscopic images for all genetic backgrounds used in this study, with the 8 µm zone indicated, is shown in Figure 3C–F, main manuscript.

The extended version of MTB Cell Counter and the MiToBo operator "PlastidNucleiDistanceAnalyzer2D" are publicly available through MiToBo's update site in Fiji as of version 2.4.1.

**Step 2: Stromule promoting zone analysis** The distance table produced by PlastidNucleiDistanceAnalyzer2D formed the basis for all SPZ analyses shown in Figure 4A–D, main manuscript. Using R and R-studio, cells were first sorted into size classes (small: <1000 µm<sup>2</sup>; medium: 1000–3000 µm<sup>2</sup>; large: >3000 µm<sup>2</sup>) as described in Erickson et al. (2014). Within each size class and genotype, plastids with and without stromules were binned into consecutive 4 µm distance zones from the nucleus surface, a bin width chosen to correspond to the average plastid body length. Stromule frequency was calculated for each zone as the proportion of plastids bearing a stromule relative to the total plastid count in that zone (Figure 4A–D, main manuscript). Because measurements were collected from at least three independent transgenic lines derived from different plants, stromule frequency values were averaged across biological replicates and the standard deviation was calculated to assess reproducibility.

Distance zones containing three or fewer plastids were excluded from the stromule frequency analysis. This was predominantly the case for zones at greater distances from the nucleus, well outside the stromule promoting zone, where fewer plastids are present. In these small samples, the discrete nature of stromule counts produces extreme variation in the calculated frequency ratio, for example, a single stromule difference in a zone of three plastids shifts the frequency by 33 percentage points, whereas the same difference in a zone of ten plastids shifts it by only 10 percentage points. As these zones fall outside the SPZ and their exclusion therefore does not affect the conclusions drawn from the analysis, this threshold was applied to avoid frequency values that reflect sampling artefacts of small denominators rather than genuine biological patterns.

The resulting data were used to generate the bar charts presented in Figure 4A–D.

##### 1.1.2 Supplemental Methods 2 - Imaging setups

**Epifluorescence System:** Zeiss ObserverZ1 fluorescence microscope equipped with an HXP 120V light source and AxioCamMRm monochrome camera. This setup included filter cubes 43HE (DsRed: excitation 550/25 nm, beam splitter 570 nm, emission 605/70 nm) and 38HE (GFP: excitation 470/40 nm, beam splitter 495 nm, emission 525/50 nm), along with two objectives: a 20x/0.5 ECPlan-Neofluoar Zeiss and a 40x/0.75 ECPlan-Neofluoar, controlled by ZenBlue software (Carl Zeiss).

**Confocal System I:** Zeiss LSM780 microscope featuring an AxioObserverZ1 inverted stage and a C-Apochromat 63x/1.20 water immersion objective, controlled by ZenBlack software (Carl Zeiss). Fluorescence was excited using a 488 nm laser line from a multiline argon laser (25 mW) at 1.2% power and a 561 nm diode laser at 0.5% power. Emission was collected in separate channels: eGFP (493-556 nm), mCherry (596-623 nm), and chlorophyll autofluorescence (658-758 nm).

**Confocal System II:** Leica Stellaris 8 confocal laser scanning microscope equipped with a SuperK Extreme white light laser and a HC PL APO CS2 40x/1.10 WATER objective. Settings included EGFP (excitation: 489 nm at 3.38% laser power, detection: 480-524 nm using HyD X 1 detector), mKOK (excitation: 548 nm at 2.00% laser power, detection: 553-593 nm using HyD S 2 detector), and E2-Crimson (excitation: 611 nm at 7.10% laser power, detection: 626-649 nm using HyD X 3 detector). Image acquisition was controlled using LAS X software version 4.2.1.23810 (Leica Microsystems). Lightning processing was applied with auto setup at different iterations (Figure 8 A-V, Supplementary Information 21-23, and Videos 6-18).

##### 1.1.3 Supplemental Methods 3 - Plasmids

**Plastid-Nucleus and only Plastid marker** For visualizing plastids and nuclei, two established fluorescent protein constructs were used. The *plsu4pn* construct combines p35S-FNRtp-eGFP-t35S for plastid stroma labeling with pUBQ10-H2B-mcherry-tNOS for nuclear visualization in the hygromycin-resistant *plsu4* vector backbone (Erickson et al., 2017). The pGreenII0129-FNR-eGFP construct (Schattat et al., 2011) was created by cloning the FNR-EGFP fusion between BamHI and XbaI sites in a modified pRT100 expression cassette, followed by transferring the complete cassette into pGreenII0129 using PstI restriction sites.

**Plastid-Actin-Nucleus marker** The construct is based on the L2 vector pAGM4723 and contains four L1 expression cassettes: 1) *AtuNos-npt-AtuNos* for plant selection (Engler et al., 2014), pAtUBQ10-eGFP-ABD2-eGFP-tNOS for actin visualization (Position 2), pUBQ10-NLS40-mKOK-GUS-tOCS for nuclear labeling (Position 3), and pOCS-SSU-E2Crimson-tOCS for plastid targeting (Position 4). Assembly was performed by first using L0 modules of promotor, coding sequence and terminator modules using BsaI-based Golden Gate cloning. Next the L1 modules were assembled in the L2 by using BpiI-based Golden Gate cloning, combining individual Level 1 constructs into the final Level 2 vector. For position 2 a L1 Module with Gateway recombination sites was used to create an intermediate L2 vector (P1\_KanR--P2\_GW--P3\_NLSmKOKGUS--P4\_ssuE2-Crimson). The pAtUBQ10-eGFP-ABD2-eGFP-tNOS was sourced from (Wang et al., 2008). The whole expression cassette was PCR amplified with attB sites, subcloned into a pDonor vector and then inserted into the intermediate L2 vector via a Gateway LR recombination reaction.

##### 1.1.4 Calculating cell size nucleus size correlation

**Image acquisition** The upper epidermis of *A. thaliana* consists of guard cells, trichomes and pavement cells. Among those pavement cells vary significantly in size. Because cell size might impact nucleus movement and the stromule promoting zone, cell size was taken into consideration during our analysis. A limited recording area during time-lapse microscopy of the epidermal layer is the reason why not every cell size can be determined from cell outlines. As described in (Melaragno et al., 1993) the size of the epidermal cell is related to the size of the nucleus in *A. thaliana*. When epidermal cells increase in size the respective nucleus also gets larger. Therefore, the size of a nucleus can be used as an indirect measure for cell size. Because this correlation is known for wildtype, but not the here used mutant lines (*wit1\_wit2*, *xi-i*, *3ko* and *4ko*) this correlation had to be measured for these lines too. For this, z-stacks from the upper epidermis of pn transgenic plants were taken with a 20x objective by an epifluorescence microscope (AxioObserver Z1, Zeiss, Jena, Germany) in transmitted light and in a mcherry fluorescence channel, capturing with the later the H2B:mCherry fluorescence, which localizes to the nucleus.

**Data processing** In ZEN LE Software (Zeiss, 2007), z-stacks were isolated and exported for further analysis. Using CombineZP, the image stacks were aligned to enhance nuclei brightness and improve the visibility of cell borders, facilitating more efficient outlining. The exported image stacks, now highlighting cell borders, were then processed in LSM Image Browser Version 3.5 (Carl Zeiss GmbH, Jena). This stacking allowed for variation between cell layers, enhancing precision. Subsequently, the outlined images were exported to Fiji v.2.0.0-rc-43 (Schindelin et al., 2012), where the contrast was inverted to black. To exclude stomata, these regions were filled with the outline color. The images were then converted to 8-bit grayscale. Using the “threshold” function, Fiji measured the sizes of the outlined cells with the “analyze particles” tool, which also assigned numbers to the outlines corresponding to their nuclei. For accurate scaling, the images were calibrated using a defined scale bar and the “set scale” function, determining the actual size of the cells. For nuclei size measurements, the image stacks were also exported to Fiji and overlaid on the outlined cell images. Due to strong fluorescence causing reflection by the cell walls, Fiji could not automatically detect the nuclei borders, necessitating manual measurements.

#### 1.2 Supplemental Information

##### 1.2.1 Supplemental Information 1 - Stromule Frequency in Myosin Mutants - Lines

**Supported Figure: 1A** This table contains supplemental information for data shown in Figure 1A. For each of the analyzed genetic backgrounds at least three independent transgenic lines (FNR-eGFP) were analyzed. Because wildtype was always grown along with each mutant batch to control for growth conditions a larger number (n) of wildtype data was available for the analysis.

| Genotype | Locus | Insertion line | Plant lines | Total lines | Total plants | Images analyzed |
| --- | --- | --- | --- | --- | --- | --- |
| WT - Col 0 | - | - | 3, 6, 8 | 3 | 48 | 144 |
| xi-a | At1g04600 | GABI_622E02<sup>1</sup> | 4, 7, 9 | 3 | 9 | 27 |
| xi-b | At1g04160 | SALK_113062<sup>1</sup> | 8, 14, 15 | 3 | 9 | 27 |
| xi-c | At1g08730 | GABI_262B03<sup>1</sup> | 1, 2, 3 | 3 | 9 | 27 |
| xi-d | At2g33240 | SALK_082078<sup>1</sup> | 4, 5, 6 | 3 | 9 | 27 |
| xi-e | At1g54560 | SALK_072023<sup>1</sup> | 5, 9, 11 | 3 | 9 | 27 |
| xi-f | At2g31900 | GABI_070F03<sup>2</sup> | 1, 2, 4 | 3 | 9 | 27 |
| xi-g | At2g20290 | SALK_018032<sup>1</sup> | 3, 5, 9 | 3 | 9 | 27 |
| xi-h | At4g28710 | SAIL_365_D03<sup>1</sup> | 7, 4, 2 | 3 | 9 | 27 |
| xi-i | At4g33200 | SALK_082443<sup>1</sup> | 2, 3, 5, 9 | 4 | 17 | 51 |
| xi-j | At3g58160 | SALK_063159<sup>1</sup> | 2, 3, 4 | 3 | 9 | 27 |
| xi-k | At5g20490 | SALK_067972<sup>1</sup> | 1, 2, 7 | 3 | 9 | 27 |
| xi-1 | At1g17580 | SALK_019031<sup>1</sup> | 1, 5, 9 | 3 | 9 | 27 |
| xi-2 | At5g43900 | SALK_055785<sup>1</sup> | 2, 3, 4 | 3 | 9 | 27 |
| 3KO<sup>3</sup> xi-1, xi-2,<br>xi-k | At1g17580,<br>At5g43900,<br>At5g20490 | SALK_055785,<br>SALK_019031,<br>SALK_067972 | 2, 3, 7 | 3 | 18 | 54 |
| 4KO<sup>3</sup> xi-1, xi-2,<br>xi-k, xi-i | At1g17580,<br>At5g43900,<br>At5g20490,<br>At4g33200 | SALK_055785,<br>SALK_019031,<br>SALK_067972,<br>SALK_082443 | 1, 2, 3 | 3 | 9 | 27 |

**Table 1.1:** Myosin mutant lines analyzed for stromule quantification in *A. thaliana* (ecotype Columbia-0) wild type (WT) and T-DNA insertion mutants in all 13 myosin class XI genes as well as a triple and a quadruple mutant were stably transformed with FNRtp:eGFP to visualize plastids and stromules. Stromule frequency counts were collected from multiple plants originating from at least three different primary transformants (plant lines chosen are listed in column 4). Plant sample sizes used for data displayed in Figure 1A as well as total images analysed are shown in columns 6 and 7, respectively. Seeds were provided by: The Arabidopsis Information Resource (TAIR), www.arabidopsis.org<sup>1</sup>, The Nottingham Arabidopsis Stock Centre (NASC), http://arabidopsis.info/<sup>2</sup> and Valerian Dolja, Oregon State University, Corvallis, Oregon, USA, published in Peremyslov et al. (2010)<sup>3</sup>.

##### 1.2.2 Supplemental Information 2 - Stromule Frequency in Myosin Mutants - Stats

**Supported Figure: 1A** Table 1.2 on the following page and Table 1.3 on page 11 contain supplemental information for data shown in Figure 1A. Because stromule frequency values in this part of the study are ratios, values were arcsin transformed prior to statistical analysis ( 1.2 on the following page). Using SigmaPlot 12.5 software a One Way Analysis of Variance was performed, because the Normality Test (Kolmogorov-Smirnov) Failed ($P < 0.050$), an ANOVA on Ranks analysis was started: Kruskal-Wallis One Way Analysis of Variance on Ranks ($H = 166.485$ with 15 degrees of freedom). ($P = <0.001$) The differences in the median values among the treatment groups are greater than would be expected by chance; there is a statistically significant difference ($P = <0.001$). In order to isolate the group or groups that differ from the others a multiple comparison procedure versus a control group (Dunn's Method) was used. For plotting, the data was back transformed ( 1.3 on page 11).

| ANOVA on Ranks |  |  |  |  |  |
| --- | --- | --- | --- | --- | --- |
| Group | N | Missing | Median | 25% | 75% |
| WT | 144 | 0 | 0.476 | 0.421 | 0.523 |
| myosin xi-a | 27 | 0 | 0.345 | 0.3 | 0.523 |
| myosin xi-b | 27 | 0 | 0.508 | 0.431 | 0.56 |
| myosin xi-c | 27 | 0 | 0.372 | 0.335 | 0.512 |
| myosin xi-d | 27 | 0 | 0.426 | 0.36 | 0.52 |
| myosin xi-e | 27 | 0 | 0.489 | 0.407 | 0.546 |
| myosin xi-f | 27 | 0 | 0.485 | 0.415 | 0.511 |
| myosin xi-g | 27 | 0 | 0.45 | 0.419 | 0.486 |
| myosin xi-h | 27 | 0 | 0.424 | 0.385 | 0.462 |
| myosin xi-i | 51 | 0 | 0.318 | 0.269 | 0.427 |
| myosin xi-j | 27 | 0 | 0.451 | 0.406 | 0.504 |
| myosin xi-k | 27 | 0 | 0.495 | 0.447 | 0.562 |
| myosin xi-1 | 27 | 0 | 0.466 | 0.419 | 0.585 |
| myosin xi-2 | 27 | 0 | 0.527 | 0.469 | 0.563 |
| 3KO | 54 | 0 | 0.436 | 0.34 | 0.531 |
| 4KO | 27 | 0 | 0.154 | 0.11 | 0.224 |
| Dunn's |  |  |  |  |  |
| Comparison | Diff of Ranks | Q | P | P<0.050 |  |
| 4KO vs WT | 323.233 | 8.891 | <0.001 | Yes |  |
| I vs WT | 205.069 | 7.26 | <0.001 | Yes |  |
| A vs WT | 112.622 | 3.098 | 0.029 | Yes |  |
| C vs WT | 90.751 | 2.496 | 0.188 | No |  |
| H vs WT | 77.381 | 2.129 | 0.499 | Do Not Test |  |
| 2 vs WT | 76.582 | 2.107 | 0.527 | Do Not Test |  |
| D vs WT | 59.677 | 1.642 | 1 | Do Not Test |  |
| 3KO vs WT | 58.668 | 2.121 | 0.509 | Do Not Test |  |
| J vs WT | 50.066 | 1.377 | 1 | Do Not Test |  |
| K vs WT | 46.582 | 1.281 | 1 | Do Not Test |  |
| B vs WT | 40.082 | 1.103 | 1 | Do Not Test |  |
| G vs WT | 33.807 | 0.93 | 1 | Do Not Test |  |
| 1 vs WT | 17.582 | 0.484 | 1 | Do Not Test |  |
| E vs WT | 14.582 | 0.401 | 1 | Do Not Test |  |
| F vs WT | 4.73 | 0.13 | 1 | Do Not Test |  |

Table 1.2: Details of the performed statistical test: N = sample size considered in each group (each sample corresponds to the arcsin transformed average stromule frequency of a single tested leaf, calculated from three image stacks taken for each leaf); Missing = missing values; Median = median value of n arcsin transformed stromule frequency values; 25% and 75% = respective percentiles defining the box); Comparison = compared groups; Diff of Ranks = difference of ranks; Q = the difference between the two means divided by the standard error of that difference; P = p-value of each pair; p>0.050 = significance statement by Sigmaplot 'Yes' states that the difference is larger than it would be expected by chance, 'No' states that the difference can be explained by chance, 'Do Not Test' stats that the previous results of paired tests does make the test obsolete and is therefore not performed.

|  | Mean | absolute value<br>for lower back-transformed<br>CI | absolute value<br>for Upper<br>Backtransformed<br>CI | difference to<br>Median of<br>Lower 95 CI | difference to<br>Median of<br>Upper 95 CI |
| --- | --- | --- | --- | --- | --- |
| WT | 0.207533955 | 0.196056665 | 0.219250455 | 0.011477291 | 0.0117165 |
| myosin xi-a | 0.155250784 | 0.121994817 | 0.19170801 | 0.033255966 | 0.036457227 |
| myosin xi-b | 0.229007917 | 0.199322344 | 0.260113345 | 0.029685573 | 0.031105428 |
| myosin xi-c | 0.170008427 | 0.134701345 | 0.208508095 | 0.035307082 | 0.038499668 |
| myosin xi-d | 0.194708584 | 0.15387713 | 0.239084345 | 0.040831453 | 0.044375762 |
| myosin xi-e | 0.213240007 | 0.189586835 | 0.237891101 | 0.023653172 | 0.024651094 |
| myosin xi-f | 0.207533955 | 0.187396659 | 0.228419911 | 0.020137297 | 0.020885955 |
| myosin xi-g | 0.192338236 | 0.172948429 | 0.2125032 | 0.019389807 | 0.020164964 |
| myosin xi-h | 0.177586726 | 0.156193521 | 0.200041914 | 0.021393205 | 0.022455188 |
| myosin xi-i | 0.102564083 | 0.083918669 | 0.122847022 | 0.018645414 | 0.020282939 |
| myosin xi-j | 0.185293987 | 0.16873316 | 0.202447488 | 0.016560827 | 0.017153501 |
| myosin xi-k | 0.230690858 | 0.204218388 | 0.258272942 | 0.02647247 | 0.027582084 |
| myosin xi-1 | 0.223985217 | 0.192969249 | 0.256611489 | 0.031015968 | 0.032626272 |
| myosin xi-2 | 0.238317024 | 0.216608296 | 0.260727754 | 0.021708729 | 0.02241073 |
| 3KO | 0.182195803 | 0.15402149 | 0.212176028 | 0.028174313 | 0.029980225 |
| 4KO | 0.027303817 | 0.018489864 | 0.037773916 | 0.008813953 | 0.010470099 |

**Table 1.3:** Statistic data for stromule frequency analysis of different FNReGFP transgenic myosin XI mutants and wt. The table presents the values after the backtransformation "Lower 95% CI" and "Upper 95% CI" represent the lower and upper bounds of the 95% confidence intervals, before back transformation. The "Mean" column provides the arithmetic mean of the data, before back transformation. The "Absolute value for lower back-transformed CI" and "Absolute value for upper back-transformed CI" columns display the absolute values of the lower and upper bounds of the back-transformed confidence intervals. The columns "Difference to median of lower 95% CI" and "Difference to median of upper 95% CI" show the differences between the confidence interval bounds and their respective median values of the back transformed values. Note that CI stands for confidence intervals.

##### 1.2.3 Supplemental information 3 - Nucleus Roundness in Mutants - Lines

**Supported Figure: 2A-E** This table contains supplemental information for data shown in Figure 2F. For each tested genetic background three or two of confocal z-stacks of different transgenic lines were analyzed. In STable 1.4 the used lines are summarized.

| Genotype | Locus | Insertion line, creating mutant phenotype | Plant lines created for project | Plant lines used to measure nucleus roundness | Number of Total plants used for measurements | Images analyzed |
| --- | --- | --- | --- | --- | --- | --- |
| WT - Col 0 | - | - | 2, 3, 10, 18 | WT-pn-line-2-1<br>WT-pn-line-3-1<br>WT-pn-line-10-1 | 3 | 3 |
| xi-i | At4g33200 | SALK_082443<sup>1</sup> | 6, 12, 17 | xi-i-pn-line-4-6<br>xi-i-pn-line-12-01 | 2 | 2 |
| wit1_wit2<sup>4</sup> | At5g11390<br>At1g68910 | GABI-470E06,<br>SALK_127765 | 8, 12, 13 | wit12 pn-line-08-01<br>wit12 pn-line-12-01 | 2 | 2 |
| 3KO<sup>3</sup> xi-1, xi-2,<br>xi-k | At1g17580<br>At5g43900<br>At5g20490 | SALK_055785<br>SALK_019031<br>SALK_067972 | 2, 3, 4 | 3KO pn-line-3-02<br>3KO pn-line-9-02 | 2 | 2 |
| 4KO<sup>3</sup> xi-1, xi-2,<br>xi-k, xi-i | At1g17580<br>At5g43900<br>At5g20490<br>At4g33200 | SALK_055785<br>SALK_019031<br>SALK_067972<br>SALK_082443 | 1, 2, 3 | 4KO pn-line-1-1-02<br>4KO pn-line-2-1-02<br>4KO pn-line-3-1-02 | 3 | 3 |

**Table 1.4:** Transgenic lines created for the analysis of nucleus shape and movement. Seeds were provided by: The Arabidopsis Information Resource (TAIR), www.arabidopsis.org<sup>1</sup>, The Nottingham Arabidopsis Stock Centre (NASC), http://arabidopsis.info/<sup>2</sup> and Valerian Dolja, Oregon State University, Corvallis, Oregon, USA, published in Peremyslov et al. (2010)<sup>3</sup>. Ikuko Hara-Nishimura, Department of Botany, Graduate School of Science, Kyoto University, Kyoto 606-8502, Japan, published in Tamura et al., 2013. Originally created and published in Zhao et al., 2008<sup>4</sup>.

###### 1.2.4 Supplemental information 4 - Nucleus Roundness in Mutants - Figure

**Supported Figure: 2A-E** Because in wildtype plants nucleus shape, in the upper epidermis, seems to change with increasing nucleus size (see main manuscript figure 2A) we plotted the circularity index for nuclei in the different genetic backgrounds against the respective nucleus size (in $\mu\text{m}^2$). The formula for circularity is $\text{circularity} = 4\pi\left(\frac{\text{area}}{\text{perimeter}^2}\right)$. The more round a particle the closer the circularity index value is to 1.0. The following figure (SFigure 1.1) captures this correlation. In order to compare the nucleus shape between different sized nuclei, nuclei were sorted into three size categories (small, medium and large). The separation into these three classes was done based on the established cell size classes described in Erickson et al., 2017. A correlation curve between nucleus size and cell size was used to define the boundaries for nucleus size used in this figure (small = $< 22.6 \mu\text{m}^2$; medium = $> 22.6 \mu\text{m}^2, < 41.8 \mu\text{m}^2$; large = $> 41.8$). This correlation curve is shown for wildtype in SFigure 1.2 panel A. Additional correlation curves and circularity measurements for the different mutants in comparison to wildtype are shown on SFigure B - F). To better test differences between different sized nuclei, size classes were formed (small, medium and large) based on cell size (see main manuscript and Erickson et al., 2017) in which all nuclei of respective cells were gathered. These were then compared across the various genetic backgrounds. The resulting boxplots are depicted in SFigure 1.1 panel E. In summary SFigure 1.1 shows that mutants with a mutant *xi-i* allele show more round nuclei, which becomes specifically evident in larger epidermis cells.

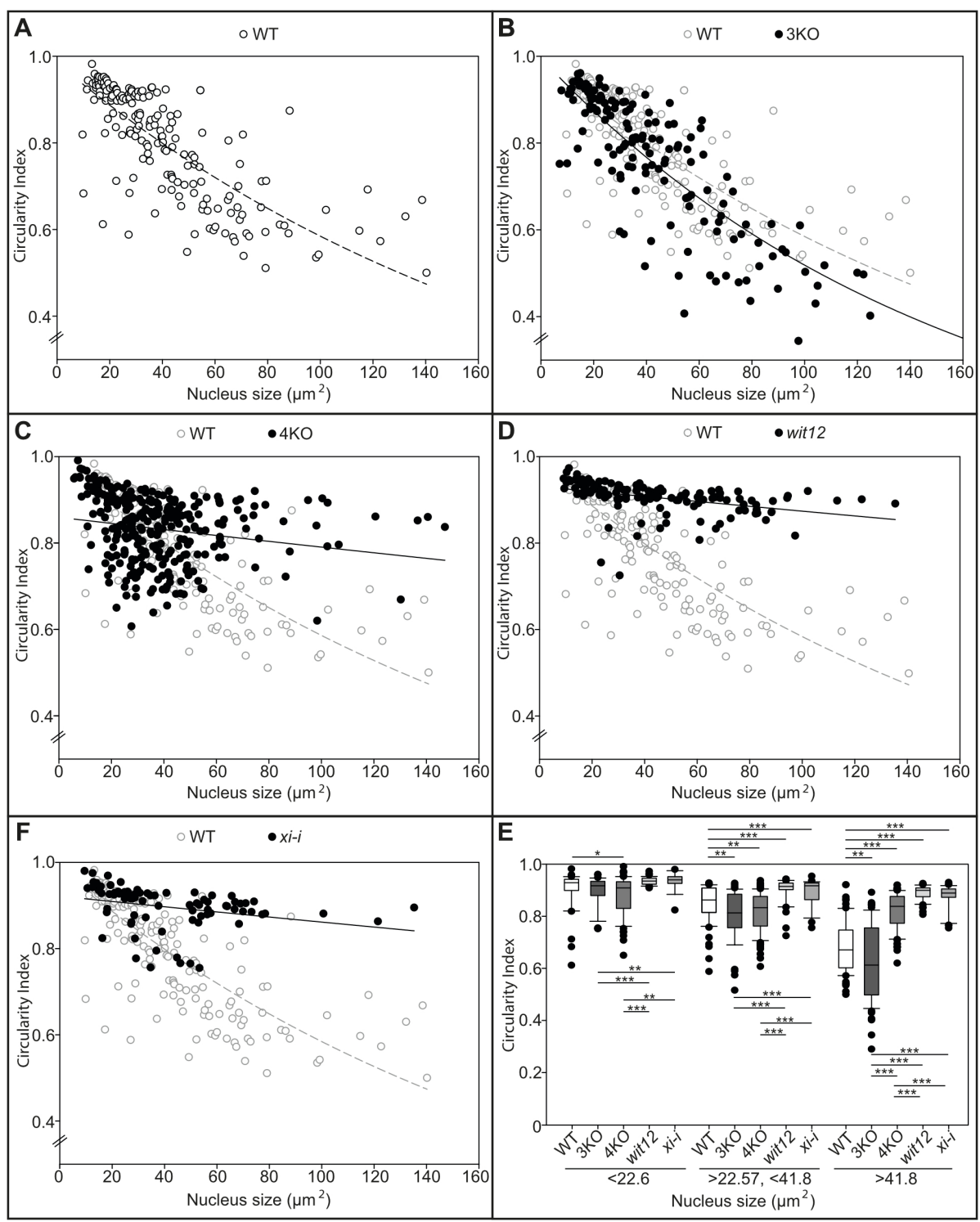

**Figure 1.1:** Correlation between nucleus roundness and nucleus size in WT and mutant plants. A) Correlation of WT nucleus size and roundness. A clear correlation between roundness and size is visible: the larger the nuclei, the less round they are. Hollow circles represent individual WT nuclei. B) - F) Correlations between nucleus roundness and nucleus size in the different mutant backgrounds compared to WT. Mutant nuclei are represented by black dots, while WT nuclei are shown as hollow circles. It becomes evident, that except for the 3ko mutant nuclei stay more round with increasing size than the WT nuclei do. E) To better test differences between different sized nuclei, size classes were formed and compared across the various genetic backgrounds. The groups of nuclei are presented as box plots. Asterisks (\*) indicate the degrees of significance, with an increasing number of asterisks representing higher significance levels.

#### 1.2.5 Supplemental information 5 - Nucleus Roundness in Mutants - Stats

**Supported Figure: 2A-E** In order to test if the different genetic backgrounds differ in their nucleus roundness a One Way Analysis of Variance was initiated on the circularity data obtained for all the backgrounds with the data separated into different cell size classes (small, medium, large). Test were performed with SigmaPlot 12.5. These cell size classes were defined based on nucleus area in the 2D z-projections and the correlation equation (2D projected nucleus area vs. 2D projected cell area) according to Erickson et al., 2017. The program used for the analysis was SigmaPlot 12.5. Because all the normality tests failed (see below) analysis for small, medium and large cells was continued as a Kruskal-Wallis One Way Analysis of Variance on Ranks. The results of the test were as follows:

• small: Normality Test (Shapiro-Wilk) Failed ($P < 0.050$); small: $H = 24.802$ with 4 degrees of freedom. ($P = <0.001$) = The differences in the median values among the treatment groups are greater than would be expected by chance; there is a statistically significant difference ($P = <0.001$)
• medium: Normality Test (Shapiro-Wilk) Failed ($P < 0.050$); medium: $H = 79.228$ with 4 degrees of freedom. ($P = <0.001$) = The differences in the median values among the treatment groups are greater than would be expected by chance; there is a statistically significant difference ($P = <0.001$)
• large: Normality Test (Shapiro-Wilk) Failed ($P < 0.050$) ; large: $H = 196.667$ with 4 degrees of freedom. ($P = <0.001$) = The differences in the median values among the treatment groups are greater than would be expected by chance; there is a statistically significant difference ($P = <0.001$)

| Group | N | Missing | Median | 25% | 75% |
| --- | --- | --- | --- | --- | --- |
| WT-small | 43 | 0 | 0.929 | 0.899 | 0.942 |
| 3KO-small | 37 | 0 | 0.917 | 0.879 | 0.934 |
| 4KO-small | 65 | 0 | 0.909 | 0.830 | 0.933 |
| wit12-small | 35 | 0 | 0.935 | 0.923 | 0.946 |
| xi-i-small | 19 | 0 | 0.940 | 0.926 | 0.952 |
| WT-medium | 66 | 0 | 0.863 | 0.815 | 0.908 |
| 3KO-medium | 49 | 0 | 0.813 | 0.756 | 0.885 |
| 4KO-medium | 125 | 0 | 0.833 | 0.763 | 0.876 |
| wit12-medium | 40 | 0 | 0.915 | 0.902 | 0.928 |
| xi-i-medium | 28 | 0 | 0.917 | 0.864 | 0.928 |
| WT-large | 71 | 0 | 0.671 | 0.602 | 0.747 |
| 3KO-large | 64 | 0 | 0.613 | 0.498 | 0.755 |
| 4KO-large | 99 | 0 | 0.838 | 0.773 | 0.876 |
| wit12-large | 58 | 0 | 0.899 | 0.875 | 0.909 |
| xi-i-large | 34 | 0 | 0.889 | 0.873 | 0.905 |

**Table 1.5:** Parameters describing the data used to test roundness in different pn transgenic mutant lines: Group = name of the genetic background; N = number of considered nuclei; Missing = missing values; Median = median of all roundness values of the respective genetic background; 25% and 75% = 25 and 75 percentile, describing the lower and upper limit of the boxes in figure 1.1 on the previous page.

| Comparison | Diff of Ranks | Q | P<0.05 |
| --- | --- | --- | --- |
| xi-i-small vs 4KO-small | 51.246 | 3.412 | Yes |
| xi-i-small vs 3KO-small | 46.157 | 2.840 | Yes |
| xi-i-small vs WT-small | 27.394 | 1.727 | No |
| xi-i-small vs wit12-small | 2.527 | 0.154 | Do Not Test |
| wit12-small vs 4KO-small | 48.719 | 4.035 | Yes |
| wit12-small vs 3KO-small | 43.630 | 3.213 | Yes |
| wit12-small vs WT-small | 24.866 | 1.897 | Do Not Test |
| WT-small vs 4KO-small | 23.852 | 2.107 | No |
| WT-small vs 3KO-small | 18.764 | 1.453 | Do Not Test |
| 3KO-small vs 4KO-small | 5.089 | 0.429 | Do Not Test |
| wit12-medium vs 3KO-medium | 119.652 | 6.305 | Yes |
| wit12-medium vs 4KO-medium | 116.703 | 7.214 | Yes |
| wit12-medium vs WT-medium | 75.496 | 4.231 | Yes |
| wit12-medium vs xi-i-medium | 11.948 | 0.544 | No |
| xi-i-medium vs 3KO-medium | 107.704 | 5.105 | Yes |
| xi-i-medium vs 4KO-medium | 104.754 | 5.626 | Yes |
| xi-i-medium vs WT-medium | 63.548 | 3.164 | Yes |
| WT-medium vs 3KO-medium | 44.156 | 2.629 | No |
| WT-medium vs 4KO-medium | 41.207 | 3.041 | Do Not Test |
| 4KO-medium vs 3KO-medium | 2.950 | 0.197 | Do Not Test |
| wit12-large vs 3KO-large | 191.307 | 11.196 | Yes |
| wit12-large vs WT-large | 170.720 | 10.234 | Yes |
| wit12-large vs 4KO-large | 80.246 | 5.149 | Yes |
| wit12-large vs xi-i-large | 15.996 | 0.786 | No |
| xi-i-large vs 3KO-large | 175.311 | 8.765 | Yes |
| xi-i-large vs WT-large | 154.723 | 7.871 | Yes |
| xi-i-large vs 4KO-large | 64.249 | 3.429 | Yes |
| 4KO-large vs 3KO-large | 111.062 | 7.347 | Yes |
| 4KO-large vs WT-large | 90.474 | 6.172 | Yes |
| WT-large vs 3KO-large | 20.588 | 1.267 | No |

Table 1.6: Pairwise Multiple Comparison Procedures (Dunn's Method); Comparison = compared groups; Diff of Ranks = difference of ranks; Q = the difference between the two means divided by the standard error of that difference; P = p-value of each pair; $p > 0.050$ = significance statement by Sigmaplot 'Yes' states that the difference is larger than it would be expected by chance, 'No' states that the difference can be explained by chance, 'Do Not Test' states that the previous results of paired tests make the test of this comparison obsolete and is therefore not performed. Note: The multiple comparisons on ranks do not include an adjustment for ties.

#### 1.2.6 Supplemental information 6 - Movies for Nucleus Velocity Measurements

**Supported Figure: 2F** Table 1.7 shows the number of movies used to analyse nucleus velocity. Due to expression level only a selection of created transgenic lines was suitable for time lapse confocal microscopy. In order to avoid positional effects of the T-DNA in cases where only one suitable line was available movies were recorded using heterozygous plants.

| Genotype | Locus | Insertion line | Plant lines | n movies |
| --- | --- | --- | --- | --- |
| WT - Col 0 | - | - | 2, 3, 10, 18 | line10 = 9<br>line 2 = 3<br>line 3 = 1 |
| xi-i<sup>1</sup> | At4g33200 | SALK_082443 | 6, 12, 17 | line 4 = 8 |
| wit1_wit2<sup>3</sup> | At5g11390<br>At1g68910 | GABI-470E06,<br>SALK_127765 | 8, 12, 13 | line 8 = 12 |
| 3KO<sup>3</sup> xi-1, xi-2,<br>xi-k | At1g17580,<br>At5g43900,<br>At5g20490 | SALK_055785,<br>SALK_019031,<br>SALK_067972 | 2, 3, 7 | line 3 = 13 |
| 4KO<sup>2</sup> xi-1, xi-2,<br>xi-k, xi-i | At1g17580<br>At5g43900<br>At5g20490<br>At4g33200 | SALK_055785<br>SALK_019031<br>SALK_067972<br>SALK_082443 | 1, 2, 3 | line1 = 11 |

Table 1.7: Number of movies used for nucleus speed calculation. Seeds were provided by: The Nottingham Arabidopsis Stock Centre (NASC), http://arabidopsis.info/<sup>1</sup> and Valerian Dolja, Oregon State University, Corvallis, Oregon, USA, published in Peremyslov et al. (2010)<sup>2</sup>. Ikuko Hara-Nishimura, Department of Botany, Graduate School of Science, Kyoto University, Kyoto 606-8502, Japan, published in Tamura et al., 2013. Originally created and published in Zhao et al., 2008<sup>3</sup>.

##### 1.2.7 Supplemental information 7 - Cell size and nucleus size correlation

**Supported Figure: 2F** In order to assign in the recorded confocal laser scanning microscope recorded time lapse experiments nuclei to a certain cell size we used the in the upper epidermis of *A. thaliana* existing correlation between nucleus size and cell size. For this of one pn transgenic line was chosen and the cell size as well as the corresponding nucleus size was masured in maximum projected fluorescence images. Figure 1.2 shows the results of the correlation measurements in the different genetic backgrounds. Wildtype results were added to all mutant background graphs in grey as a reference. For each genetick background a fitting curve was chosen to optimise $R^2$. The respective euqtions as well as $R^2$ values are shown in SFigure 1.2. For visualizing the different cell size catagories in each plot a white bar for small, a light greybar for medium and a darker grey bar for large cell was added to the background.

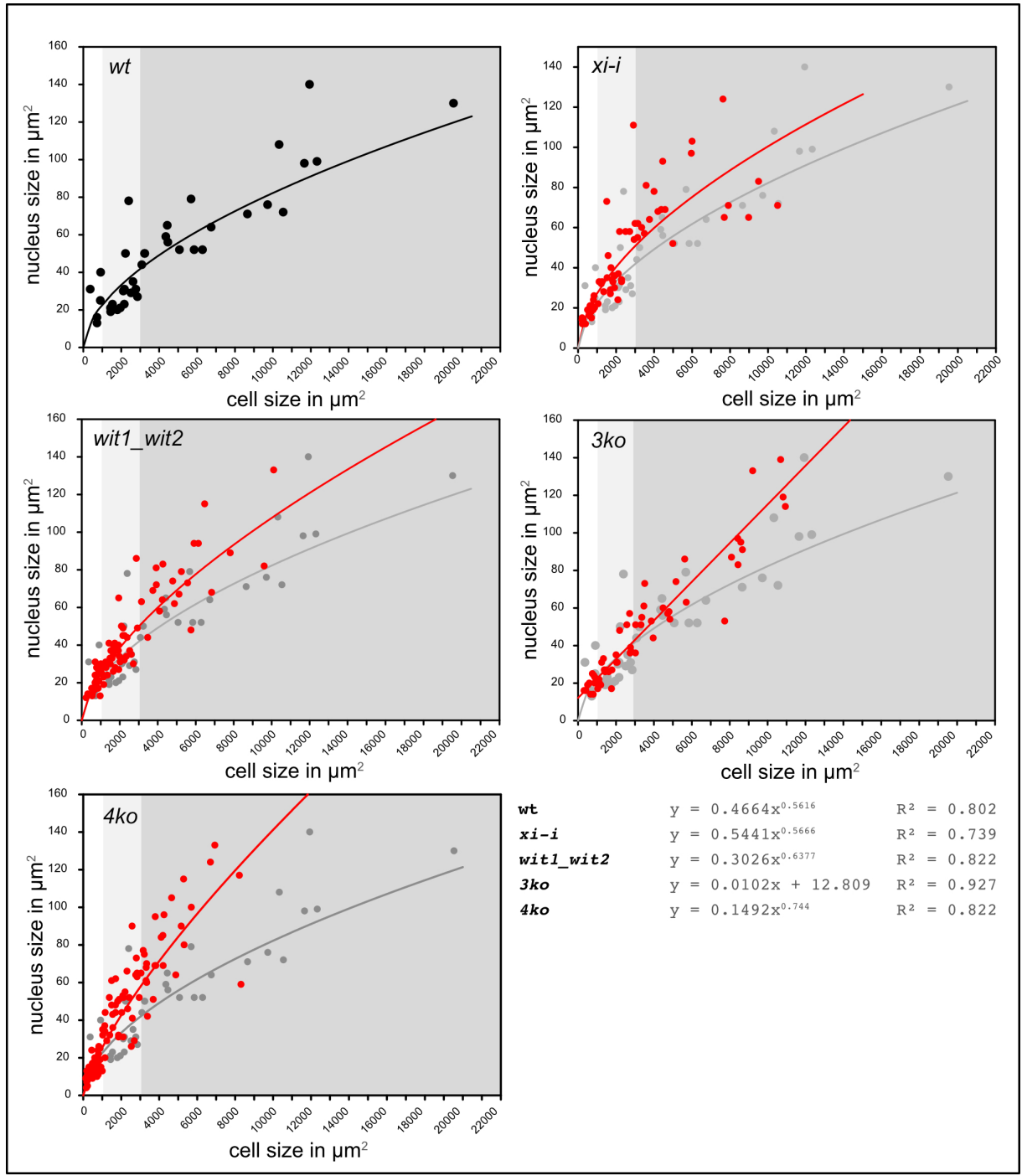

Figure 1.2: Scatter plot for measurements of nucleus size and cell size in columbia wildtype and the different mutant backgrounds; plot background corresponds to the different cell size classes, white = small cells, light grey = medium cells, dark grey = large cells; black and gray dots = wildtype nuclei, red dots = mutant nuclei; lines represent fitting curves for all nuclei of the respective genetic background

##### 1.2.8 Supplemental information 8 - Nucleus Speed in pn - Stats

**Supported Figure: 2F** Stats for nucleus speed measurements in pn transgenic *A. thaliana* lines. Tests were performed with Sigmaplot. Table 1.8, 1.9 and 1.10 summarize the result of the One Way Analysis of Variance performed to test the differences in nucleus speed. Calculations were done with SigmaPlot 12.5. Normality Test (Shapiro-Wilk) Failed ($P < 0.050$) therefore an ANOVA on Ranks Kruskal-Wallis One Way Analysis of Variance on Ranks was started.

$H = 719.005$ with 14 degrees of freedom. The test showed that ($P = <0.001$) the differences in the median values among the treatment groups are greater than would be expected by chance; there is a statistically significant difference ($P = <0.001$) To isolate the group or groups that differ from the others a multiple comparison procedure was used with the Dunn's Method as post-hoc test. For details see table 1.9 and table 1.10.

| Group | N | Missing | Median | 25% | 75% |
| --- | --- | --- | --- | --- | --- |
| WT s | 35 | 0 | 17.046 | 11.534 | 22.550 |
| xi-i s | 51 | 0 | 8.407 | 6.053 | 12.470 |
| wit12 s | 58 | 0 | 9.249 | 6.814 | 11.811 |
| 3ko s | 77 | 0 | 15.784 | 12.984 | 19.500 |
| 4ko s | 172 | 0 | 4.152 | 3.156 | 5.771 |
| WT m | 61 | 0 | 24.187 | 20.770 | 31.571 |
| xi-i m | 56 | 0 | 11.063 | 8.082 | 15.772 |
| wit12 m | 53 | 0 | 12.105 | 9.347 | 15.204 |
| 3ko m | 67 | 0 | 20.537 | 16.591 | 24.268 |
| 4ko m | 105 | 0 | 5.650 | 4.396 | 7.016 |
| WT l | 43 | 0 | 29.500 | 25.543 | 39.090 |
| xi-i l | 33 | 0 | 17.838 | 10.811 | 25.221 |
| wit12 l | 42 | 0 | 22.437 | 14.346 | 31.655 |
| 3ko l | 44 | 0 | 21.978 | 18.070 | 28.200 |
| 4ko l | 65 | 0 | 5.786 | 4.900 | 6.937 |

Table 1.8: Discriptive stats for nucleus speed measurements.

| Comparison | Diff of Ranks | Q | P<0.05 | Comparison | Diff of Ranks | Q | P<0.05 |
| --- | --- | --- | --- | --- | --- | --- | --- |
| WT l vs 4ko s | 712.221 | 15.034 | Yes | wit12 l in vs 4ko s | 587.726 | 12.290 | Yes |
| WT l vs 4ko m | 629.471 | 12.513 | Yes | wit12 l in vs 4ko m | 504.976 | 9.955 | Yes |
| WT l vs 4ko l | 623.040 | 11.407 | Yes | wit12 l in vs 4ko l | 498.545 | 9.063 | Yes |
| WT l vs xi-i s | 465.366 | 8.090 | Yes | wit12 l in vs xi-i s in | 340.871 | 5.888 | Yes |
| WT l vs wit12- s | 448.528 | 8.022 | Yes | wit12 l in vs wit12 s | 324.033 | 5.756 | Yes |
| WT l vs xi-i m | 356.745 | 6.332 | Yes | wit12 l in vs xi-i m i | 232.250 | 4.095 | Yes |
| WT l vs wit12 m | 335.190 | 5.878 | Yes | wit12 l in vs wit12 m | 210.695 | 3.671 | Yes |
| WT l vs 3ko s | 232.443 | 4.394 | Yes | wit12 l in vs 3ko s | 107.948 | 2.025 | Do Not Test |
| WT l vs xi-i l | 231.937 | 3.607 | Yes | wit12 l in vs xi-i l | 107.442 | 1.662 | Do Not Test |
| WT l vs WT s | 212.866 | 3.365 | No | wit12 l in μm/h vs<br>WT s | 88.371 | 1.390 | Do Not Test |
| WT l vs 3ko m | 138.373 | 2.549 | Do Not Test | wit12 l in vs 3ko m | 13.878 | 0.254 | Do Not Test |
| WT l vs wit12 l | 124.495 | 2.065 | Do Not Test | 3ko m vs 4ko s | 573.847 | 14.341 | Yes |
| WT l vs 3ko l | 103.414 | 1.736 | Do Not Test | 3ko m vs 4ko m | 491.098 | 11.304 | Yes |
| WT l vs WT m | 71.127 | 1.286 | Do Not Test | 3ko m vs 4ko l | 484.667 | 10.019 | Yes |
| WT m vs 4ko s | 641.094 | 15.483 | Yes | 3ko m vs xi-i s in | 326.993 | 6.333 | Yes |
| WT m vs 4ko m | 558.344 | 12.482 | Yes | 3ko m vs wit12 s | 310.155 | 6.224 | Yes |
| WT m vs 4ko l | 551.913 | 11.143 | Yes | 3ko m vs xi-i m i | 218.372 | 4.341 | Yes |
| WT m vs xi-i s | 394.239 | 7.478 | Yes | 3ko m vs wit12 m | 196.817 | 3.853 | Yes |
| WT m vs wit12- s | 377.401 | 7.406 | Yes | 3ko m vs 3ko s | 94.070 | 2.026 | Do Not Test |
| WT m vs xi-i m | 285.618 | 5.554 | Yes | 3ko m vs xi-i l | 93.563 | 1.583 | Do Not Test |
| WT m vs wit12 m | 264.063 | 5.061 | Yes | 3ko m vs WT s | 74.493 | 1.286 | Do Not Test |
| WT m vs 3ko s | 161.316 | 3.387 | No | WT s vs 4ko s | 499.354 | 9.692 | Yes |
| WT m vs xi-i l | 160.809 | 2.678 | Do Not Test | WT s vs 4ko m | 416.605 | 7.682 | Yes |
| WT m vs WT s | 141.739 | 2.406 | Do Not Test | WT s vs 4ko l | 410.174 | 7.041 | Yes |
| WT m vs 3ko m | 67.246 | 1.368 | Do Not Test | WT s vs xi-i s | 252.500 | 4.140 | Yes |
| WT m vs wit12 l | 53.368 | 0.958 | Do Not Test | WT s vs wit12- s | 235.662 | 3.963 | Yes |
| WT m vs 3ko l | 32.287 | 0.587 | Do Not Test | WT s vs xi-i m | 143.879 | 2.403 | No |
| 3ko l vs 4ko s | 608.807 | 12.970 | Yes | WT s vs wit12 m | 122.324 | 2.021 | Do Not Test |
| 3ko l vs 4ko m | 526.057 | 10.543 | Yes | WT s vs 3ko s | 19.577 | 0.346 | Do Not Test |
| 3ko l vs 4ko l | 519.626 | 9.580 | Yes | WT s vs xi-i l | 19.070 | 0.283 | Do Not Test |
| Do Not Test | 361.952 | 6.331 | Yes | xi-i l in vs 4ko s | 480.284 | 9.096 | Yes |
| 3ko l vs wit12- s | 345.114 | 6.213 | Yes | xi-i l in vs 4ko m | 397.535 | 7.169 | Yes |
| 3ko l vs xi-i m i | 253.331 | 4.526 | Yes | xi-i l in μm/h vs 4ko l | 391.103 | 6.585 | Yes |
| 3ko l vs wit12 m | 231.777 | 4.090 | Yes | xi-i l in vs xi-i s in | 233.430 | 3.761 | Yes |
| 3ko l vs 3ko s | 129.029 | 2.457 | Do Not Test | xi-i l in vs wit12 s | 216.592 | 3.575 | Yes |
| 3ko l vs xi-i l | 128.523 | 2.009 | Do Not Test | xi-i l in vs xi-i m i | 124.808 | 2.047 | Do Not Test |
| 3ko l vs WT s | 109.453 | 1.739 | Do Not Test | xi-i l in vs wit12 m | 103.254 | 1.676 | Do Not Test |
| 3ko l vs 3ko m | 34.960 | 0.648 | Do Not Test | xi-i l in vs 3ko s | 0.506 | 0.00876 | Do Not Test |
| 3ko l vs wit12 l | 21.081 | 0.352 | Do Not Test |  |  |  |  |

**Table 1.9:** Post-hoc test results of the Dunn's Method post-hoc test. s = small sized cells, m = medium sized cells, l = large sized cells; Comparison = pairs of groups being compared, Diff of Ranks = Difference of Ranks (the difference in the rank sums between the two groups being compared), Q = test statistic for the pairwise comparison, P<0.05 = indicates whether the difference is statistically significant. "Yes" means that the P-value is less than 0.05, indicating a statistically significant difference between the groups. "Do Not Test" indicates that a particular comparison or analysis was deemed unnecessary.

| Comparison | Diff of Ranks | Q | P<0.05 | Comparison | Diff of Ranks | Q | P<0.05 |
| --- | --- | --- | --- | --- | --- | --- | --- |
| 3ko s vs 4ko s | 479.778 | 12.593 | Yes | wit12- s vs 4ko s | 263.693 | 6.250 | Yes |
| 3ko s vs 4ko m | 397.028 | 9.524 | Yes | wit12- s vs 4ko m | 180.943 | 3.981 | Yes |
| 3ko s vs 4ko l | 390.597 | 8.346 | Yes | wit12- s vs 4ko l | 174.512 | 3.477 | No |
| 3ko s vs xi-i s | 232.923 | 4.643 | Yes | wit12- s vs xi-i s | 16.838 | 0.316 | Do Not Test |
| 3ko s vs wit12- s | 216.085 | 4.473 | Yes | xi-i s in vs 4ko s | 246.855 | 5.572 | Yes |
| 3ko s vs xi-i m i | 124.302 | 2.547 | Do Not Test | xi-i s in vs 4ko m | 164.105 | 3.460 | No |
| 3ko s vs wit12 m | 102.747 | 2.072 | Do Not Test | xi-i s in vs 4ko l | 157.674 | 3.034 | Do Not Test |
| wit12 m vs 4ko s | 377.030 | 8.637 | Yes | 4ko l vs 4ko s | 89.181 | 2.204 | No |
| wit12 m vs 4ko m | 294.281 | 6.286 | Yes | 4ko l vs 4ko m | 6.431 | 0.147 | Do Not Test |
| wit12 m vs 4ko l | 287.850 | 5.598 | Yes | 4ko m vs 4ko s | 82.750 | 2.405 | Do Not Test |
| wit12 m vs xi-i s | 130.176 | 2.389 | No |  |  |  |  |
| wit12 m vs wit12 s | 113.338 | 2.147 | Do Not Test |  |  |  |  |
| wit12 m vs xi-i m | 21.555 | 0.405 | Do Not Test |  |  |  |  |
| xi-i m i vs 4ko s | 355.476 | 8.316 | Yes |  |  |  |  |
| xi-i m i vs 4ko m | 272.726 | 5.932 | Yes |  |  |  |  |
| xi-i m i vs 4ko l | 266.295 | 5.257 | Yes |  |  |  |  |
| xi-i m i vs xi-i s in | 108.621 | 2.020 | Do Not Test |  |  |  |  |
| xi-i m i vs wit12 s | 91.783 | 1.763 | Do Not Test |  |  |  |  |

Table 1.10: Post-hoc test results of the Dunn's Method post-hoc test. s = small sized cells, m = medium sized cells, l = large sized cells; Comparison = pairs of groups being compared, Diff of Ranks = Difference of Ranks (the difference in the rank sums between the two groups being compared), Q = test statistic for the pairwise comparison, P<0.05 = indicates whether the difference is statistically significant. "Yes" means that the P-value is less than 0.05, indicating a statistically significant difference between the groups. "Do Not Test" indicates that a particular comparison or analysis was deemed unnecessary.

##### 1.2.9 Supplemental information 9 - Stromule Frequency in pn - Stats

**Supported Figure: 2G** Stats for stromule frequency measurements in *plsu4pn* transgenic *A. thaliana* lines. Table 1.11, 1.12 and 1.13 summarize the statistic data for the stromule frequency measurements. Table 1.11 and 1.12 show the data of the ANOVA analysis. Using Sigmaplot 12.5, a one-way analysis of variance (ANOVA) was performed. The normality test (Kolmogorov-Smirnov) passed ($P = 0.184$), but the equal variance test (Brown-Forsythe) failed ($P < 0.050$). Therefore, an ANOVA on ranks was conducted instead. The Kruskal-Wallis one-way analysis of variance on ranks resulted in $H = 81.873$ with 4 degrees of freedom ($P < 0.001$), indicating that the differences in the median values among the treatment groups are statistically significant. To isolate the groups that differ, a multiple comparison procedure using Dunn's method was performed.

| Group | N | Missing | Median | 25% | 75% |
| --- | --- | --- | --- | --- | --- |
| WT | 57 | 0 | 0.369 | 0.321 | 0.413 |
| xi-i | 33 | 0 | 0.298 | 0.275 | 0.351 |
| wit12 | 30 | 0 | 0.299 | 0.269 | 0.325 |
| 3KO | 18 | 0 | 0.301 | 0.25 | 0.33 |
| 4KO | 31 | 0 | 0.189 | 0.165 | 0.224 |

**Table 1.11:** Details of the performed statistical test: N = sample size considered in each group (each sample corresponds to the arcsin transformed average stromule frequency of a single tested leaf, calculated from three image stacks taken for each leaf except for WT here for each mutant 1 WT control plant was grown, imaged and counted, the value represents the average of these plants); Missing = missing values; Median = median value of n arcsin transformed stromule frequency values; 25% and 75% = respective percentiles defining the box boundaries)

| Comparison | Diff of Ranks | Q | P | P<0.050 |
| --- | --- | --- | --- | --- |
| WT vs 4KO | 98.283 | 9.001 | <0.001 | Yes |
| WT vs 3KO | 42.731 | 3.23 | 0.012 | Yes |
| WT vs wit12 | 42.392 | 3.841 | 0.001 | Yes |
| WT vs xi-i | 35.357 | 3.304 | 0.01 | Yes |
| xi-i vs 4KO | 62.926 | 5.142 | <0.001 | Yes |
| xi-i vs 3KO | 7.374 | 0.514 | 1 | No |
| xi-i vs wit12 | 7.035 | 0.57 | 1 | Do Not Test |
| wit12 vs 4KO | 55.891 | 4.46 | <0.001 | Yes |
| wit12 vs 3KO | 0.339 | 0.0232 | 1 | Do Not Test |
| 3KO vs 4KO | 55.552 | 3.831 | 0.001 | Yes |

**Table 1.12:** Comparison = compared groups; Diff of Ranks = difference of ranks; Q = the difference between the two means divided by the standard error of that difference; P = p-value of each pair; $p > 0.050$ = significance statement by Sigmaplot 'Yes' states that the difference is larger than it would be expected by chance, 'No' states that the difference can be explained by chance, 'Do Not Test' states that the previous results of paired tests does make the test obsolete and is therefore not performed.

|  | Mean | absolute value<br>for lower back-transformed<br>CI | absolute value<br>for Upper<br>Backtran-<br>formed<br>CI | difference to<br>Median of<br>Lower 95 CI | difference to<br>Median of<br>Upper 95 CI |
| --- | --- | --- | --- | --- | --- |
| WT all | 0.14 | 0.12 | 0.1308 | 0.0141 | 0.0135 |
| wit12 | 0.10 | 0.08 | 0.0862 | 0.0104 | 0.0099 |
| xii | 0.10 | 0.08 | 0.0913 | 0.0107 | 0.0102 |
| 3KO | 0.10 | 0.07 | 0.0856 | 0.0149 | 0.0138 |
| 4KO | 0.05 | 0.03 | 0.0387 | 0.0076 | 0.0069 |

Table 1.13: Stats for stromule frequency analysis pn transgenic *A. thaliana* lines. The table presents the values after the backtransformation "Lower 95% CI" and "Upper 95% CI" represent the lower and upper bounds of the 95% confidence intervals, before back transformation. The "Mean" column provides the arithmetic mean of the data, before back transformation. The "Absolute value for lower back-transformed CI" and "Absolute value for upper back-transformed CI" columns display the absolute values of the lower and upper bounds of the back-transformed confidence intervals. The columns "Difference to median of lower 95% CI" and "Difference to median of upper 95% CI" show the differences between the confidence interval bounds and their respective median values of the back transformed values. Note that CI stands for confidence intervals.

##### 1.2.10 Supplemental information 10 - Lines for stromule promoting zone measurements

**Supported Figure: 3 and 4** For analysing the impact of the different mutant lines on stromule formation within and outside of the stromule promoting zone for each used genetic background plants of at least three independent transgenic lines were imaged and analysed. Table 1.14 summarizes the imaged lines, total plants and analysed images.

| Genotype | Locus | Insertion line | Plant lines | Total lines | Total plants | Images analyzed |
| --- | --- | --- | --- | --- | --- | --- |
| WT - Col 0 | - | - | 2, 3, 10, 18 | 4 | 18 | 70 |
| xi-i<sup>1</sup> | At4g33200 | SALK_082443 | 6, 12, 17 | 3 | 19 | 31 |
| wit1_wit2<sup>3</sup> | At5g11390<br>At1g68910 | GABI-470E06,<br>SALK_127765 | 8, 12, 13 | 3 | 11 | 32 |
| 4KO<sup>2</sup> xi-1, xi-2,<br>xi-k, xi-i | At1g17580<br>At5g43900<br>At5g20490<br>At4g33200 | SALK_055785<br>SALK_019031<br>SALK_067972<br>SALK_082443 | 1, 2, 3 | 3 | 10 | 28 |

Table 1.14: Seeds were provided by: The Nottingham Arabidopsis Stock Centre (NASC), http://arabidopsis.info/<sup>1</sup> and Valerian Dolja, Oregon State University, Corvallis, Oregon, USA, published in Peremyslov et al. (2010)<sup>2</sup>. Ikuko Hara-Nishimura, Department of Botany, Graduate School of Science, Kyoto University, Kyoto 606-8502, Japan, published in Tamura et al., 2013. Originally created and published in Zhao et al., 2008<sup>3</sup>.

##### 1.2.11 Supplemental information 11 - Plastid distribution - Figure

**Supported Figure: 4A-D** In order to test if the considered mutant alleles have an impact on plastid distribution in general the distance distribution for plastids in all mutants were compared to wt. Figure 1.3 shows the result of this comparison. It is clearly visible that in all plant lines and cell sizes plastids accumulate in the direct vicinity of the nucleus. Residual plastids are spread throughout the cell, depending on the respective cell size. In 4ko plants the proportion of plastids in the cluster seems lower than in other plant lines, but still form a clear cluster.

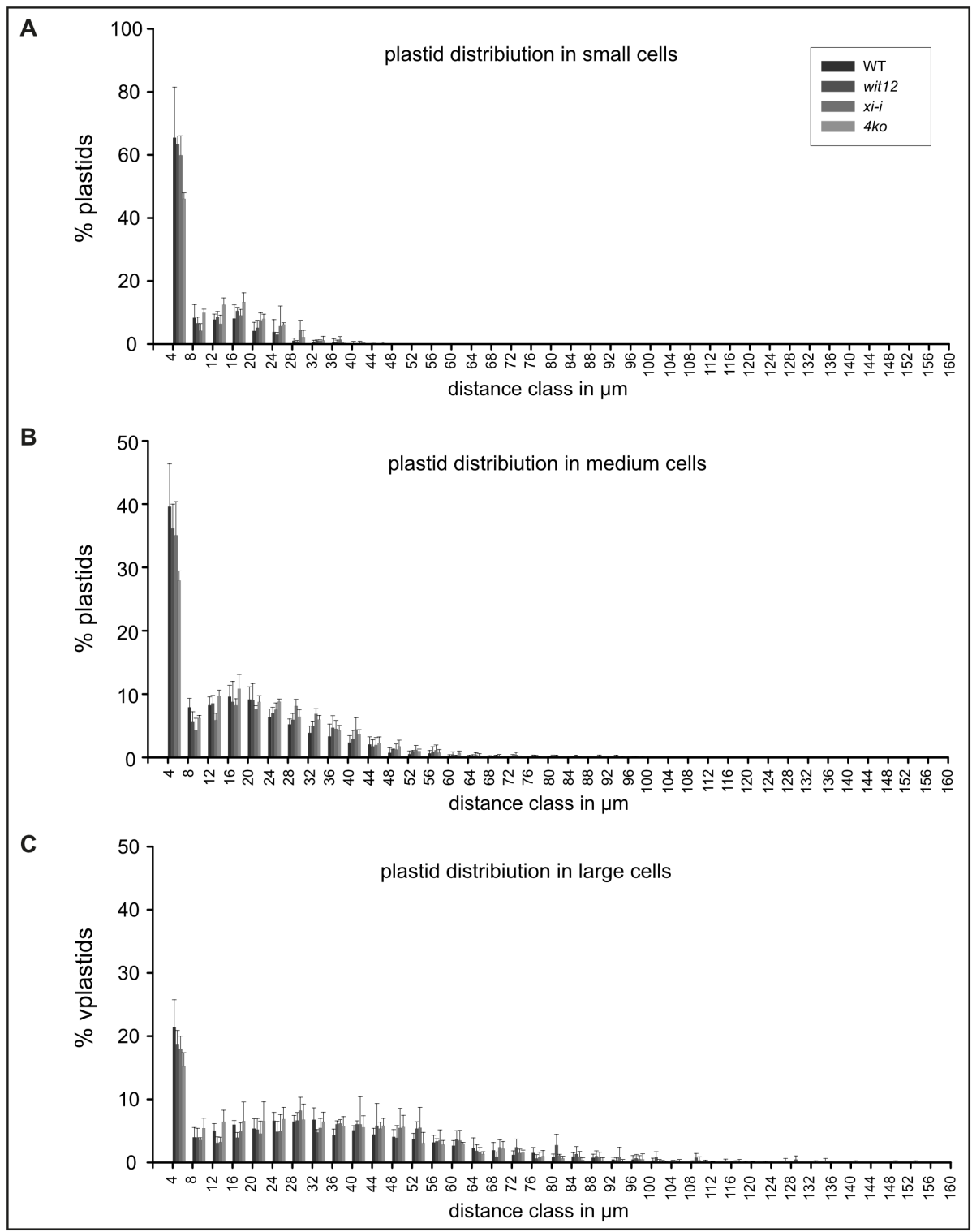

Figure 1.3: Plastid distance distribution relative to the nucleus. A) small cells, B) medium cells and C) large cells. On the x-axis the upper boarder of the created distance class is given (4 = distances from 0 $\mu\text{m}$ to 4$\mu\text{m}$).

##### 1.2.12 Supplemental Information 12 - Plastid distribution - Stats

**Supported Figure: 4A-D** This table provides the statistical data underlying the plastid distribution analysis shown in Figure 4A-D. For each genotype (wt, *xi-i*, *wit1/2*, and *4ko*) and cell size class (large, medium, and small), the percentage of total plastids found within consecutive 4 $\mu\text{m}$ distance bins from the nucleus is given. These values form the basis for the bar charts in Supplemental Information 11 and confirm that plastid distribution relative to the nucleus is broadly similar across all genotypes, supporting the interpretation that differences in SPZ stromule frequency reflect genuine changes in stromule formation propensity rather than redistribution of the plastid population.

| $\mu\text{m}$ | wt lrg | wt m | wt s | wit1/2 lrg | wit1/2 m | wit1/2 s | xi-i lrg | xi-i m | xi-i s | 4ko lrg | 4ko m | 4ko s |
| --- | --- | --- | --- | --- | --- | --- | --- | --- | --- | --- | --- | --- |
| 4 | 21.38 | 39.59 | 65.45 | 18.76 | 36.18 | 63.54 | 18.01 | 35.08 | 59.91 | 15.21 | 27.93 | 46.07 |
| 8 | 3.98 | 7.91 | 8.33 | 3.96 | 5.7 | 6.59 | 3.57 | 4.34 | 4.23 | 5.43 | 6.22 | 9.92 |
| 12 | 5.05 | 8.24 | 7.76 | 3.1 | 8.54 | 8.63 | 3.24 | 5.89 | 6.39 | 6.42 | 9.71 | 12.47 |
| 16 | 5.98 | 9.59 | 8.11 | 3.93 | 8.77 | 10.52 | 4.93 | 8.22 | 9.09 | 6.57 | 10.84 | 13.35 |
| 20 | 5.35 | 9.15 | 4.16 | 5.2 | 9.08 | 5.16 | 4.56 | 7.68 | 7.51 | 6.54 | 8.76 | 7.95 |
| 24 | 6.59 | 6.39 | 3.83 | 4.87 | 6.99 | 2.99 | 4.96 | 7.53 | 5.66 | 6.86 | 8.8 | 6.25 |
| 28 | 6.47 | 5.21 | 1.03 | 6.65 | 5.95 | 0.8 | 8.22 | 8.13 | 4.44 | 6.79 | 6.42 | 2.23 |
| 32 | 6.77 | 3.87 | 0.55 | 4.77 | 4.98 | 1.09 | 5.46 | 6.88 | 1.04 | 6.44 | 5.97 | 1.18 |
| 36 | 4.28 | 3.31 | 0.48 | 6.05 | 4.69 | 0.66 | 6.17 | 4.51 | 1.41 | 5.78 | 4.24 | 0.21 |
| 40 | 5.08 | 2.33 | 0.24 | 6.05 | 2.91 | 0 | 6.03 | 4.41 | 0.32 | 5.57 | 3.64 | 0.18 |
| 44 | 4.42 | 2.04 | 0.07 | 5.81 | 1.73 | 0 | 5.36 | 1.96 | 0 | 5.81 | 2.29 | 0.21 |
| 48 | 4.05 | 0.72 | 0 | 3.9 | 1.27 | 0 | 5.46 | 1.24 | 0 | 5.61 | 1.74 | 0 |
| 52 | 3.7 | 0.52 | 0 | 5.35 | 0.97 | 0 | 5.49 | 1.17 | 0 | 3.09 | 0.99 | 0 |
| 56 | 3.16 | 0.63 | 0 | 3.36 | 0.88 | 0 | 3.51 | 1.15 | 0 | 2.82 | 0.75 | 0 |
| 60 | 2.66 | 0.19 | 0 | 3.65 | 0.46 | 0 | 3.48 | 0.18 | 0 | 2.88 | 0.7 | 0 |
| 64 | 2.29 | 0.16 | 0 | 1.84 | 0.21 | 0 | 1.63 | 0.37 | 0 | 1.33 | 0.28 | 0 |
| 68 | 1.92 | 0.09 | 0 | 0.89 | 0.07 | 0 | 2.4 | 0.18 | 0 | 2.19 | 0.22 | 0 |
| 72 | 1.2 | 0 | 0 | 2.4 | 0.22 | 0 | 1.5 | 0.39 | 0 | 1.55 | 0.21 | 0 |
| 76 | 1.5 | 0.02 | 0 | 0.64 | 0.14 | 0 | 0.86 | 0.12 | 0 | 1 | 0.07 | 0 |
| 80 | 0.85 | 0.05 | 0 | 2.74 | 0.14 | 0 | 1.13 | 0.18 | 0 | 0.65 | 0 | 0 |
| 84 | 0.92 | 0 | 0 | 1.33 | 0.07 | 0 | 0.93 | 0.13 | 0 | 0.4 | 0.07 | 0 |
| 88 | 0.76 | 0 | 0 | 0.99 | 0 | 0 | 0.86 | 0 | 0 | 0.28 | 0.14 | 0 |
| 92 | 0.46 | 0 | 0 | 0.35 | 0 | 0 | 0.88 | 0.18 | 0 | 0.17 | 0 | 0 |
| 96 | 0.46 | 0 | 0 | 0.66 | 0.07 | 0 | 0.53 | 0.07 | 0 | 0.5 | 0 | 0 |
| 100 | 0.29 | 0 | 0 | 0.83 | 0 | 0 | 0.17 | 0 | 0 | 0.12 | 0 | 0 |
| 104 | 0.13 | 0 | 0 | 0.1 | 0 | 0 | 0.17 | 0 | 0 | 0 | 0 | 0 |
| 108 | 0.1 | 0 | 0 | 0.76 | 0 | 0 | 0.33 | 0 | 0 | 0 | 0 | 0 |
| 112 | 0.05 | 0 | 0 | 0 | 0 | 0 | 0 | 0 | 0 | 0 | 0 | 0 |
| 116 | 0.09 | 0 | 0 | 0.1 | 0 | 0 | 0.17 | 0 | 0 | 0 | 0 | 0 |
| 120 | 0.05 | 0 | 0 | 0 | 0 | 0 | 0 | 0 | 0 | 0 | 0 | 0 |
| 124 | 0 | 0 | 0 | 0 | 0 | 0 | 0 | 0 | 0 | 0 | 0 | 0 |
| 128 | 0 | 0 | 0 | 0.46 | 0 | 0 | 0 | 0 | 0 | 0 | 0 | 0 |
| 132 | 0 | 0 | 0 | 0.1 | 0 | 0 | 0 | 0 | 0 | 0 | 0 | 0 |
| 136 | 0 | 0 | 0 | 0 | 0 | 0 | 0 | 0 | 0 | 0 | 0 | 0 |
| 140 | 0 | 0 | 0 | 0.1 | 0 | 0 | 0 | 0 | 0 | 0 | 0 | 0 |
| 144 | 0 | 0 | 0 | 0 | 0 | 0 | 0 | 0 | 0 | 0 | 0 | 0 |
| 148 | 0 | 0 | 0 | 0.1 | 0 | 0 | 0 | 0 | 0 | 0 | 0 | 0 |
| 152 | 0 | 0 | 0 | 0.1 | 0 | 0 | 0 | 0 | 0 | 0 | 0 | 0 |
| 156 | 0 | 0 | 0 | 0 | 0 | 0 | 0 | 0 | 0 | 0 | 0 | 0 |
| 160 | 0 | 0 | 0 | 0 | 0 | 0 | 0 | 0 | 0 | 0 | 0 | 0 |
| 164 | 0 | 0 | 0 | 0.1 | 0 | 0 | 0 | 0 | 0 | 0 | 0 | 0 |

Table 1.15: Plastid distribution in cells of wildtype = wt and three mutant backgrounds. $\mu\text{m}$ = upper boarder of distance class in $\mu\text{m}$, lrg = large sized cells, m = medium sized cells, s = small sized cells

##### 1.2.13 Supplemental Information 13 - Zone Measurements - SF

**Supported Figure: 4A-D** This table provides the numerical values underlying the stromule promoting zone bar charts shown in Figure 4A–D. For each genotype (wt, xi-i, wit1/2, and 4ko) and cell size class (large [L], medium [M], and small [S]), mean stromule frequency (SF), standard deviation (SD), and plastid count (n) are given for consecutive 4 μm distance bins measured from the nucleus surface. Distance zones containing three or fewer plastids were excluded from the analysis to avoid sampling artefacts arising from small denominators (see Supplemental Methods 1 for details).

|  | wt |  |  | wt |  |  | wt |  |  | xi-i |  |  | xi-i |  |  | xi-i |  |  |
| --- | --- | --- | --- | --- | --- | --- | --- | --- | --- | --- | --- | --- | --- | --- | --- | --- | --- | --- |
|  | L |  |  | M |  |  | S |  |  | L |  |  | M |  |  | S |  |  |
| $\mu\text{m}$ | SF | SD | n | SF | SD | n | SF | SD | n | SF | SD | n | SF | SD | n | SF | SD | n |
| 4 | 0.38 | 0.04 | 455 | 0.21 | 0.12 | 1136 | 0.06 | 0.03 | 566 | 0.17 | 0.15 | 64 | 0.04 | 0.04 | 377 | 0.01 | 0.01 | 249 |
| 8 | 0.24 | 0.14 | 85 | 0.17 | 0.09 | 230 | 0.03 | 0.04 | 59 | 0.21 | 0.18 | 13 | 0.04 | 0.07 | 46 | 0.00 | 0.00 | 20 |
| 12 | 0.08 | 0.04 | 107 | 0.02 | 0.02 | 232 | 0.00 | 0.00 | 65 | 0.07 | 0.12 | 11 | 0.02 | 0.04 | 64 | 0.00 | 0.00 | 23 |
| 16 | 0.05 | 0.05 | 125 | 0.04 | 0.07 | 259 | 0.02 | 0.04 | 59 | 0.00 | 0.00 | 20 | 0.01 | 0.03 | 85 | 0.00 | 0.00 | 36 |
| 20 | 0.03 | 0.06 | 116 | 0.03 | 0.04 | 240 | 0.03 | 0.07 | 31 | 0.00 | 0.00 | 17 | 0.00 | 0.00 | 80 | 0.00 | 0.00 | 28 |
| 24 | 0.03 | 0.07 | 136 | 0.01 | 0.03 | 173 | 0.02 | 0.05 | 27 | 0.00 | 0.00 | 21 | 0.00 | 0.00 | 75 | 0.00 | 0.00 | 25 |
| 28 | 0.04 | 0.04 | 136 | 0.00 | 0.01 | 146 | 0.00 | 0.00 | 8 | 0.00 | 0.00 | 28 | 0.02 | 0.02 | 84 | 0.00 | 0.00 | 14 |
| 32 | 0.02 | 0.05 | 141 | 0.03 | 0.06 | 102 | 0.00 | 0.00 | 4 | 0.05 | 0.08 | 22 | 0.00 | 0.00 | 69 | 0.00 | 0.00 | 4 |
| 36 | 0.00 | 0.00 | 91 | 0.01 | 0.02 | 79 | 0.00 | 0.00 | 2 | 0.00 | 0.00 | 23 | 0.00 | 0.00 | 45 | 0.00 | 0.00 | 5 |
| 40 | 0.01 | 0.03 | 107 | 0.00 | 0.00 | 58 | 0.00 | 0.00 | 1 | 0.00 | 0.00 | 21 | 0.00 | 0.00 | 41 | 0.00 | 0.00 | 2 |
| 44 | 0.02 | 0.06 | 91 | 0.00 | 0.00 | 50 | 0.00 | 0.00 | 1 | 0.00 | 0.00 | 21 | 0.00 | 0.00 | 17 | 0.00 | 0.00 | 0 |
| 48 | 0.01 | 0.03 | 84 | 0.00 | 0.00 | 20 | 0.00 | 0.00 | 0 | 0.00 | 0.00 | 20 | 0.00 | 0.00 | 13 | 0.00 | 0.00 | 0 |
| 52 | 0.04 | 0.06 | 75 | 0.00 | 0.00 | 18 | 0.00 | 0.00 | 0 | 0.00 | 0.00 | 18 | 0.00 | 0.00 | 11 | 0.00 | 0.00 | 0 |
| 56 | 0.01 | 0.03 | 65 | 0.00 | 0.00 | 16 | 0.00 | 0.00 | 0 | 0.00 | 0.00 | 11 | 0.00 | 0.00 | 10 | 0.00 | 0.00 | 0 |
| 60 | 0.00 | 0.00 | 55 | 0.00 | 0.00 | 7 | 0.00 | 0.00 | 0 | 0.06 | 0.10 | 12 | 0.00 | 0.00 | 2 | 0.00 | 0.00 | 0 |
| 64 | 0.01 | 0.03 | 45 | 0.00 | 0.00 | 6 | 0.00 | 0.00 | 0 | 0.00 | 0.00 | 7 | 0.00 | 0.00 | 4 | 0.00 | 0.00 | 0 |
| 68 | 0.06 | 0.14 | 38 | 0.00 | 0.00 | 2 | 0.00 | 0.00 | 0 | 0.00 | 0.00 | 10 | 0.00 | 0.00 | 2 | 0.00 | 0.00 | 0 |
| 72 | 0.00 | 0.00 | 24 | 0.00 | 0.00 | 0 | 0.00 | 0.00 | 0 | 0.00 | 0.00 | 5 | 0.00 | 0.00 | 3 | 0.00 | 0.00 | 0 |
| 76 | 0.00 | 0.00 | 31 | 0.00 | 0.00 | 1 | 0.00 | 0.00 | 0 | 0.00 | 0.00 | 4 | 0.00 | 0.00 | 1 | 0.00 | 0.00 | 0 |
| 80 | nv | nv | 17 | 0.00 | 0.00 | 2 | 0.00 | 0.00 | 0 | 0.00 | 0.00 | 4 | 0.00 | 0.00 | 2 | 0.00 | 0.00 | 0 |
| 84 | 0.00 | 0.00 | 18 | 0.00 | 0.00 | 0 | 0.00 | 0.00 | 0 | 0.00 | 0.00 | 4 | 0.00 | 0.00 | 1 | 0.00 | 0.00 | 0 |
| 88 | 0.06 | 0.14 | 15 | 0.00 | 0.00 | 0 | 0.00 | 0.00 | 0 | 0.00 | 0.00 | 4 | 0.00 | 0.00 | 0 | 0.00 | 0.00 | 0 |
| 92 | nv | nv | 10 | 0.00 | 0.00 | 0 | 0.00 | 0.00 | 0 | 0.00 | 0.00 | 2 | 0.00 | 0.00 | 2 | 0.00 | 0.00 | 0 |
| 96 | 0.00 | 0.00 | 9 | 0.00 | 0.00 | 0 | 0.00 | 0.00 | 0 | 0.00 | 0.00 | 2 | 0.00 | 0.00 | 1 | 0.00 | 0.00 | 0 |
| 100 | 0.00 | 0.00 | 6 | 0.00 | 0.00 | 0 | 0.00 | 0.00 | 0 | 0.00 | 0.00 | 1 | 0.00 | 0.00 | 0 | 0.00 | 0.00 | 0 |
| 104 | 0.00 | 0.00 | 3 | 0.00 | 0.00 | 0 | 0.00 | 0.00 | 0 | 0.00 | 0.00 | 1 | 0.00 | 0.00 | 0 | 0.00 | 0.00 | 0 |
| 108 | 0.00 | 0.00 | 2 | 0.00 | 0.00 | 0 | 0.00 | 0.00 | 0 | 0.00 | 0.00 | 2 | 0.00 | 0.00 | 0 | 0.00 | 0.00 | 0 |
| 112 | 0.00 | 0.00 | 1 | 0.00 | 0.00 | 0 | 0.00 | 0.00 | 0 | 0.00 | 0.00 | 0 | 0.00 | 0.00 | 0 | 0.00 | 0.00 | 0 |
| 116 | 0.00 | 0.00 | 2 | 0.00 | 0.00 | 0 | 0.00 | 0.00 | 0 | 0.00 | 0.00 | 1 | 0.00 | 0.00 | 0 | 0.00 | 0.00 | 0 |
| 120 | 0.00 | 0.00 | 1 | 0.00 | 0.00 | 0 | 0.00 | 0.00 | 0 | 0.00 | 0.00 | 0 | 0.00 | 0.00 | 0 | 0.00 | 0.00 | 0 |
| 124 | 0.00 | 0.00 | 0 | 0.00 | 0.00 | 0 | 0.00 | 0.00 | 0 | 0.00 | 0.00 | 0 | 0.00 | 0.00 | 0 | 0.00 | 0.00 | 0 |
| 128 | 0.00 | 0.00 | 0 | 0.00 | 0.00 | 0 | 0.00 | 0.00 | 0 | 0.00 | 0.00 | 0 | 0.00 | 0.00 | 0 | 0.00 | 0.00 | 0 |
| 132 | 0.00 | 0.00 | 0 | 0.00 | 0.00 | 0 | 0.00 | 0.00 | 0 | 0.00 | 0.00 | 0 | 0.00 | 0.00 | 0 | 0.00 | 0.00 | 0 |
| 136 | 0.00 | 0.00 | 0 | 0.00 | 0.00 | 0 | 0.00 | 0.00 | 0 | 0.00 | 0.00 | 0 | 0.00 | 0.00 | 0 | 0.00 | 0.00 | 0 |
| 140 | 0.00 | 0.00 | 0 | 0.00 | 0.00 | 0 | 0.00 | 0.00 | 0 | 0.00 | 0.00 | 0 | 0.00 | 0.00 | 0 | 0.00 | 0.00 | 0 |
| 144 | 0.00 | 0.00 | 0 | 0.00 | 0.00 | 0 | 0.00 | 0.00 | 0 | 0.00 | 0.00 | 0 | 0.00 | 0.00 | 0 | 0.00 | 0.00 | 0 |
| 148 | 0.00 | 0.00 | 0 | 0.00 | 0.00 | 0 | 0.00 | 0.00 | 0 | 0.00 | 0.00 | 0 | 0.00 | 0.00 | 0 | 0.00 | 0.00 | 0 |
| 152 | 0.00 | 0.00 | 0 | 0.00 | 0.00 | 0 | 0.00 | 0.00 | 0 | 0.00 | 0.00 | 0 | 0.00 | 0.00 | 0 | 0.00 | 0.00 | 0 |
| 156 | 0.00 | 0.00 | 0 | 0.00 | 0.00 | 0 | 0.00 | 0.00 | 0 | 0.00 | 0.00 | 0 | 0.00 | 0.00 | 0 | 0.00 | 0.00 | 0 |
| 160 | 0.00 | 0.00 | 0 | 0.00 | 0.00 | 0 | 0.00 | 0.00 | 0 | 0.00 | 0.00 | 0 | 0.00 | 0.00 | 0 | 0.00 | 0.00 | 0 |
| 164 | 0.00 | 0.00 | 0 | 0.00 | 0.00 | 0 | 0.00 | 0.00 | 0 | 0.00 | 0.00 | 0 | 0.00 | 0.00 | 0 | 0.00 | 0.00 | 0 |

**Table 1.16:** Average stromule frequency values (SF) and corresponding standard deviation (SD) as well as number of scored plastids (n) for each distance zone from the nucleus ($\mu\text{m}$) for wt and xi-i lines. Values used to plot Figure 5A and B. wt = wildtype, xi-i = xi-i mutant lines, L = large cells, M = medium cells, S = small cells

|  | w12 |  |  | w12 |  |  | w12 |  |  | 4ko |  |  | 4ko |  |  | 4ko |  |  |
| --- | --- | --- | --- | --- | --- | --- | --- | --- | --- | --- | --- | --- | --- | --- | --- | --- | --- | --- |
|  | L |  |  | M |  |  | S |  |  | L |  |  | M |  |  | S |  |  |
| $\mu\text{m}$ | SF | SD | n | SF | SD | n | SF | SD | n | SF | SD | n | SF | SD | n | SF | SD | n |
| 4 | 0.20 | 0.07 | 101 | 0.03 | 0.03 | 390 | 0.00 | 0.00 | 336 | 0.04 | 0.06 | 112 | 0.01 | 0.01 | 395 | 0.00 | 0.01 | 215 |
| 8 | 0.13 | 0.23 | 19 | 0.05 | 0.04 | 60 | 0.02 | 0.04 | 33 | 0.00 | 0.00 | 38 | 0.01 | 0.02 | 88 | 0.00 | 0.00 | 47 |
| 12 | 0.20 | 0.08 | 20 | 0.02 | 0.02 | 92 | 0.00 | 0.00 | 44 | 0.02 | 0.03 | 45 | 0.01 | 0.02 | 138 | 0.00 | 0.00 | 57 |
| 16 | 0.02 | 0.04 | 23 | 0.02 | 0.03 | 104 | 0.01 | 0.02 | 56 | 0.02 | 0.04 | 45 | 0.01 | 0.01 | 154 | 0.02 | 0.03 | 60 |
| 20 | 0.00 | 0.00 | 31 | 0.02 | 0.04 | 101 | 0.00 | 0.00 | 30 | 0.00 | 0.00 | 50 | 0.01 | 0.02 | 124 | 0.00 | 0.00 | 36 |
| 24 | 0.02 | 0.03 | 30 | 0.00 | 0.00 | 79 | 0.00 | 0.00 | 15 | 0.02 | 0.03 | 49 | 0.02 | 0.04 | 125 | 0.00 | 0.00 | 29 |
| 28 | 0.00 | 0.00 | 38 | 0.05 | 0.05 | 64 | 0.00 | 0.00 | 4 | 0.02 | 0.03 | 51 | 0.01 | 0.02 | 91 | 0.04 | 0.07 | 12 |
| 32 | 0.02 | 0.04 | 27 | 0.00 | 0.00 | 53 | 0.00 | 0.00 | 6 | 0.00 | 0.00 | 48 | 0.01 | 0.02 | 85 | 0.00 | 0.00 | 6 |
| 36 | 0.04 | 0.07 | 33 | 0.01 | 0.02 | 51 | 0.00 | 0.00 | 3 | 0.00 | 0.00 | 41 | 0.06 | 0.11 | 60 | 0.00 | 0.00 | 1 |
| 40 | 0.00 | 0.00 | 33 | 0.05 | 0.08 | 34 | 0.00 | 0.00 | 0 | 0.00 | 0.00 | 39 | 0.00 | 0.00 | 51 | 0.00 | 0.00 | 1 |
| 44 | 0.00 | 0.00 | 27 | 0.00 | 0.00 | 20 | 0.00 | 0.00 | 0 | 0.00 | 0.00 | 43 | 0.04 | 0.06 | 33 | 0.00 | 0.00 | 1 |
| 48 | 0.00 | 0.00 | 22 | 0.00 | 0.00 | 14 | 0.00 | 0.00 | 0 | 0.00 | 0.00 | 41 | 0.08 | 0.14 | 25 | 0.00 | 0.00 | 0 |
| 52 | 0.00 | 0.00 | 29 | 0.00 | 0.00 | 11 | 0.00 | 0.00 | 0 | 0.03 | 0.04 | 23 | 0.00 | 0.00 | 14 | 0.00 | 0.00 | 0 |
| 56 | 0.00 | 0.00 | 20 | 0.00 | 0.00 | 11 | 0.00 | 0.00 | 0 | 0.00 | 0.00 | 20 | 0.00 | 0.00 | 11 | 0.00 | 0.00 | 0 |
| 60 | 0.05 | 0.08 | 17 | 0.00 | 0.00 | 6 | 0.00 | 0.00 | 0 | 0.00 | 0.00 | 21 | 0.00 | 0.00 | 10 | 0.00 | 0.00 | 0 |
| 64 | 0.00 | 0.00 | 13 | 0.00 | 0.00 | 2 | 0.00 | 0.00 | 0 | 0.00 | 0.00 | 10 | 0.00 | 0.00 | 4 | 0.00 | 0.00 | 0 |
| 68 | 0.00 | 0.00 | 6 | 0.00 | 0.00 | 1 | 0.00 | 0.00 | 0 | 0.08 | 0.14 | 15 | 0.00 | 0.00 | 3 | 0.00 | 0.00 | 0 |
| 72 | 0.00 | 0.00 | 14 | 0.00 | 0.00 | 2 | 0.00 | 0.00 | 0 | 0.00 | 0.00 | 11 | 0.00 | 0.00 | 3 | 0.00 | 0.00 | 0 |
| 76 | 0.00 | 0.00 | 5 | 0.00 | 0.00 | 2 | 0.00 | 0.00 | 0 | 0.00 | 0.00 | 8 | 0.00 | 0.00 | 1 | 0.00 | 0.00 | 0 |
| 80 | 0.00 | 0.00 | 16 | 0.00 | 0.00 | 1 | 0.00 | 0.00 | 0 | 0.00 | 0.00 | 5 | 0.00 | 0.00 | 0 | 0.00 | 0.00 | 0 |
| 84 | 0.00 | 0.00 | 9 | 0.00 | 0.00 | 1 | 0.00 | 0.00 | 0 | 0.00 | 0.00 | 3 | 0.00 | 0.00 | 1 | 0.00 | 0.00 | 0 |
| 88 | 0.00 | 0.00 | 7 | 0.00 | 0.00 | 0 | 0.00 | 0.00 | 0 | 0.00 | 0.00 | 2 | 0.00 | 0.00 | 2 | 0.00 | 0.00 | 0 |
| 92 | 0.00 | 0.00 | 2 | 0.00 | 0.00 | 0 | 0.00 | 0.00 | 0 | 0.00 | 0.00 | 1 | 0.00 | 0.00 | 0 | 0.00 | 0.00 | 0 |
| 96 | 0.00 | 0.00 | 4 | 0.00 | 0.00 | 1 | 0.00 | 0.00 | 0 | 0.00 | 0.00 | 3 | 0.00 | 0.00 | 0 | 0.00 | 0.00 | 0 |
| 100 | 0.00 | 0.00 | 7 | 0.00 | 0.00 | 0 | 0.00 | 0.00 | 0 | 0.00 | 0.00 | 1 | 0.00 | 0.00 | 0 | 0.00 | 0.00 | 0 |
| 104 | 0.00 | 0.00 | 1 | 0.00 | 0.00 | 0 | 0.00 | 0.00 | 0 | 0.00 | 0.00 | 0 | 0.00 | 0.00 | 0 | 0.00 | 0.00 | 0 |
| 108 | 0.00 | 0.00 | 5 | 0.00 | 0.00 | 0 | 0.00 | 0.00 | 0 | 0.00 | 0.00 | 0 | 0.00 | 0.00 | 0 | 0.00 | 0.00 | 0 |
| 112 | 0.00 | 0.00 | 0 | 0.00 | 0.00 | 0 | 0.00 | 0.00 | 0 | 0.00 | 0.00 | 0 | 0.00 | 0.00 | 0 | 0.00 | 0.00 | 0 |
| 116 | 0.00 | 0.00 | 1 | 0.00 | 0.00 | 0 | 0.00 | 0.00 | 0 | 0.00 | 0.00 | 0 | 0.00 | 0.00 | 0 | 0.00 | 0.00 | 0 |
| 120 | 0.00 | 0.00 | 0 | 0.00 | 0.00 | 0 | 0.00 | 0.00 | 0 | 0.00 | 0.00 | 0 | 0.00 | 0.00 | 0 | 0.00 | 0.00 | 0 |
| 124 | 0.00 | 0.00 | 0 | 0.00 | 0.00 | 0 | 0.00 | 0.00 | 0 | 0.00 | 0.00 | 0 | 0.00 | 0.00 | 0 | 0.00 | 0.00 | 0 |
| 128 | 0.00 | 0.00 | 2 | 0.00 | 0.00 | 0 | 0.00 | 0.00 | 0 | 0.00 | 0.00 | 0 | 0.00 | 0.00 | 0 | 0.00 | 0.00 | 0 |
| 132 | 0.00 | 0.00 | 1 | 0.00 | 0.00 | 0 | 0.00 | 0.00 | 0 | 0.00 | 0.00 | 0 | 0.00 | 0.00 | 0 | 0.00 | 0.00 | 0 |
| 136 | 0.00 | 0.00 | 0 | 0.00 | 0.00 | 0 | 0.00 | 0.00 | 0 | 0.00 | 0.00 | 0 | 0.00 | 0.00 | 0 | 0.00 | 0.00 | 0 |
| 140 | 0.00 | 0.00 | 1 | 0.00 | 0.00 | 0 | 0.00 | 0.00 | 0 | 0.00 | 0.00 | 0 | 0.00 | 0.00 | 0 | 0.00 | 0.00 | 0 |
| 144 | 0.00 | 0.00 | 0 | 0.00 | 0.00 | 0 | 0.00 | 0.00 | 0 | 0.00 | 0.00 | 0 | 0.00 | 0.00 | 0 | 0.00 | 0.00 | 0 |
| 148 | 0.00 | 0.00 | 1 | 0.00 | 0.00 | 0 | 0.00 | 0.00 | 0 | 0.00 | 0.00 | 0 | 0.00 | 0.00 | 0 | 0.00 | 0.00 | 0 |
| 152 | 0.00 | 0.00 | 1 | 0.00 | 0.00 | 0 | 0.00 | 0.00 | 0 | 0.00 | 0.00 | 0 | 0.00 | 0.00 | 0 | 0.00 | 0.00 | 0 |
| 156 | 0.00 | 0.00 | 0 | 0.00 | 0.00 | 0 | 0.00 | 0.00 | 0 | 0.00 | 0.00 | 0 | 0.00 | 0.00 | 0 | 0.00 | 0.00 | 0 |
| 160 | 0.00 | 0.00 | 0 | 0.00 | 0.00 | 0 | 0.00 | 0.00 | 0 | 0.00 | 0.00 | 0 | 0.00 | 0.00 | 0 | 0.00 | 0.00 | 0 |
| 164 | 0.00 | 0.00 | 1 | 0.00 | 0.00 | 0 | 0.00 | 0.00 | 0 | 0.00 | 0.00 | 0 | 0.00 | 0.00 | 0 | 0.00 | 0.00 | 0 |

Table 1.17: Average stromule frequency values (SF) and corresponding standard deviation (SD) as well as number of in total scored plastids (n) for each distance zone from the nucleus ($\mu\text{m}$) for wit1\_wit2 and 4ko lines. Values used to plot Figure 5C and D. w12 = wit1\_wit2, 4ko = 4ko mutant lines, L = large cells, M = medium cells, S = small cells

##### 1.2.14 Supplemental Information 14 - Zone Measurements - Stats2

**Supported Figure: 4A-D** This table provides the per-replicate stromule frequency values used to assess reproducibility of the SPZ measurements shown in Figure 4A–D. For each genotype (wt, xi-i, wit1/2, and 4ko) and cell size class (large [L], medium [M], and small [S]), stromule frequency is given for the 4 $\mu\text{m}$, 8 $\mu\text{m}$, and 12 $\mu\text{m}$ distance zones for each of the three independent transgenic lines analyzed. These values illustrate the degree of biological variability across lines and support the averaged data presented in Supplemental Information 13 and Figure 4A–D.

| wt |  |  | xi-i |  |  | w12 |  |  | 4ko |  |  |
| --- | --- | --- | --- | --- | --- | --- | --- | --- | --- | --- | --- |
| L |  |  | L |  |  | L |  |  | L |  |  |
| 4$\mu\text{m}$ | 8$\mu\text{m}$ | 12$\mu\text{m}$ | 4$\mu\text{m}$ | 8$\mu\text{m}$ | 12$\mu\text{m}$ | 4$\mu\text{m}$ | 8$\mu\text{m}$ | 12$\mu\text{m}$ | 4$\mu\text{m}$ | 8$\mu\text{m}$ | 12$\mu\text{m}$ |
| 0.35 | 0.11 | 0.07 | 0.28 | 0.29 | 0.20 | 0.12 | 0.40 | 0.25 | 0.02 | 0.00 | 0.00 |
| 0.45 | 0.20 | 0.09 | 0.00 | 0.00 | 0.00 | 0.26 | 0.00 | 0.14 | 0.00 | 0.00 | 0.00 |
| 0.37 | 0.47 | 0.04 | 0.21 | 0.33 | 0.00 | 0.21 | 0.00 | 0.50 | 0.11 | 0.00 | 0.06 |
| 0.39 | 0.33 | 0.14 |  |  |  |  |  |  |  |  |  |
| 0.33 | 0.19 | 0.09 |  |  |  |  |  |  |  |  |  |
| 0.40 | 0.13 | 0.05 |  |  |  |  |  |  |  |  |  |
| wt |  |  | xi-i |  |  | w12 |  |  | 4ko |  |  |
| M |  |  | M |  |  | M |  |  | M |  |  |
| 4$\mu\text{m}$ | 8$\mu\text{m}$ | 12$\mu\text{m}$ | 4$\mu\text{m}$ | 8$\mu\text{m}$ | 12$\mu\text{m}$ | 4$\mu\text{m}$ | 8$\mu\text{m}$ | 12$\mu\text{m}$ | 4$\mu\text{m}$ | 8$\mu\text{m}$ | 12$\mu\text{m}$ |
| 0.44 | 0.29 | 0.05 | 0.03 | 0.00 | 0.00 | 0.06 | 0.06 | 0.04 | 0.00 | 0.03 | 0.00 |
| 0.15 | 0.21 | 0.03 | 0.01 | 0.12 | 0.07 | 0.02 | 0.08 | 0.03 | 0.02 | 0.00 | 0.04 |
| 0.18 | 0.21 | 0.03 | 0.08 | 0.00 | 0.00 | 0.00 | 0.00 | 0.00 | 0.01 | 0.00 | 0.00 |
| 0.20 | 0.17 | 0.00 |  |  |  |  |  |  |  |  |  |
| 0.17 | 0.06 | 0.00 |  |  |  |  |  |  |  |  |  |
| 0.13 | 0.09 | 0.00 |  |  |  |  |  |  |  |  |  |
| wt |  |  | xi-i |  |  | w12 |  |  | 4ko |  |  |
| S |  |  | S |  |  | S |  |  | S |  |  |
| 4$\mu\text{m}$ | 8$\mu\text{m}$ | 12$\mu\text{m}$ | 4$\mu\text{m}$ | 8$\mu\text{m}$ | 12$\mu\text{m}$ | 4$\mu\text{m}$ | 8$\mu\text{m}$ | 12$\mu\text{m}$ | 4$\mu\text{m}$ | 8$\mu\text{m}$ | 12$\mu\text{m}$ |
| 0.06 | 0.00 | 0.00 | 0.01 | 0.00 | 0.00 | 0.00 | 0.07 | 0.00 | 0.00 | 0.00 | 0.00 |
| 0.09 | 0.09 | 0.00 | 0.01 | 0.00 | 0.00 | 0.01 | 0.00 | 0.00 | 0.00 | 0.00 | 0.00 |
| 0.02 | 0.00 | 0.00 | 0.00 | n<3 | 0.00 | 0.00 | 0.02 | 0.00 | 0.01 | 0.00 | 0.00 |
| 0.09 | 0.07 | 0.00 |  |  |  |  |  |  |  |  |  |
| 0.01 | 0.00 | 0.00 |  |  |  |  |  |  |  |  |  |
| 0.06 | 0.00 | 0.00 |  |  |  |  |  |  |  |  |  |

Table 1.18: Average stromule frequency values of individual transgenic lines for wildtype and the different mutant backgrounds in the first 3 distance zones. Values used to calculate averages in plot Figure 5C and D. wt = wildtype, xi-i = xi-i mutant, w12 = wit1\_wit2, 4ko = 4ko mutant lines, L = large cells, M = medium cells, S = small cells

##### 1.2.15 Supplemental Information 15 - Speed vs. SF% Correlation Analysis

**Supported Figure: 5A-C** This table provides the numerical values used to test the correlation between nucleus speed and stromule frequency (SF%) shown in Figure 5A–C. For each genotype (wt, xi-i, wit1/2, and 4ko) and cell size class (large, medium, and small), the average nucleus speed ($\mu\text{m}/\text{h}$) and the mean SF% within the 4 $\mu\text{m}$ zone, 8 $\mu\text{m}$ zone, and beyond 8 $\mu\text{m}$ are listed. These twelve data points, one per genotype–cell size combination, form the input for the linear regression analysis shown in Figure 5A–C and Supplemental Information 16.

| speed in $\mu\text{m}/\text{h}$ | SF% 4$\mu\text{m}$ | SF% 8$\mu\text{m}$ | SF% >8$\mu\text{m}$ | line/cell size |
| --- | --- | --- | --- | --- |
| 30.0000 | 38.0000 | 24.0000 | 3.0000 | wt lrg |
| 24.0000 | 21.0000 | 17.0000 | 2.0000 | wt med |
| 17.0000 | 6.0000 | 3.0000 | 1.0000 | wt small |
| 18.0000 | 17.0000 | 21.0000 | 1.0000 | *xi-i* lrg |
| 11.0000 | 4.0000 | 4.0000 | 1.0000 | *xi-i* med |
| 8.0000 | 1.0000 | 0.0000 | 0.0000 | *xi-i* small |
| 22.0000 | 20.0000 | 13.0000 | 2.0000 | *wit1/2* lrg |
| 12.0000 | 3.0000 | 5.0000 | 2.0000 | *wit1/2* med |
| 9.0000 | 0.0000 | 2.0000 | 0.0000 | *wit1/2* small |
| 6.0000 | 4.0000 | 0.0000 | 1.0000 | *4ko* lrg |
| 6.0000 | 1.0000 | 1.0000 | 2.0000 | *4ko* med |
| 4.0000 | 0.0000 | 0.0000 | 1.0000 | *4ko* small |

Table 1.19: numerical values used to test the correlation of nucleus speed and stromule frequency (SF %); lrg = large, med = medium

##### 1.2.16 Supplemental Information 16 - Speed vs. SF% Correlation Analysis - Graphs

**Supported Figure: 5A-C** This section presents the graphical output of the correlation analysis between nuclear movement speed and stromule frequency (SF%) introduced in Supplemental Information 15. For each combination of distance zone (4 $\mu\text{m}$, 8 $\mu\text{m}$, and >8 $\mu\text{m}$) and cell size (small, medium, and large), the average nuclear speed ($\mu\text{m}/\text{h}$) was plotted against the corresponding SF% value, combining data from all genetic backgrounds (wt, *xi-i*, *wit1\_wit2*, and *4ko*) to test for a correlation between the two parameters (Figure 1.4). These graphs form the basis for the correlation shown in main manuscript Figure 5A–C, where data are separated into within-SPZ (4 $\mu\text{m}$ and 8 $\mu\text{m}$ zones) and outside-SPZ (>8 $\mu\text{m}$) categories.

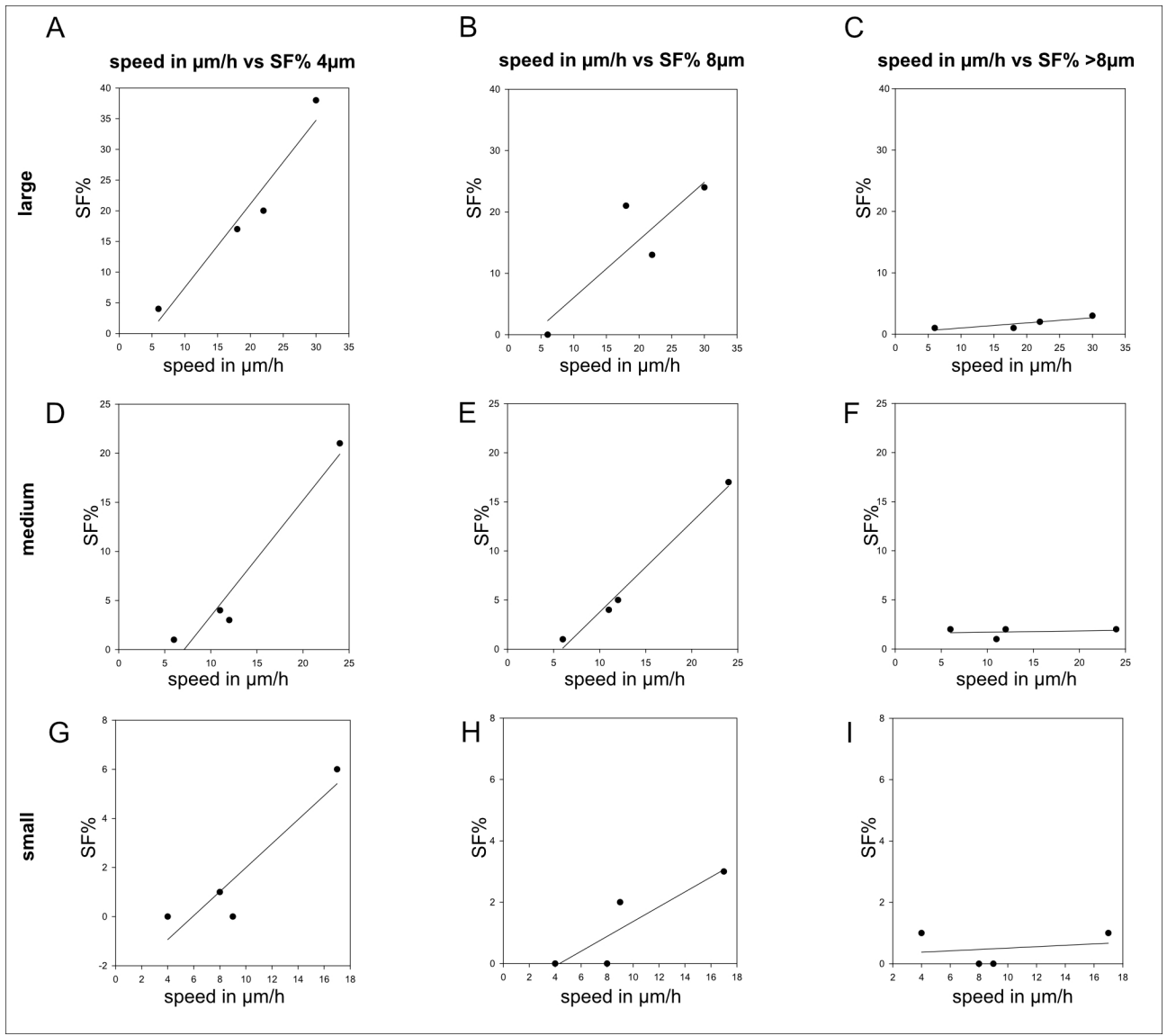

Figure 1.4: Scatter plots of stromule frequency (SF%) vs. average nucleus speed (μm/h) to test for correlation of both. One scatter plot combines the nucleus speed and stromule frequency of different genetic backgrounds (wt, wit1/2, xi-i and 4ko) in a certain distance zone (4μm, 8μm and >8μm) and cell size (large, medium and small).

##### 1.2.17 Supplemental Information 17 - Quantification of Stromule Forming Events

**Supported Figure: 6F-H** This section summarizes the data and statistical approach underlying the quantification of stromule formation events shown in Figure 6F–H. Stromule formation events identified in time-lapse recordings were categorized as cor+ (correlation positive), when stromule formation coincided with nuclear movement in both direction and distance, or cor– (correlation negative), when stromule formation occurred independently of nuclear movement, including events at a stationary nucleus or events where direction and distance did not match nuclear displacement (see main manuscript Figure 6E1–E4 for category definitions). Because the number of plastids analyzed per recording varied both within and between genotypes, raw event counts were normalized to a standard reference of 30 plastids per sample before calculating temporal event frequencies (events per frame), allowing unbiased comparison of cor+ and cor– event frequencies across genetic backgrounds.

###### Statistical Normalization Approach for Cross-Genotype Comparison

**Justification for Normalization to a Standard Plastid Number** To enable meaningful comparison of stromule forming event frequencies across genotypes with varying plastid numbers, we implemented a normalization strategy that standardizes all observations to a standard plastid number per sample. This approach was necessary because the raw plastid counts differed both between genotypes (ranging from mean 28.4 in 4KO to 32.9 in wild-type) and within genotypes (overall range: 13-59 plastids).

**Methodological Approach** For each time-lapse recording, we calculated the rate of cor+ and cor– events per plastid (events/n pl), then scaled this rate to a standard reference of 30 plastids before calculating the temporal frequency (events per frame). This normalization value was selected based on statistical analysis showing that 30 represents both the median (30.0) and approximate mean (30.4) of plastid counts across all samples, minimizing extrapolation bias while maintaining biological relevance.

**Values of counts** In the table below you find the RAW counts of stromule forming events.

Table 1.20: Per-recording quantification of stromule formation events used to generate Figure 6F–H. For each time-lapse movie, the number of cor+ (correlation positive, stromule formation coinciding with nuclear movement) and cor– (correlation negative, asynchronous stromule formation) events is given, together with the number of nucleus-associated plastids analyzed, the number of imaged frames, the genetic background (wt = wildtype, xii = xi-i, wit12 = wit1\_wit2, 4KO = 4ko), and the corresponding source movie. These values form the basis for the normalized event frequencies shown in Figure 6F–H.

| cor+ | cor- | nucleus associated plastids | number of frames | genetic background | image |
| --- | --- | --- | --- | --- | --- |
| 55 | 12 | 47 | 35 | wt | ID_07_WT_2016-03-29_WT10_pn_movie01_MaxProject |
| 26 | 5 | 36 | 36 | wt | ID_08_WT_2016-03-29_WT10_pn_movie02_MaxProject |
| 40 | 20 | 30 | 23 | wt | ID_13_WT_2016-03-29_WT10_pn_movie07_MaxProject |
| 45 | 14 | 21 | 40 | wt | ID_09_WT_2016-03-29_WT10_pn_movie03_MaxProject |
| 20 | 7 | 19 | 33 | wt | ID_10_WT_2016-03-29_WT10_pn_movie04_MaxProject |
| 52 | 9 | 59 | 36 | wt | ID_12_WT_2016-03-29_WT10_pn_movie06_MaxProject |
| 10 | 8 | 27 | 40 | wt | ID_14_WT_2016-03-29_WT10_pn_movie09_MaxProject |
| 19 | 4 | 24 | 35 | wt | ID_01_2018-05-25_WT-pn-2-1-Pflanze1-movie01_Maximum intensity projection |
| 7 | 25 | 38 | 31 | xii | 2017-04-20_xi-i-pn-line4-plant2-video2_Maximum intensity projection |
| 3 | 13 | 29 | 31 | xii | 2017-04-20_xi-i-pn-line4-plant1-video3_Maximum intensity projection |
| 7 | 6 | 13 | 30 | xii | 2017-04-20_xi-i-pn-line4-plant2-video1a_and_b_together |
| 2 | 4 | 30 | 30 | xii | 2017-04-19_xi-i-pn-line4-plant4-video4_Maximum intensity projection |
| 20 | 32 | 32 | 32 | xii | 2017-04-18_xi-i-pn-line4-plant3-video2_Maximum intensity projection |
| 5 | 4 | 40 | 32 | xii | 2017-04-20_xi-i-pn-line4-plant1-video2_Maximum intensity projection |

| genetic background | cor+ | cor- | nucleus associated plastids | sum of frames | total time imaged (frames*3min) |
| --- | --- | --- | --- | --- | --- |
| wt | 267 | 79 | 263 | 278 | 834 min |
| xii | 82 | 116 | 271 | 277 | 831 min |
| wit12 | 99 | 70 | 257 | 302 | 906 min |
| 4KO | 7 | 7 | 284 | 277 | 831 min |

Table 1.21: Summary of cor+ and cor- event totals per genetic background, aggregated across all recordings listed in table 1.20. For each genotype, the total number of cor+ and cor- events, the total number of nucleus-associated plastids analyzed, the summed number of frames, and the corresponding total imaging time (frames × 3 min) are given. wt = wildtype, xii = xi-i, wit12 = wit1\_wit2, 4KO = 4ko.

|  |  |  |  |  |  |
| --- | --- | --- | --- | --- | --- |
| 18 | 12 | 29 | 30 | xii | 2017-04-20_xi-i-pn-line4-plant2-video3_Maximum intensity projection |
| 20 | 17 | 31 | 30 | xii | 2017-04-18_xi-i-pn-line4-plant3-video3_Maximum intensity projection |
| 0 | 3 | 29 | 31 | xii | 2017-04-19_xi-i-pn-line4-plant4-video3_Maximum intensity projection |
| 16 | 8 | 25 | 31 | wit12 | wit1und2-pn-T1-Line08_movie01a+b_Maximum intensity projection |
| 3 | 3 | 23 | 34 | wit12 | wit1und2-pn-T1-Line08_movie02_Maximum intensity projection |
| 10 | 1 | 37 | 37 | wit12 | wit1und2-pn-T1-Line08_movie03a+3b_Maximum intensity projection |
| 6 | 3 | 32 | 33 | wit12 | wit1und2-pn-T1-Line08_movie04_Maximum intensity projection-1 |
| 15 | 7 | 32 | 32 | wit12 | wit1und2-pn-T1-Line08_movie05a+5b_Maximum intensity projection |
| 17 | 23 | 39 | 38 | wit12 | wit1und2-pn-T1-Line08_movie06_Maximum intensity projection-1 |
| 11 | 13 | 15 | 39 | wit12 | wit1und2-pn-T1-Line08_movie07a+b_Maximum intensity projection |
| 5 | 1 | 22 | 28 | wit12 | wit1und2-pn-T1-Line08_movie08_Maximum intensity projection-1 |
| 16 | 11 | 32 | 30 | wit12 | wit1und2-pn-T1-Line08_movie09_Maximum intensity projection-1 |
| 3 | 0 | 20 | 36 | 4KO | 2017-05-02_4KO-pn_line01_plant01_video01_Maximum intensity projection |
| 0 | 0 | 26 | 37 | 4KO | 2017-05-02_4KO-pn_line01_plant01_video02_Maximum intensity projection-1 |
| 1 | 2 | 33 | 40 | 4KO | 2017-05-02_4KO-pn_line01_plant01_video03_Maximum intensity projection |
| 1 | 2 | 22 | 30 | 4KO | 2017-05-03_4KO-pn_line01_plant01_video05_Maximum intensity projection |
| 0 | 1 | 28 | 19 | 4KO | 2017-05-10_4KO-pn_line01_plant01_video06a_Maximum intensity projection |
| 1 | 0 | 27 | 19 | 4KO | 2017-05-10_4KO-pn_line01_plant04_video07b_Maximum intensity projection |
| 1 | 0 | 23 | 16 | 4KO | 2017-05-10_4KO-pn_line01_plant04_video08a_Maximum intensity projection |
| 0 | 1 | 29 | 29 | 4KO | 2017-05-10_4KO-pn_line01_plant04_video09_Maximum intensity projection |
| 0 | 1 | 40 | 15 | 4KO | 2017-05-10_4KO-pn_line01_plant01_video10_Maximum intensity projection |
| 0 | 0 | 36 | 36 | 4KO | 2017-05-15_4KO-pn_line01_plant01_video12_Maximum intensity projection |

**Aggregated data for stromule forming events** This is a representation of the aggregated counts and summary statistics for stromule forming events for each genetic background used.

| variable | genetic background | Bottom whisker end<br>(largest point $\geq Q1 - 1.5 \times IQR$) | First quartile<br>(Q1, 25th percentile) | median | Third quartile<br>(Q3, 75th percentile) | Highest point<br>$\leq Q3 + 1.5 \times IQR$ |
| --- | --- | --- | --- | --- | --- | --- |
| cor+ | wt | 0.28 | 0.66 | 0.85 | 1.15 | 1.74 |
| cor+ | xii | 0.00 | 0.10 | 0.18 | 0.59 | 0.65 |
| cor+ | wit12 | 0.12 | 0.22 | 0.34 | 0.50 | 0.62 |
| cor+ | 4KO | 0.00 | 0.00 | 0.01 | 0.06 | 0.13 |
| cor- | wt | 0.12 | 0.14 | 0.22 | 0.38 | 0.50 |
| cor- | xii | 0.09 | 0.13 | 0.43 | 0.55 | 0.94 |
| cor- | wit12 | 0.02 | 0.09 | 0.21 | 0.34 | 0.67 |
| cor- | 4KO | 0.00 | 0.00 | 0.02 | 0.05 | 0.09 |

Table 1.22: Summary statistics (quartiles and whisker boundaries) of the normalized cor+ and cor- event frequencies per genetic background, underlying the box plots shown in Figure 6F–H. wt = wildtype, xii = xi-i, wit12 = wit1\_wit2, 4KO = 4ko.

#### Statistical testing of cor- and cor+ within the genetic backgrounds for Figure 6H

**Step1: Test for normality** To compare cor+ and cor- event frequencies within each genetic background, the normality of the paired differences (cor+ minus cor-) was first assessed using the Shapiro-Wilk test using R and R-studio. For all four genetic backgrounds, the differences did not deviate significantly from a normal distribution (wt: $W = 0.961$, $p = 0.8229$; xi-i: $W = 0.93$, $p = 0.4828$; wit1\_wit2: $W = 0.927$, $p = 0.4538$; 4ko: $W = 0.876$, $p = 0.117$), supporting the use of a paired t-test for subsequent comparisons.

**Step2: t-Test to check for difference of cor- and cor+ events within the genetic backgrounds** Having confirmed that the paired differences were normally distributed, paired t-tests were performed to compare the normalized frequencies of cor+ and cor- events within each genetic background. In wildtype, cor+ events occurred significantly more frequently than cor- events (mean cor+ = 0.950, mean cor- = 0.316; $t = 5.796$, $df = 7$, $p = 0.000666$). In contrast, no significant difference between cor+ and cor- frequencies was observed in xi-i (mean cor+ = 0.317, mean cor- = 0.418; $t = -1.308$, $df = 8$, $p = 0.2272$), wit1\_wit2 (mean cor+ = 0.357, mean cor- = 0.251; $t = 2.098$, $df = 8$, $p = 0.06914$), or 4ko (mean cor+ = 0.033, mean cor- = 0.028; $t = 0.278$, $df = 9$, $p = 0.7873$).

These results show that the predominance of nuclear movement-coupled (cor+) stromule formation observed in wildtype is lost in all three mutant backgrounds. In xi-i and wit1\_wit2, where XI-K, XI-1, and XI-2 remain functional, cor- events persist at frequencies comparable to cor+ events, indicating that cor- events do not depend on nuclear movement itself but rather on the activity of these remaining myosins. In 4ko, however, both cor+ and cor- event frequencies collapse to near-zero levels. The complete loss of cor- in 4ko indicates that cor- events depend specifically on XI-K, XI-1, and/or XI-2, consistent with their persistence in xi-i and wit1\_wit2, where these myosins are still present. The parallel loss of cor+ in 4ko is consistent with our earlier observation that nuclear movement, although reduced in xi-i, is not abolished unless XI-K, XI-1, and XI-2 are also removed: the residual nuclear movement present in xi-i (driven by XI-K, XI-1, and XI-2 in the absence of XI-I) is sufficient to sustain cor+ events, whereas its near-complete loss in 4ko abolishes them. Taken together, these findings indicate that cor+ events depend on nuclear movement, which itself can be driven by XI-I or, in its absence, by XI-K, XI-1, and XI-2, while cor- events represent a mechanistically distinct, nucleus-movement-independent pathway that depends directly on XI-K, XI-1, and/or XI-2.

This points to two separable mechanisms of stromule formation near the nucleus: one coupled to nuclear movement (cor+), and one acting directly through myosin-actin dynamics independent of nuclear displacement (cor-).

**Step3: complementary Wilcoxon signed-rank test to t-Test** To verify the robustness of the paired comparisons of cor+ and cor- event frequencies within each genetic background, the non-parametric

| Genetic background | Mean cor+ | Mean cor- | t | df | p-value | Significance |
| --- | --- | --- | --- | --- | --- | --- |
| wt | 0.950 | 0.316 | 5.796 | 7 | 0.000666 | *** |
| xi-i | 0.317 | 0.418 | -1.308 | 8 | 0.2272 | ns |
| wit1_wit2 | 0.357 | 0.251 | 2.098 | 8 | 0.06914 | ns |
| 4KO | 0.033 | 0.028 | 0.278 | 9 | 0.7873 | ns |

**Table 1.23:** Comparison of cor+ and cor- stromule formation event frequencies within each genetic background. Normalized event frequencies (events per frame, scaled to 30 plastids) for cor+ (correlation positive, stromule formation coinciding with nuclear movement) and cor- (correlation negative, asynchronous stromule formation) events are shown for wildtype (wt), xi-i, wit1\_wit2, and 4ko. Paired t-tests were performed within each genotype to compare cor+ and cor- frequencies. In wildtype, cor+ events occurred significantly more frequently than cor- events ($t = 5.796$, $df = 7$, $p = 0.00067$, \*\*\*), whereas no significant difference between cor+ and cor- frequencies was found in xi-i ($t = -1.308$, $df = 8$, $p = 0.227$), wit1\_wit2 ($t = 2.098$, $df = 8$, $p = 0.069$), or 4ko ($t = 0.278$, $df = 9$, $p = 0.787$). ns = not significant; \* = $p < 0.05$; \*\* = $p < 0.01$; \*\*\* = $p < 0.001$.

| Genetic background | Mean (Median) |  | Paired t-test |  |  | Wilcoxon signed-rank |
| --- | --- | --- | --- | --- | --- | --- |
|  | cor+ | cor- | t | df | p-value | V |
| wt | 0.950 (0.846) | 0.316 (0.221) | 5.796 | 7 | 0.000666 *** | 36 |
| xi-i | 0.317 (0.178) | 0.418 (0.434) | -1.308 | 8 | 0.2272 ns | 14 |
| wit1_wit2 | 0.357 (0.344) | 0.251 (0.205) | 2.098 | 8 | 0.06914 ns | 31 |
| 4KO | 0.033 (0.011) | 0.028 (0.018) | 0.278 | 9 | 0.7873 ns | 21 |

**Table 1.24:** Within-genotype comparison of cor+ and cor- event frequencies (paired t-test and Wilcoxon signed-rank test). ns = no significant difference between cor+ and cor- events; \* = $p < 0.05$; \*\* = $p < 0.01$; \*\*\* = $p < 0.001$. Values for cor+ and cor- are given as mean (median), normalized to 30 plastids per sample. Both a paired t-test and a Wilcoxon signed-rank test (non-parametric) were performed for each genetic background; the two approaches agreed on the presence or absence of a significant difference for all genotypes, though the resulting significance level for wildtype differs (\*\*\* for the t-test vs. \* for the Wilcoxon test) due to the difference in p-values between the two tests ($p = 0.000666$ vs. $p = 0.0143$).

Wilcoxon signed-rank test was performed as a complement to the paired t-test. The results were consistent between the two approaches for all genotypes. In wildtype, cor+ events occurred significantly more frequently than cor- events ($V = 36$, $p = 0.0143$), confirming the result of the paired t-test ($p = 0.00067$). No significant difference between cor+ and cor- frequencies was found in xi-i ($V = 14$, $p = 0.343$), wit1\_wit2 ($V = 31$, $p = 0.080$), or 4ko ($V = 21$, $p = 0.726$), again matching the corresponding t-test results ($p = 0.227$, $0.069$, and $0.787$, respectively). The agreement between the parametric and non-parametric tests indicates that the observed differences (and lack thereof) are robust to deviations from normality in the underlying data.

#### Statistical testing of cor- and cor+ between the genetic backgrounds for Figure 6F and G

To test for differences in cor+ and cor- event frequencies between genetic backgrounds (main manuscript Figure 6F, G), data were first normalized to a standard of 30 plastids per sample to account for variation in plastid numbers between recordings (see 1.2.17). One-way ANOVA was initially considered for between-genotype comparisons; however, assumption checks revealed violations for both variables: for cor+, Levene's test indicated significantly unequal variances across genotypes ($F = 5.686$, $p = 0.00308$), while for cor-, Shapiro-Wilk tests on the model residuals indicated significant deviation from normality ($W = 0.922$, $p = 0.0143$). These violations likely reflect the heavily zero-inflated distribution of cor+ and cor- values in the 4ko mutant (see table 1.20 on page 33). The non-parametric Kruskal-Wallis test was therefore used as the primary omnibus test for between-genotype differences for both variables, followed by pairwise Wilcoxon rank-sum tests with Holm correction for multiple comparisons as post-hoc. For cor+, the Kruskal-Wallis test revealed a significant overall difference between genotypes ($\chi^2 = 24.152$, $df = 3$, $p = 2.32 \times 10^{-5}$). Post-hoc comparisons showed that cor+ frequency was significantly reduced in xi-i ($p = 0.0099$), wit1\_wit2 ($p = 0.0099$), and 4ko ($p = 0.0020$) compared to wildtype. Furthermore, 4ko showed significantly lower

| Comparison | Holm-adjusted p-value | Significance |
| --- | --- | --- |
| wt vs. xi-i | 0.0099 | * |
| wt vs. wit1_wit2 | 0.0099 | * |
| wt vs. 4ko | 0.0020 | ** |
| xi-i vs. wit1_wit2 | 0.7304 | ns |
| xi-i vs. 4ko | 0.0124 | * |
| wit1_wit2 vs. 4ko | 0.0020 | ** |

**Table 1.25:** Pairwise comparisons of cor+ event frequency between genetic backgrounds for use in main manuscript Figure 6F (Wilcoxon rank-sum test, Holm-adjusted p-values); ns = not significant; \* = $p < 0.05$; \*\* = $p < 0.01$. P-values were computed using the Wilcoxon rank-sum test with normal approximation (exact p-values could not be computed due to ties in the data) and adjusted for multiple comparisons using the Holm method.

| Comparison | Holm-adjusted p-value | Significance |
| --- | --- | --- |
| wt vs. xi-i | 0.8469 | ns |
| wt vs. wit1_wit2 | 0.8469 | ns |
| wt vs. 4ko | 0.0020 | ** |
| xi-i vs. wit1_wit2 | 0.5708 | ns |
| xi-i vs. 4ko | 0.0015 | ** |
| wit1_wit2 vs. 4ko | 0.0138 | * |

**Table 1.26:** Pairwise comparisons of cor- event frequency between genetic backgrounds for use in main manuscript Figure 6G (Wilcoxon rank-sum test, Holm-adjusted p-values); ns = not significant; \* = $p < 0.05$; \*\* = $p < 0.01$. P-values were computed using the Wilcoxon rank-sum test with normal approximation (exact p-values could not be computed due to ties in the data) and adjusted for multiple comparisons using the Holm method.

cor+ frequency than both xi-i ($p = 0.0124$) and wit1\_wit2 ($p = 0.0020$), while xi-i and wit1\_wit2 did not differ from each other ($p = 0.730$). For cor-, the Kruskal-Wallis test likewise revealed a significant overall difference between genotypes ($\chi^2 = 19.883$, $df = 3$, $p = 0.00018$). Post-hoc comparisons showed that 4ko had significantly lower cor- frequency than wildtype ($p = 0.0020$), xi-i ($p = 0.0015$), and wit1\_wit2 ($p = 0.0138$), whereas wildtype, xi-i, and wit1\_wit2 did not differ from one another (all $p \geq 0.571$). These results indicate that cor+ event frequency is progressively reduced as components of the myosin XI/WIT1-WIT2 nuclear movement machinery are removed, with the most severe reduction occurring in 4ko, consistent with cor+ events reflecting nucleus-movement-driven stromule formation. The selective reduction of cor- frequency only in 4ko, but not in xi-i or wit1\_wit2, supports the conclusion that this nucleus-movement-independent subpopulation of stromules depends specifically on XI-K, XI-1, and XI-2.

##### 1.2.18 Supplemental Information 18 - Nucleus Tracks

**Supported Figure: 7C-F** The following figures show the tracks of all the nuclei considered to test if nucleus movement can explain the measured extend of the stromule promoting zone in wildtype plants. During the image capture and tracking, some nuclei moved out of the field of view, resulting in aberrant tracks, which became visible in the graphs of the tracks. Respective nuclei have been removed from the analysis. If such tracks are still represented in the figures, figure legends disclose those tracks as 'removed from analysis'.

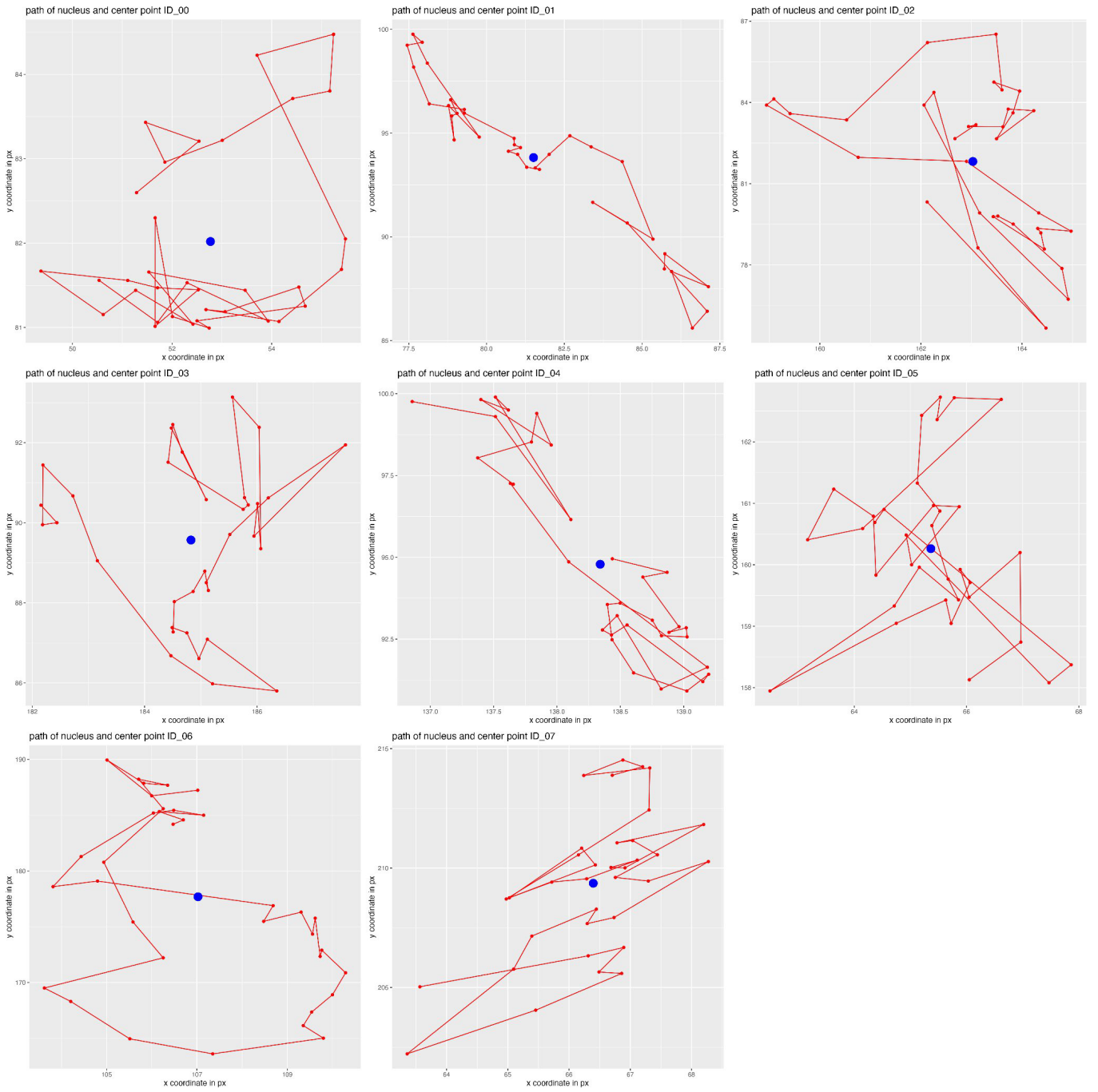

Figure 1.5: Nuclei tracks (red) with calculated center point (blue) for movie\_ID01

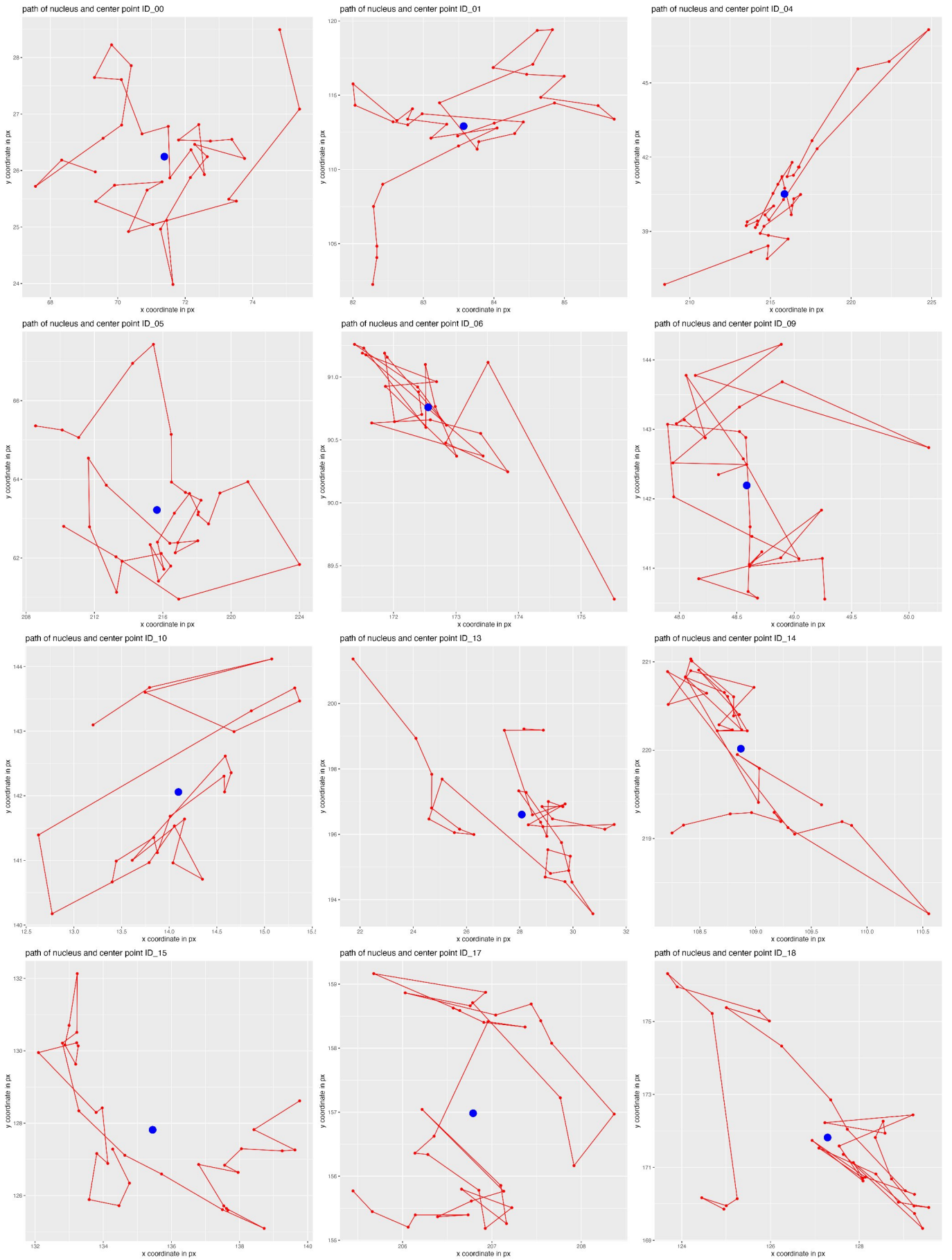

Figure 1.6: Nuclei tracks (red) with calculated center point (blue) for movie\_ID02

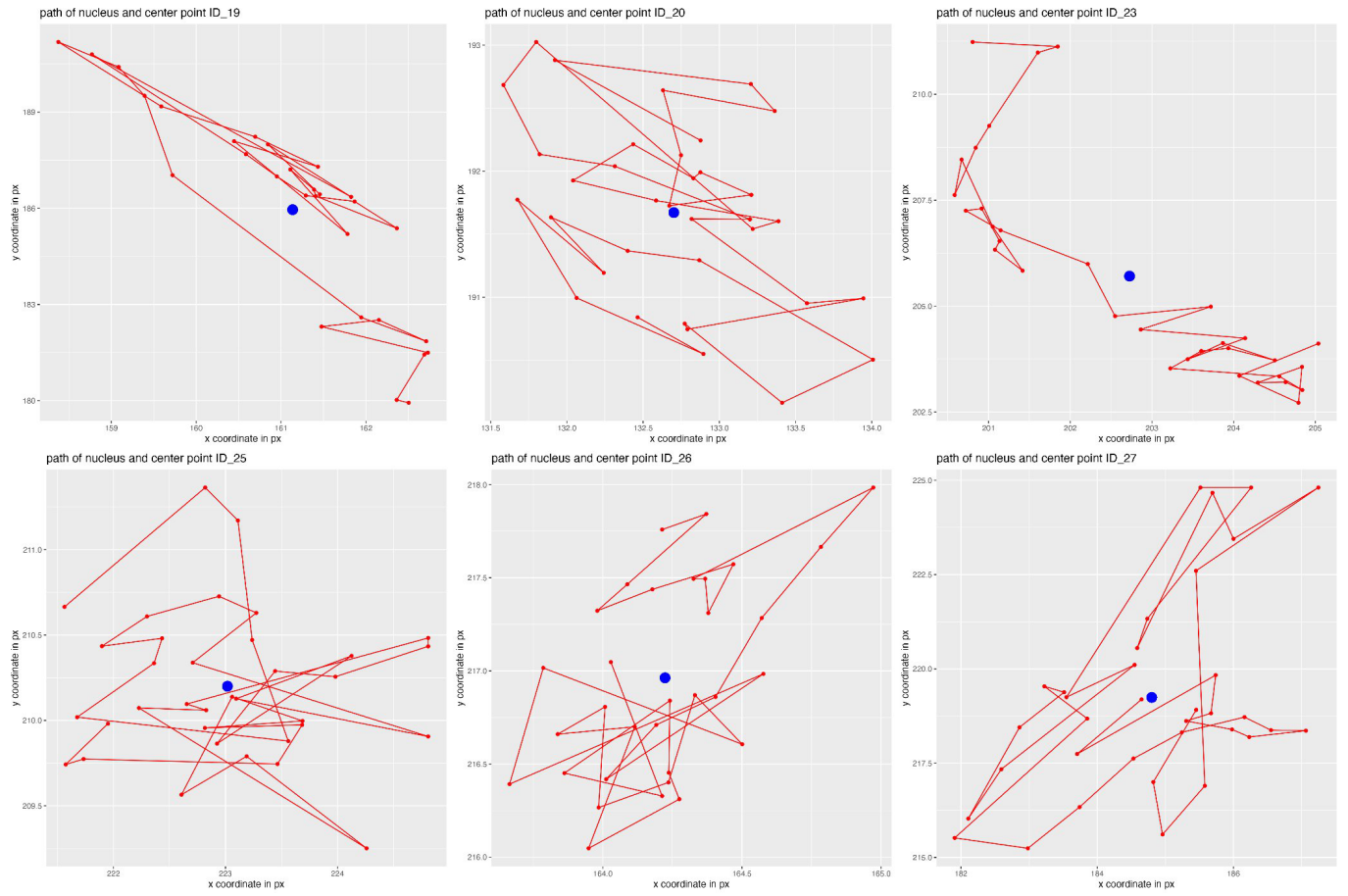

Figure 1.7: Nuclei tracks (red) with calculated center point (blue) for movie\_ID02

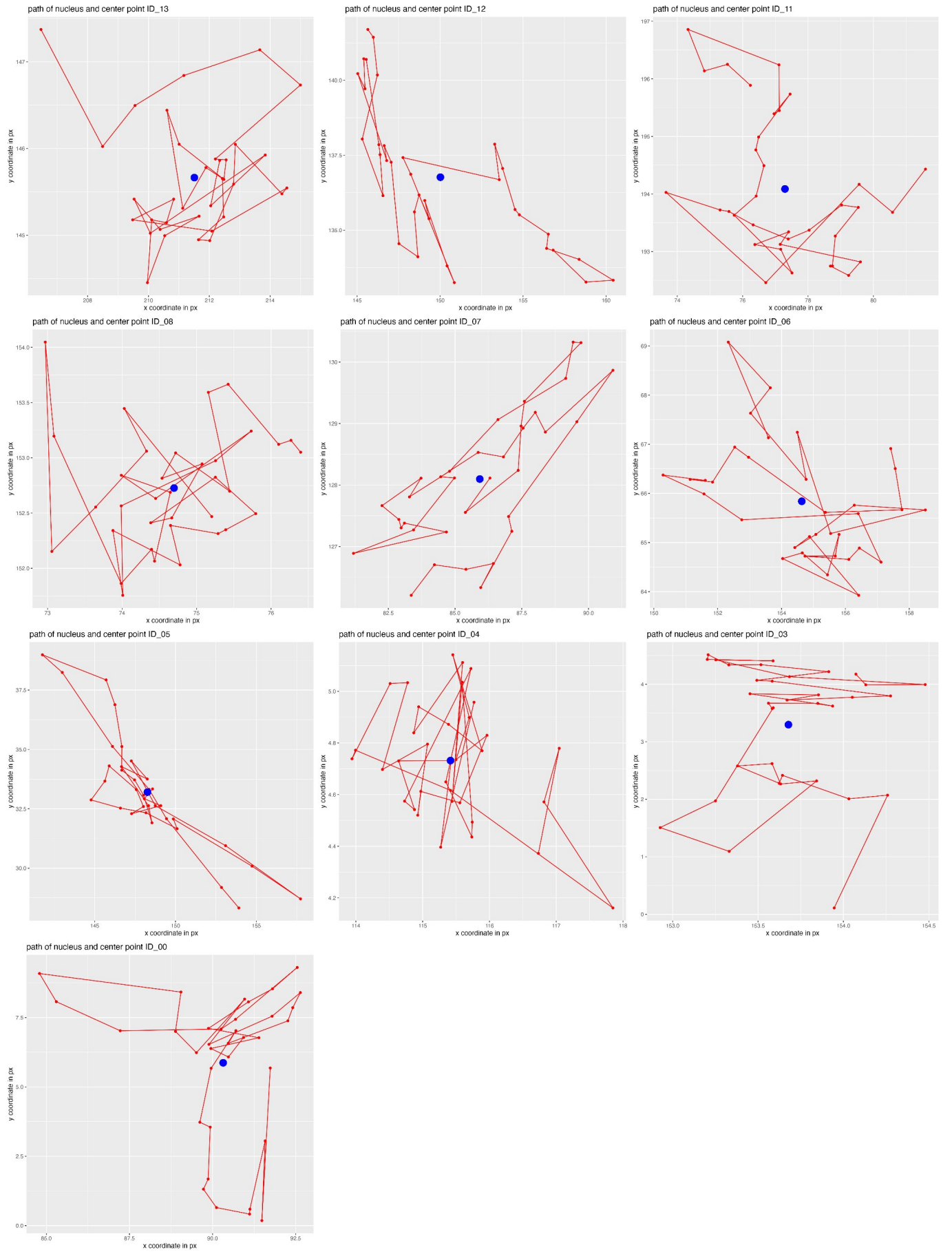

Figure 1.8: Nuclei tracks (red) with calculated center point (blue) for movie\_ID03

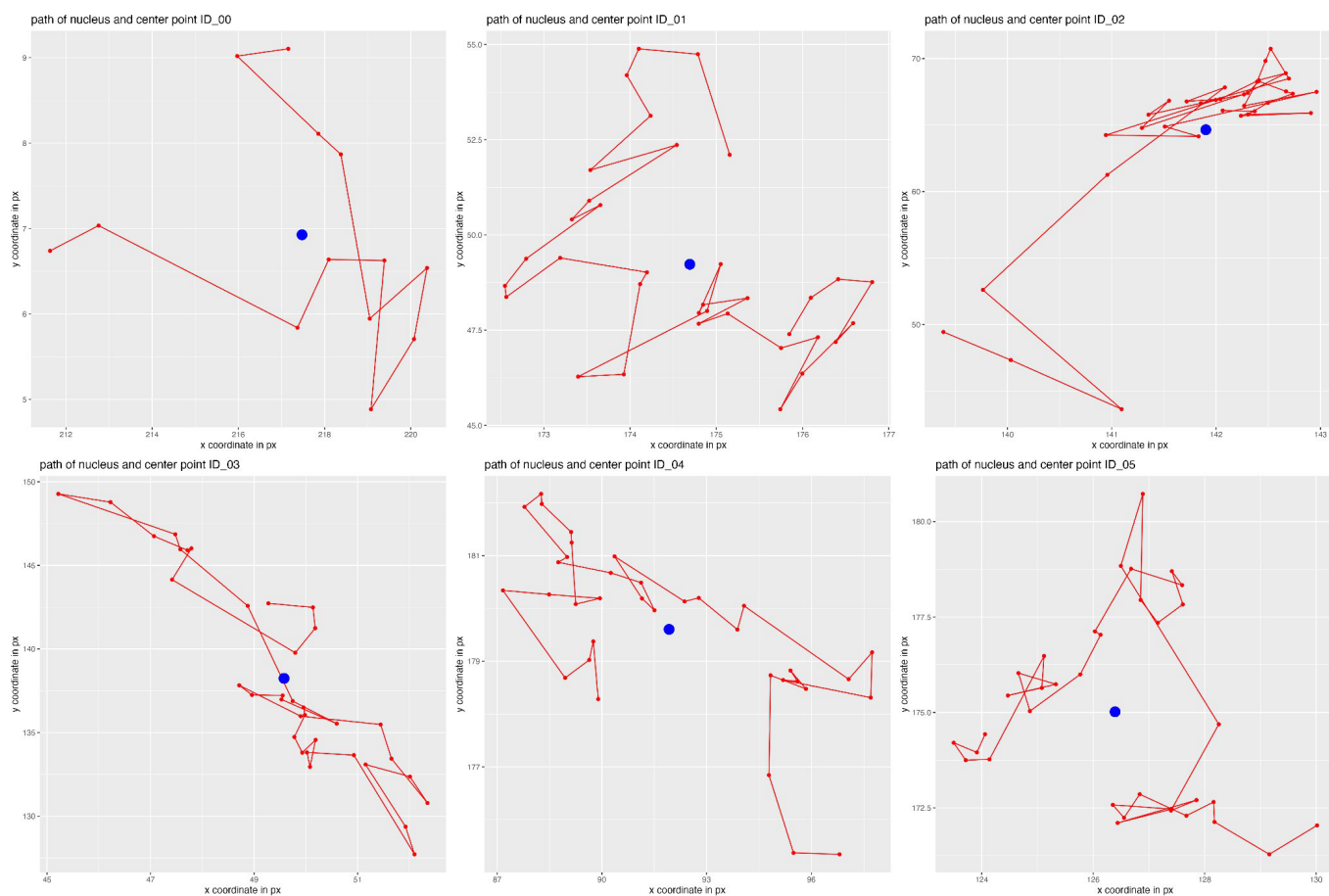

Figure 1.9: Nuclei tracks (red) with calculated center point (blue) for movie\_ID04

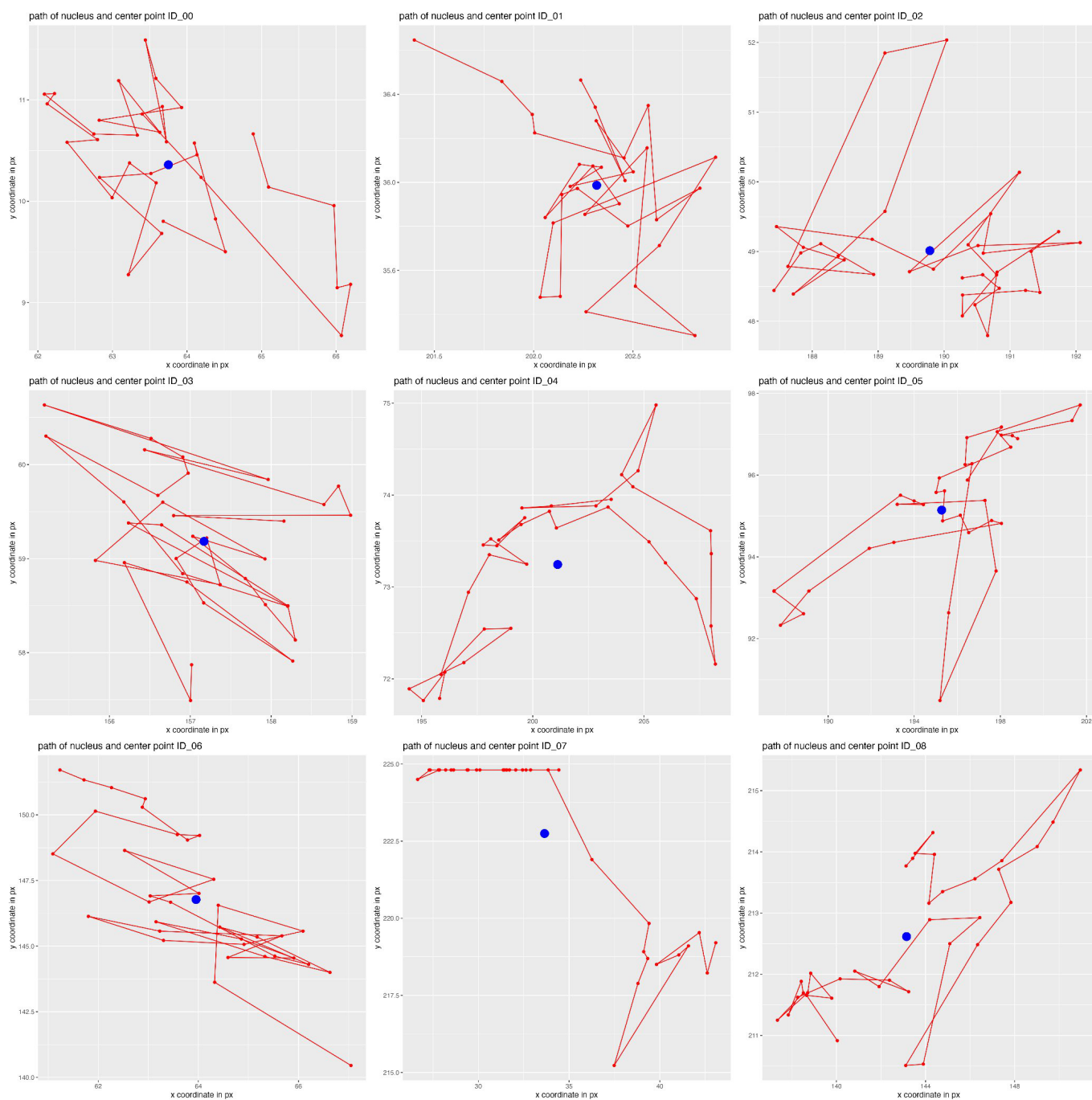

Figure 1.10: Nuclei tracks (red) with calculated center point (blue) for movie\_ID05

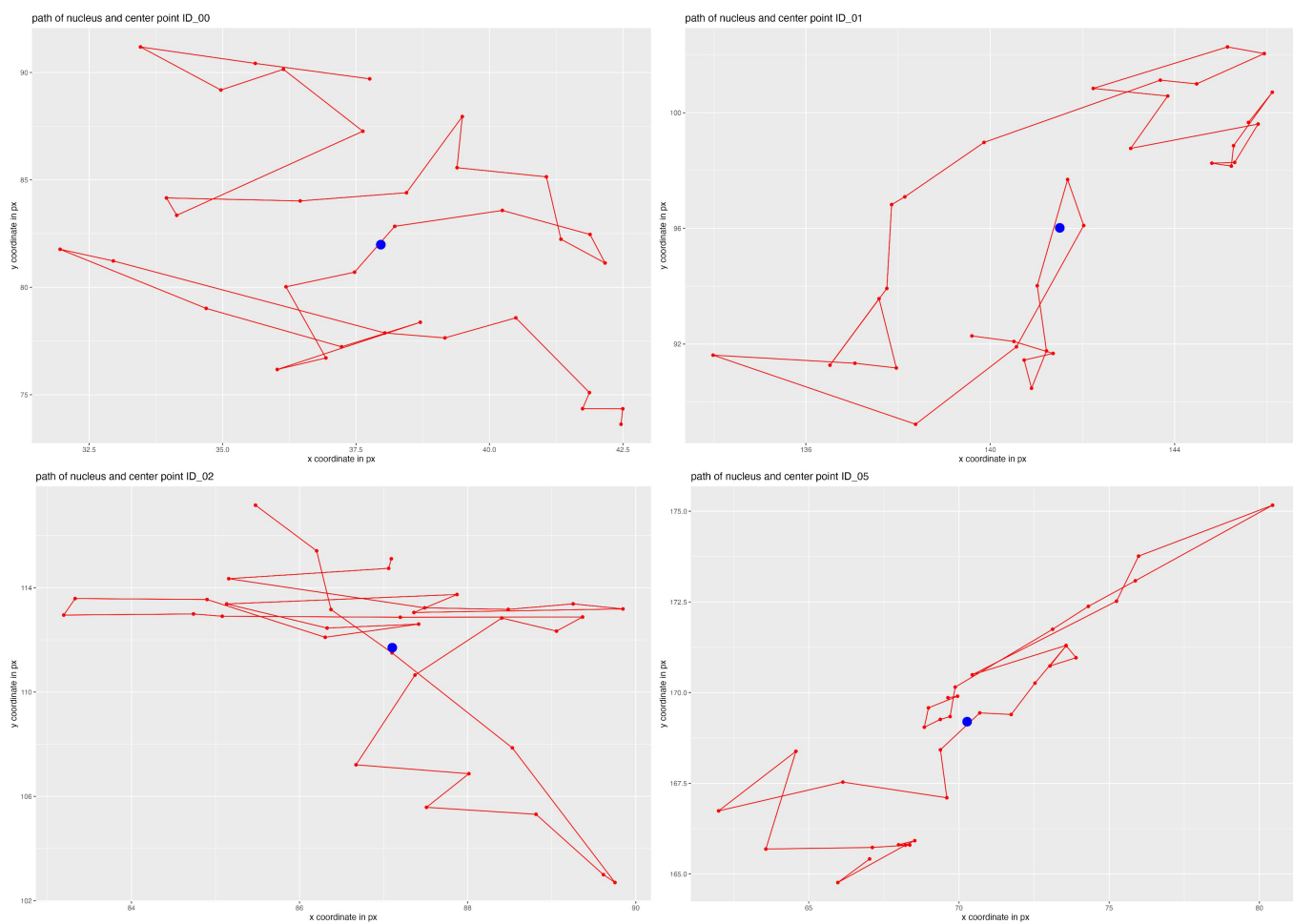

Figure 1.11: Nuclei tracks (red) with calculated center point (blue) for movie\_ID06

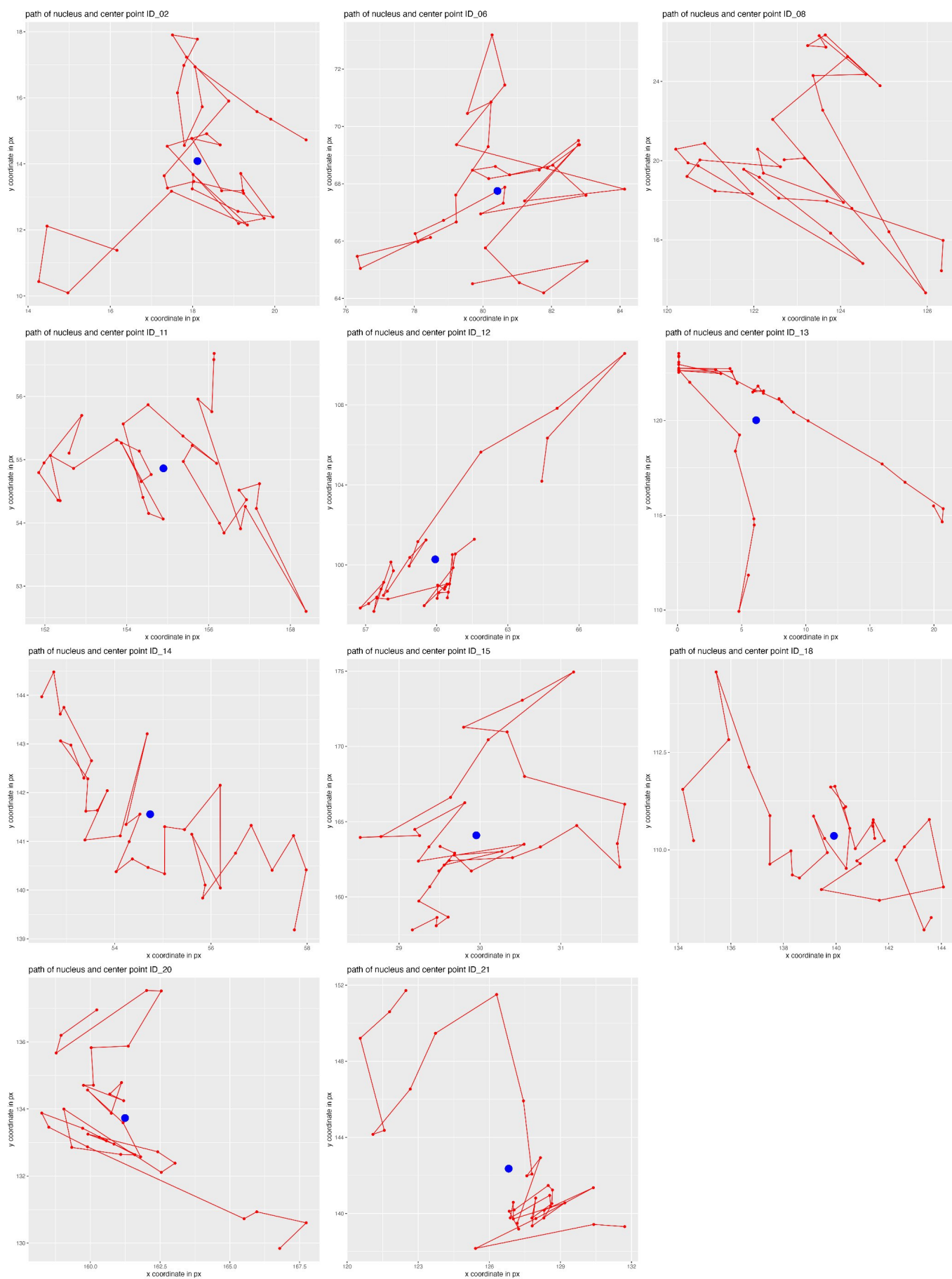

Figure 1.12: Nuclei tracks (red) with calculated center point (blue) for movie\_ID07

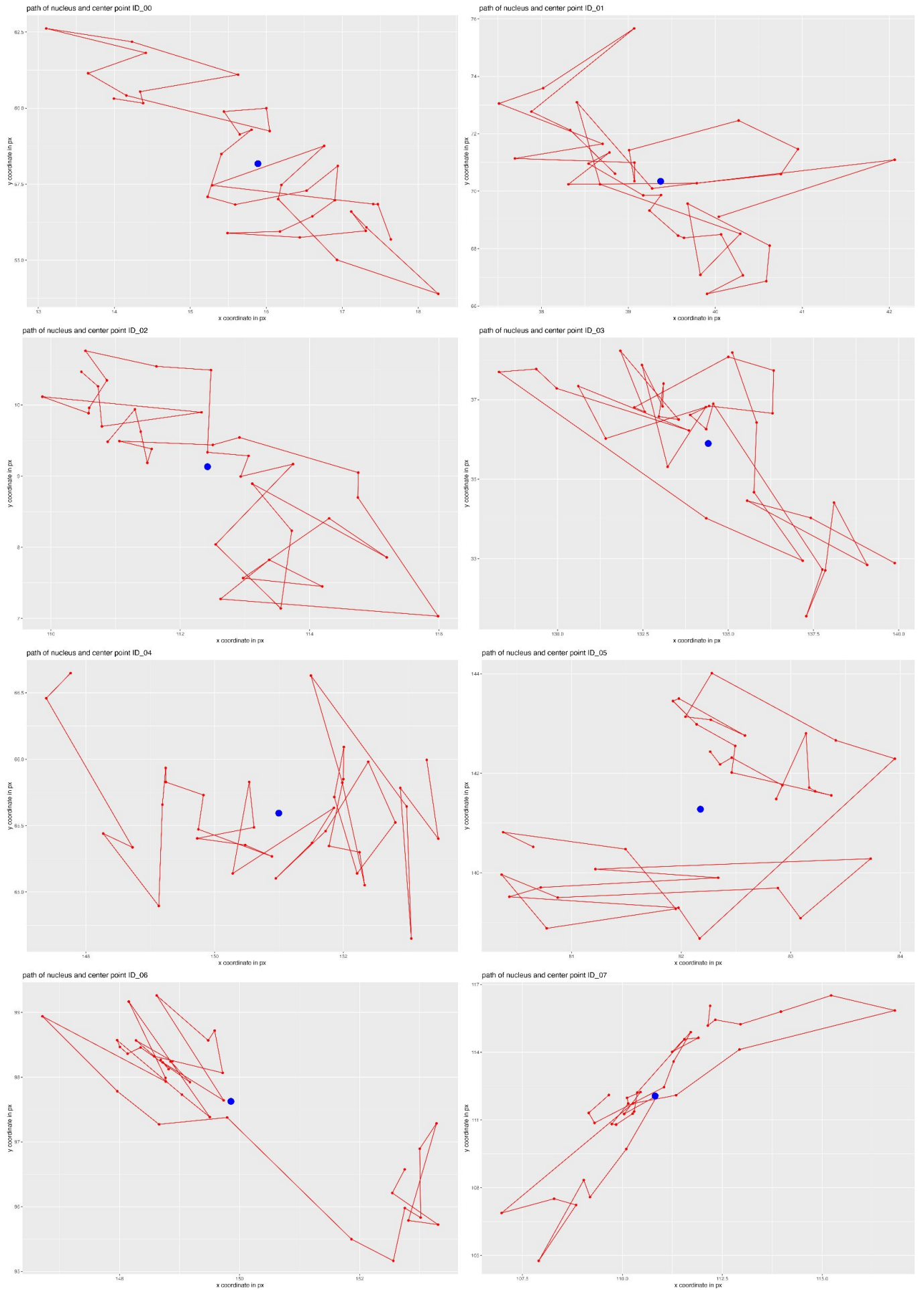

Figure 1.13: Nuclei tracks (red) with calculated center point (blue) for movie\_ID08

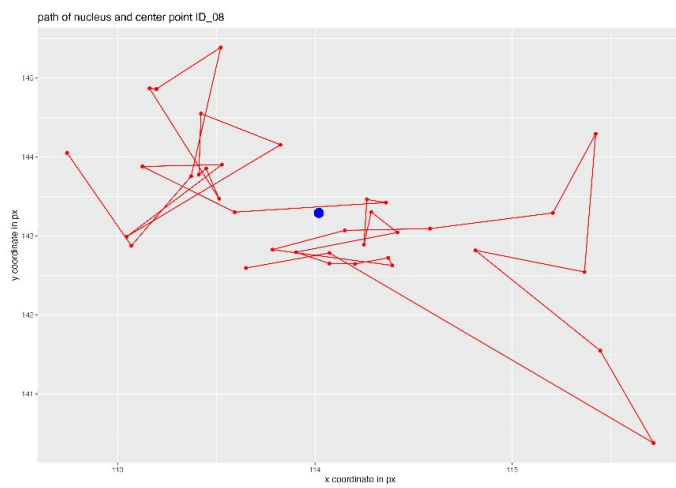

Figure 1.14: Nuclei tracks (red) with calculated center point (blue) for movie\_ID08

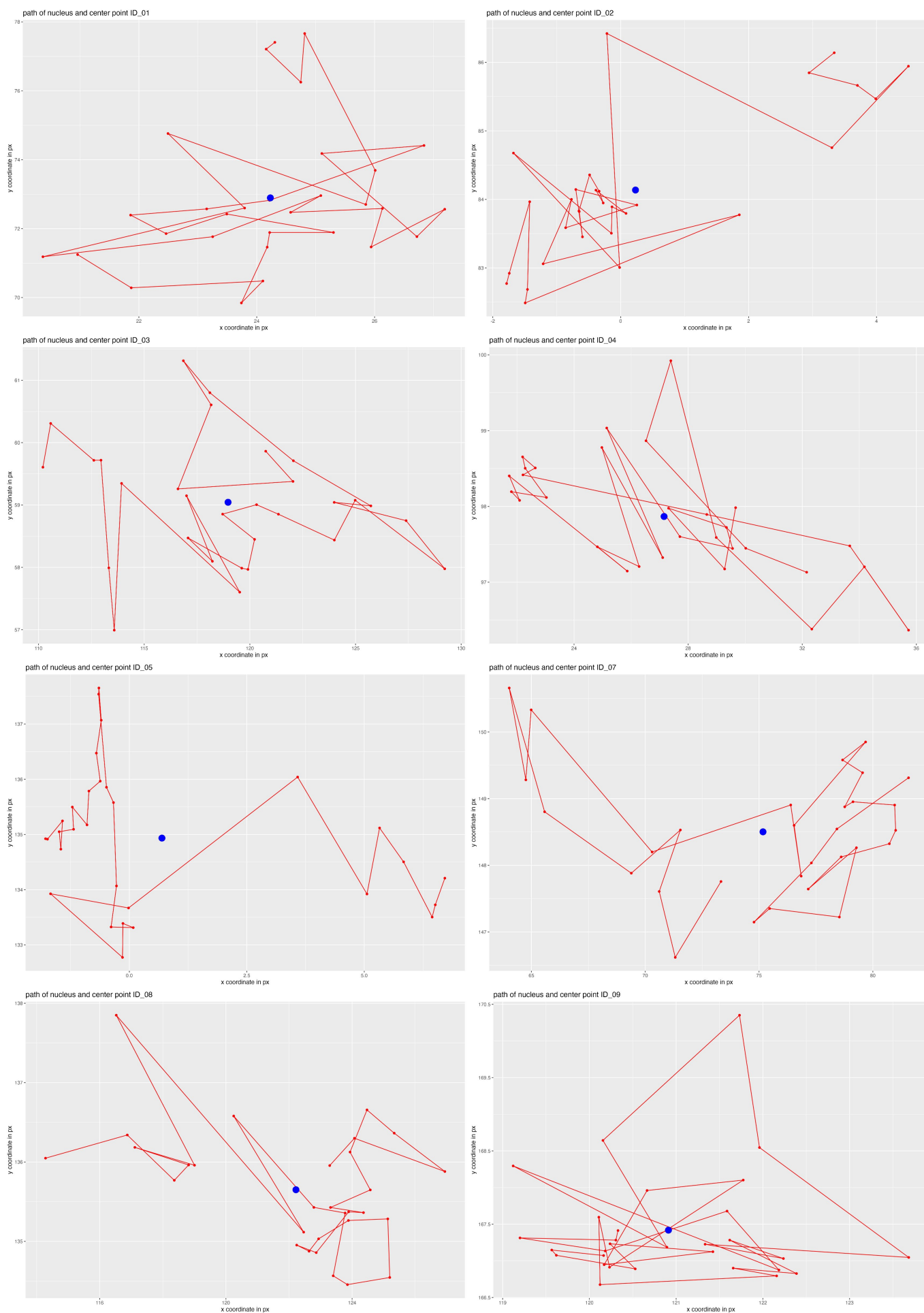

Figure 1.15: Nuclei tracks (red) with calculated center point (blue) for movie\_ID09

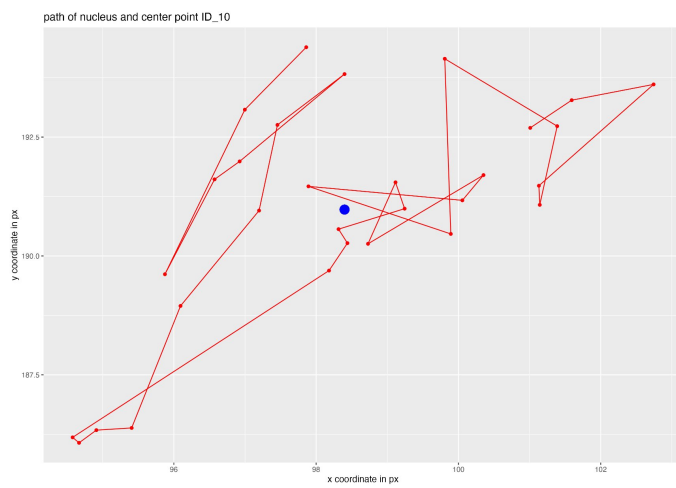

Figure 1.16: Nuclei tracks (red) with calculated center point (blue) for movie\_ID09

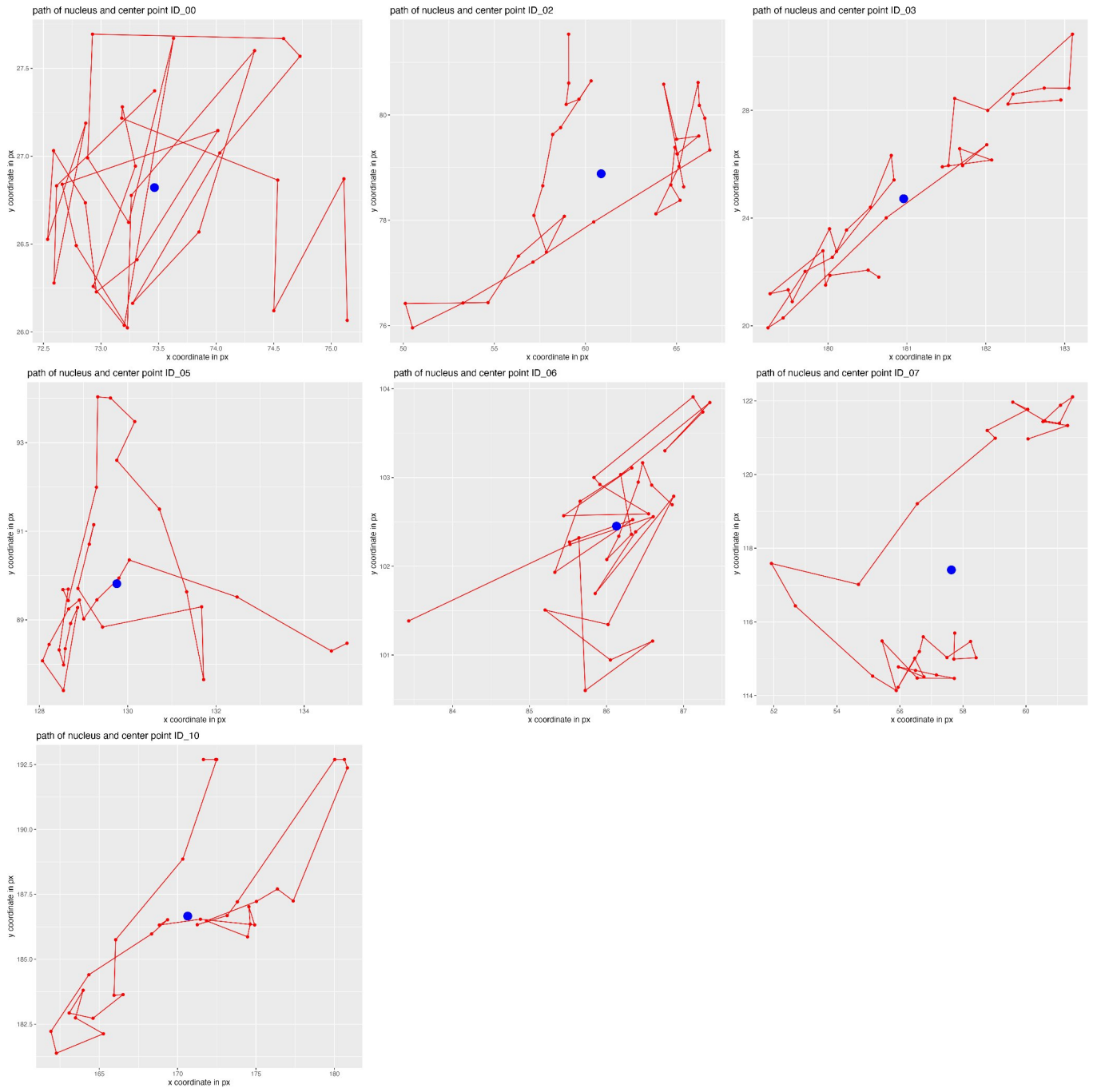

Figure 1.17: Nuclei tracks (red) with calculated center point (blue) for movie\_ID10

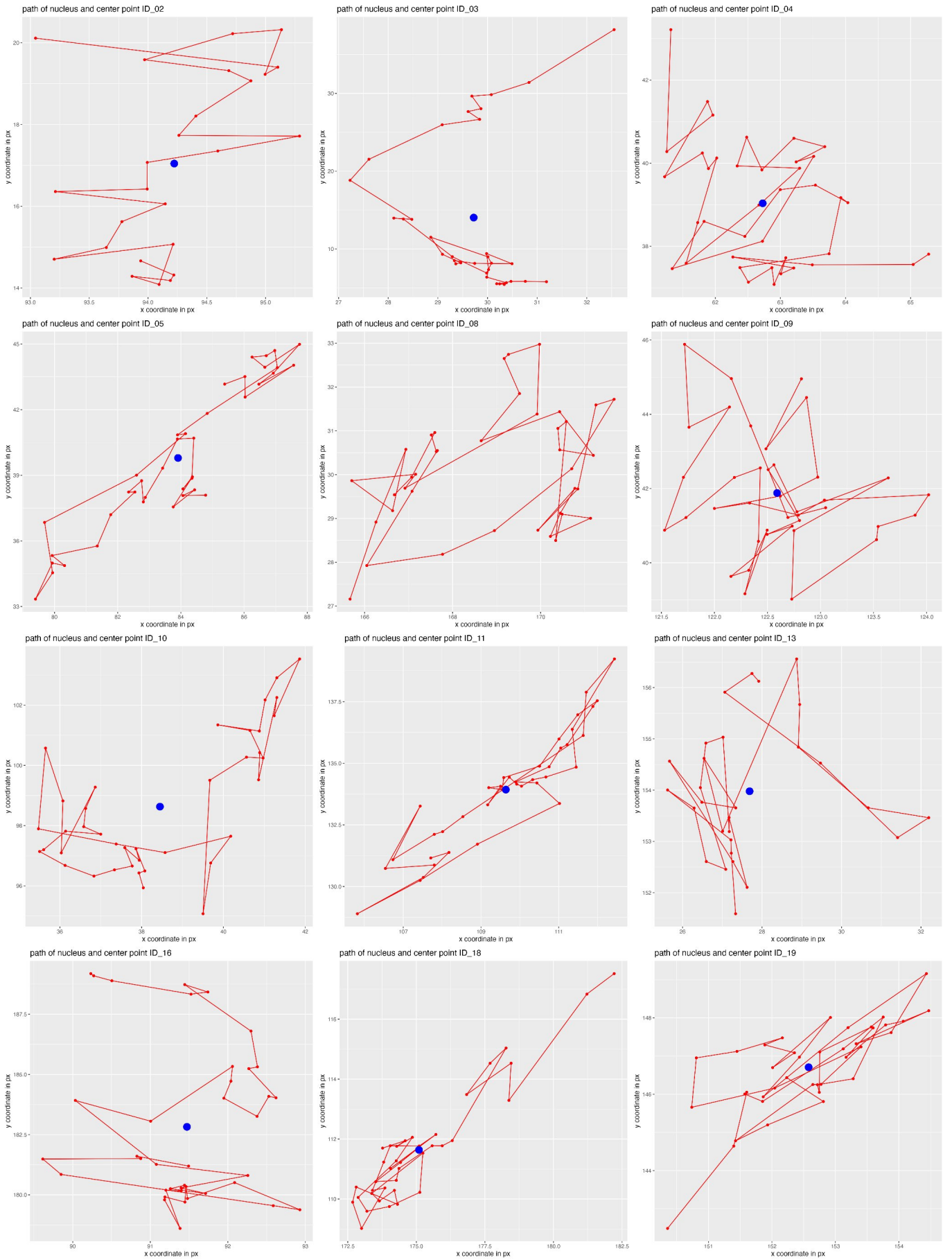

Figure 1.18: Nuclei tracks (red) with calculated center point (blue) for movie\_ID11

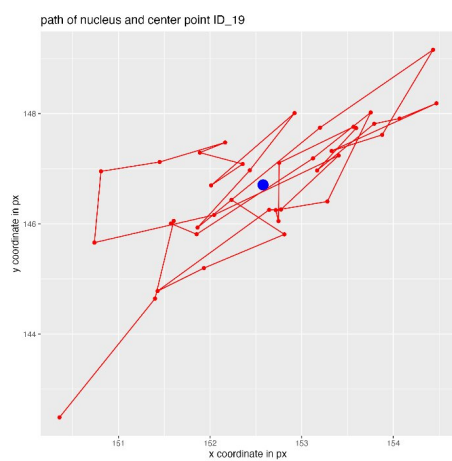

Figure 1.19: Nuclei tracks (red) with calculated center point (blue) for movie\_ID11

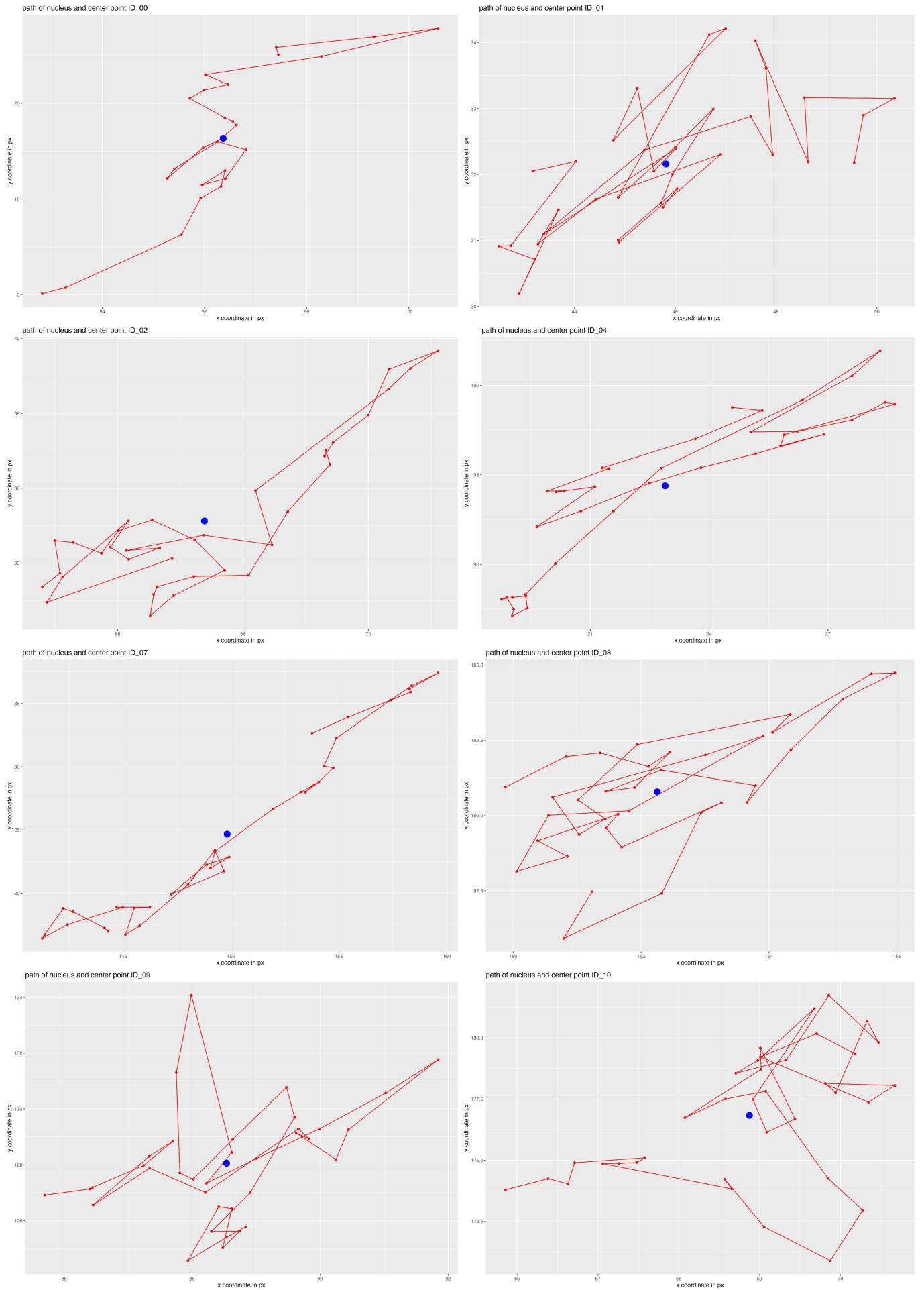

Figure 1.20: Nuclei tracks (red) with calculated center point (blue) for movie\_ID12

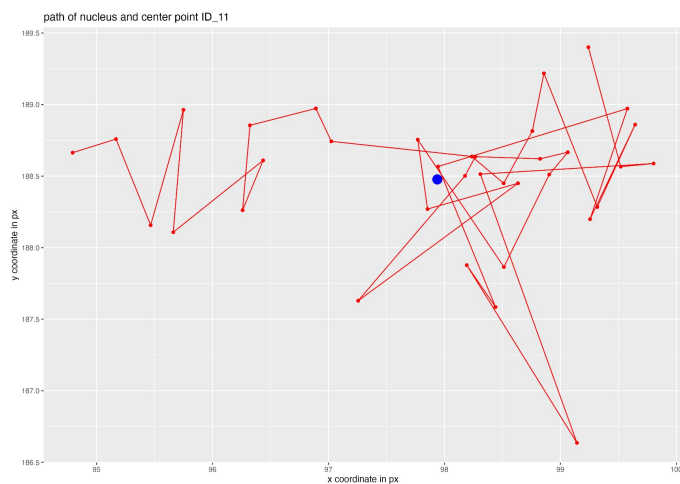

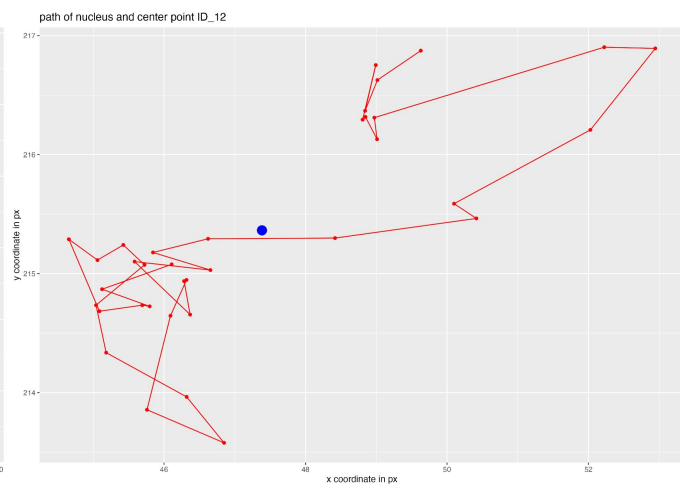

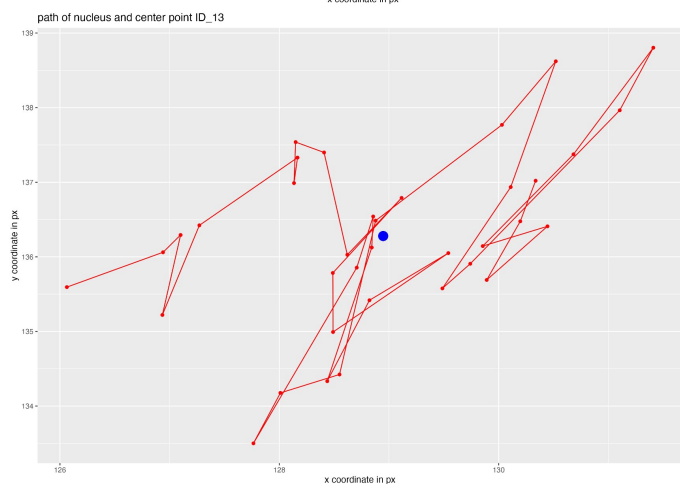

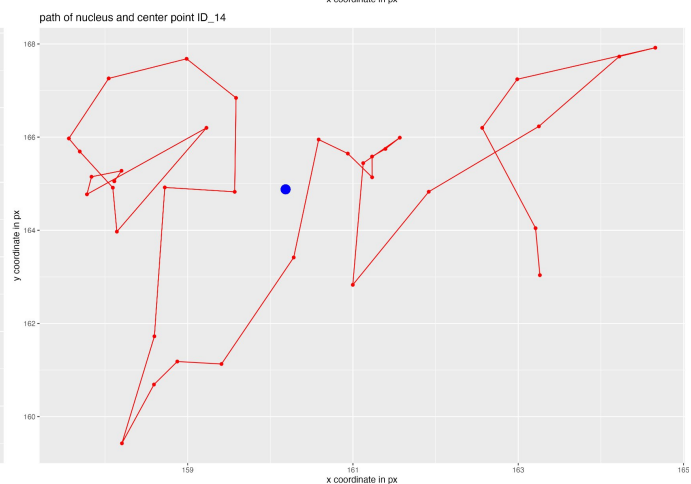

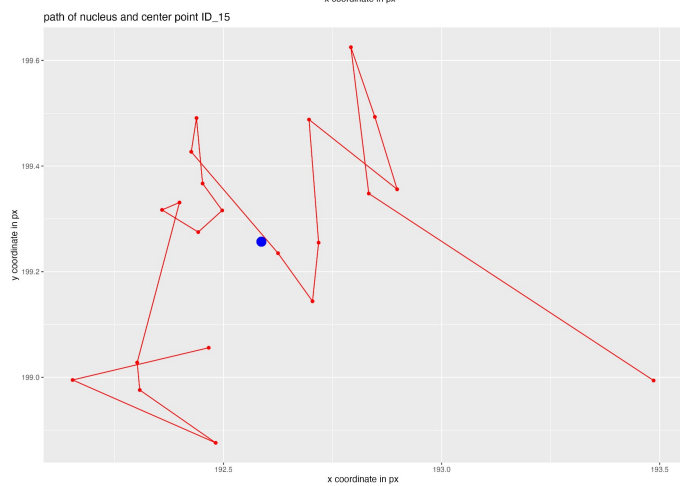

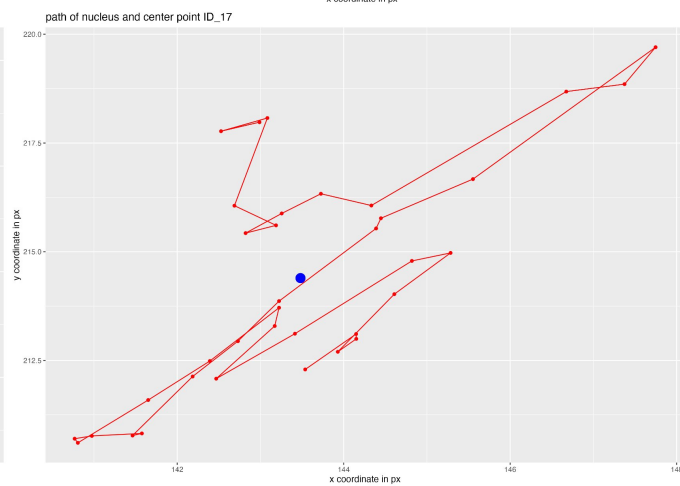

Figure 1.21: Nuclei tracks (red) with calculated center point (blue) for movie\_ID12

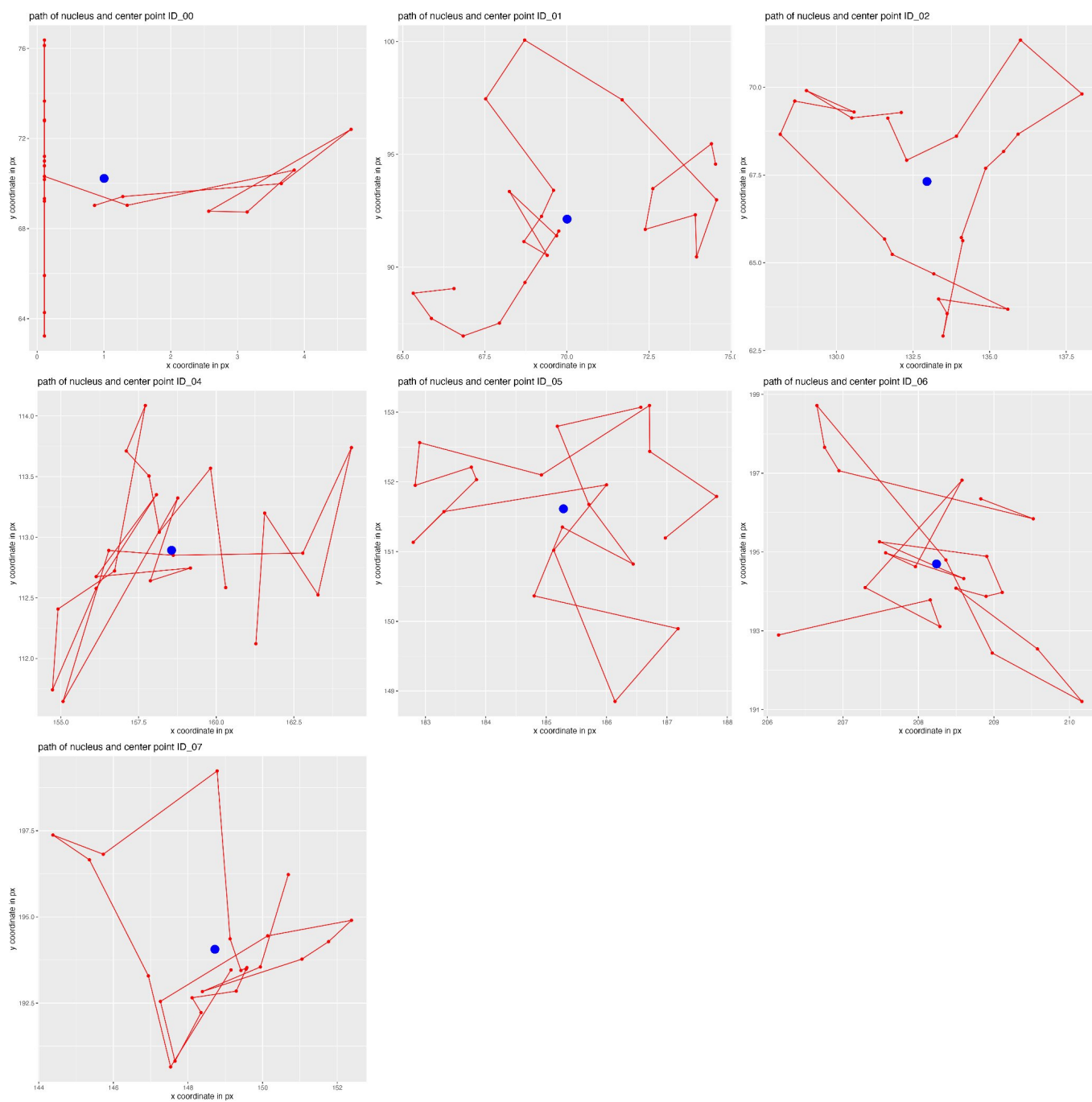

Figure 1.22: Nuclei tracks (red) with calculated center point (blue) for movie\_ID13: nucleus with the ID00 was excluded from the analysis. The track indicates that the nucleus moved to the edge of the field of observation.

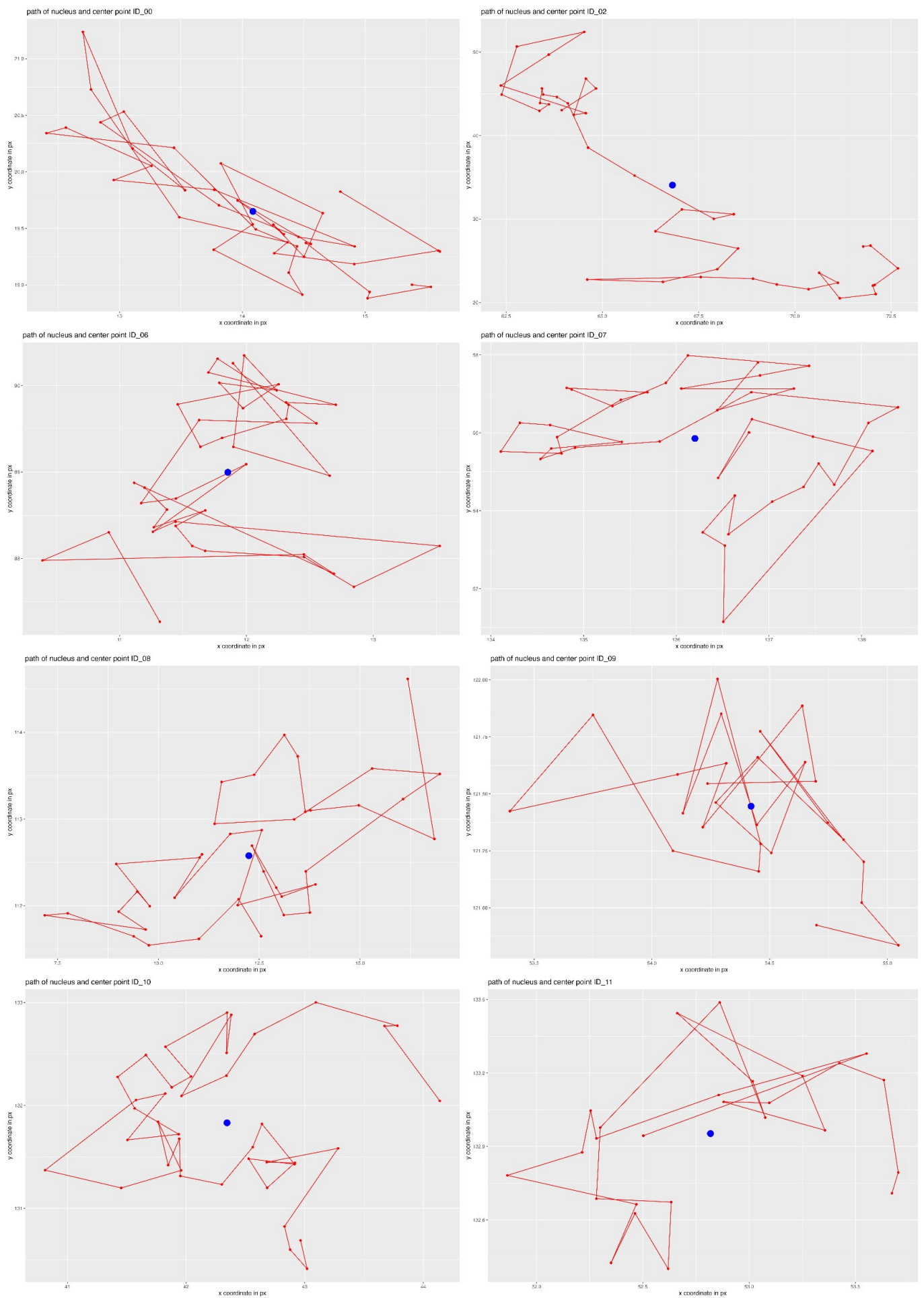

Figure 1.23: Nuclei tracks (red) with calculated center point (blue) for movie\_ID14

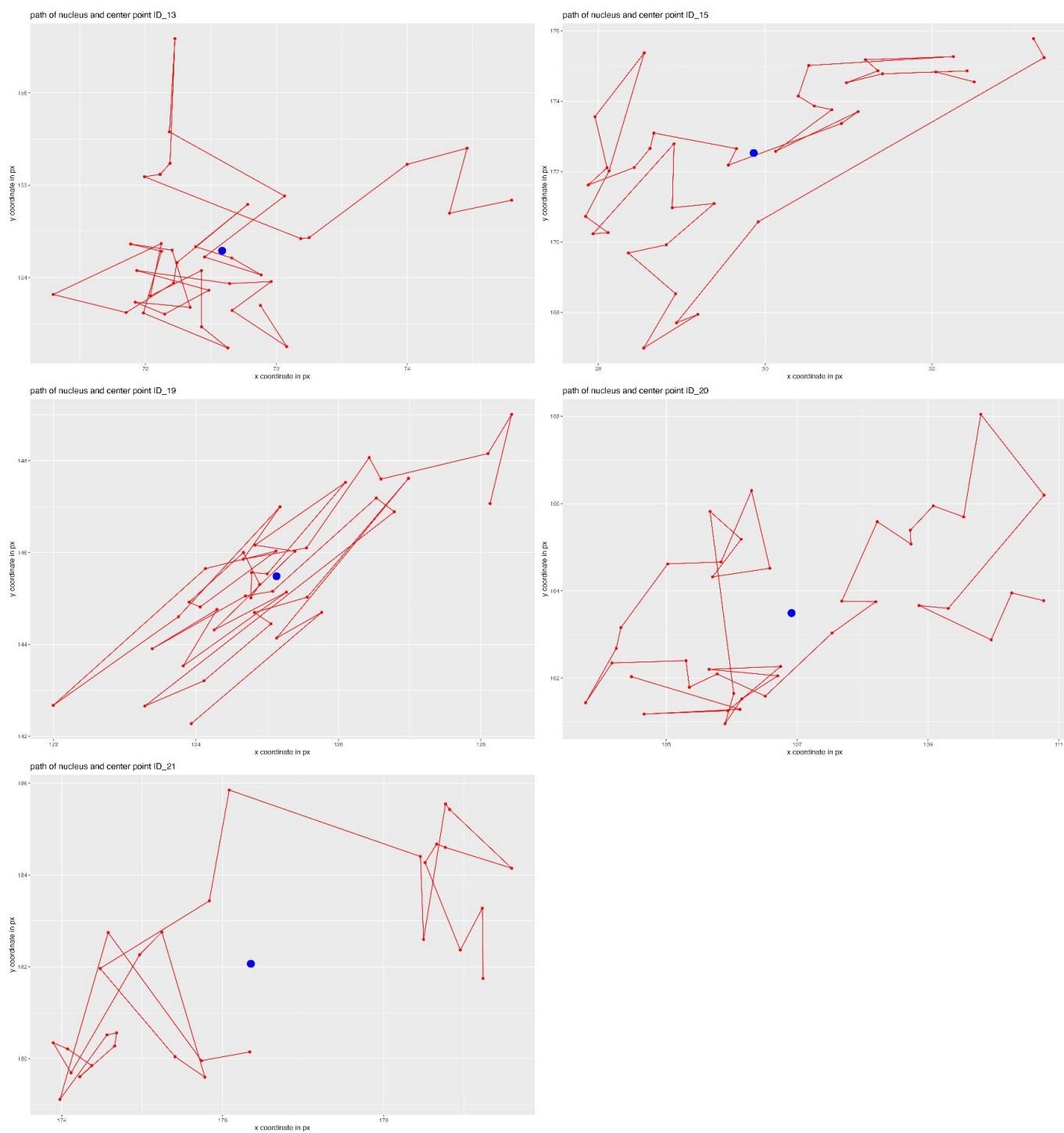

Figure 1.24: Nuclei tracks (red) with calculated center point (blue) for movie\_ID14

##### 1.2.19 Supplemental Information 19 - Distance Distribution to Center Point

**Supported Figure: 7C-F** The following figures depict box plots summarizing all distance measurements from the calculated center point for all considered nuclei. The distance from the center point is measured in $\mu\text{m}$.

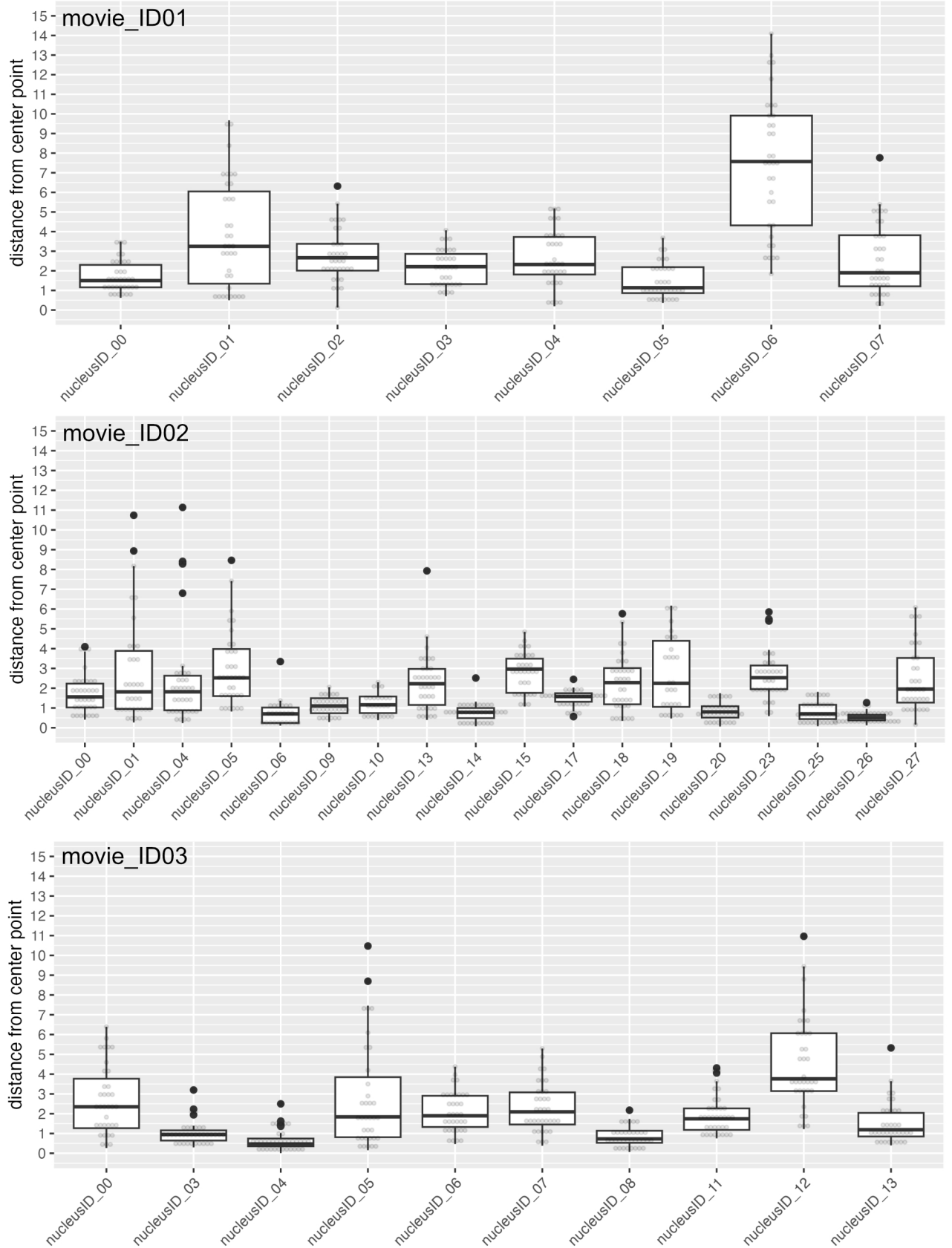

Figure 1.25: Boxplots showing the different distances of individual nuclei from the calculated center point. Individual measurements are shown as overlaid dots.

Figure 1.26: Boxplots showing the different distances of individual nuclei from the calculated center point. Individual measurements are shown as overlaid dots.

Figure 1.27: Boxplots showing the different distances of individual nuclei from the calculated center point. Individual measurements are shown as overlaid dots.

Figure 1.28: Boxplots showing the different distances of individual nuclei from the calculated center point. Individual measurements are shown as overlaid dots.

Figure 1.29: Boxplots showing the different distances of individual nuclei from the calculated center point. Individual measurements are shown as overlaid dots. Based on the track of nucleus 00 the nucleus was excluded from further calculations.

##### 1.2.20 Supplemental Information 20 - Distance Distribution and Nucleus Tracks

**Supported Figure: 7E-F** In the following tables you can find information on the nuclei, which were included in the calculations. The columns contain the following information: movie = ID of time series experiment of the respective nucleus, nucleus = nucleus identifier in the respective time lapse experiment, nuc size $\mu\text{m}^2$ = measured nucleus size in the first frame of the 2D projected 3D time series, cell size $\mu\text{m}^2$ = calculated cell size based on the measured nucleus size (for calculation the earlier described function was used), dist med. $\mu\text{m}$ = median of all the measured distances between the calculated center point and the moving nucleus throughout the time series in $\mu\text{m}$, dist av. $\mu\text{m}$ = average of all the measured distances between the calculated center point and the moving nucleus throughout the time series in $\mu\text{m}$, dist Q1 $\mu\text{m}$ = 25% quartile of all measured distances for this nucleus in $\mu\text{m}$, dist Q3 $\mu\text{m}$ = 75% quartile of all measured distances for this nucleus, dist min $\mu\text{m}$ = smallest measured distance from the calculated center point for this nucleus in $\mu\text{m}$, dist max $\mu\text{m}$ = 1st measured distance from the calculated center point for this nucleus in $\mu\text{m}$, cell class = cell size class as calculated based on the nucleus size (sm = small, meaning below 1000 $\mu\text{m}^2$; med = medium, between 1000 and 3000 $\mu\text{m}^2$; lg = large, meaning above 3000 $\mu\text{m}^2$)

| mov. | nucleus | nuc size<br>$\mu\text{m}^2$ | cell size<br>$\mu\text{m}^2$ | dist<br>med.<br>$\mu\text{m}$ | dist av.<br>$\mu\text{m}$ | dist Q1<br>$\mu\text{m}$ | dist Q3<br>$\mu\text{m}$ | dist min<br>$\mu\text{m}$ | dist max<br>$\mu\text{m}$ | cell class |
| --- | --- | --- | --- | --- | --- | --- | --- | --- | --- | --- |
| 1 | nucleusID 00 | 43.951 | 3276 | 1.5 | 1.73 | 1.16 | 2.3 | 0.62 | 3.49 | lg |
| 1 | nucleusID 01 | 68.218 | 7166 | 3.25 | 3.83 | 1.34 | 6.05 | 0.5 | 9.68 | lg |
| 1 | nucleusID 02 | 25.956 | 1282 | 2.67 | 2.79 | 2.01 | 3.38 | 0.13 | 6.32 | md |
| 1 | nucleusID 03 | 57.315 | 5256 | 2.21 | 2.17 | 1.32 | 2.86 | 0.71 | 4.07 | lg |
| 1 | nucleusID 04 | 88.481 | 11388 | 2.32 | 2.71 | 1.81 | 3.73 | 0.19 | 5.19 | lg |
| 1 | nucleusID 05 | 26.583 | 1338 | 1.14 | 1.47 | 0.86 | 2.19 | 0.37 | 3.68 | md |
| 1 | nucleusID 06 | 35.942 | 2290 | 7.57 | 7.43 | 4.32 | 9.91 | 1.84 | 14.09 | md |
| 1 | nucleusID 07 | 56.639 | 5146 | 1.9 | 2.61 | 1.21 | 3.81 | 0.21 | 7.76 | lg |
| 2 | nucleusID 00 | 77.095 | 8911 | 1.56 | 1.73 | 1.03 | 2.23 | 0.41 | 4.09 | lg |
| 2 | nucleusID 01 | 42.311 | 3061 | 1.82 | 2.84 | 0.95 | 3.89 | 0.27 | 10.74 | lg |
| 2 | nucleusID 04 | 105.029 | 15453 | 1.82 | 2.37 | 0.88 | 2.63 | 0.24 | 11.14 | lg |
| 2 | nucleusID 05 | 39.85 | 2752 | 2.52 | 2.99 | 1.6 | 3.98 | 0.82 | 8.46 | md |
| 2 | nucleusID 06 | 25.184 | 1215 | 0.7 | 0.75 | 0.24 | 1.02 | 0.11 | 3.34 | md |
| 2 | nucleusID 09 | 31.456 | 1806 | 1.1 | 1.09 | 0.74 | 1.49 | 0.29 | 2.05 | md |
| 2 | nucleusID 10 | 29.429 | 1604 | 1.16 | 1.21 | 0.74 | 1.58 | 0.38 | 2.3 | md |
| 2 | nucleusID 13 | 92.099 | 12230 | 2.23 | 2.3 | 1.16 | 2.97 | 0.4 | 7.93 | lg |
| 2 | nucleusID 14 | 11.386 | 296 | 0.78 | 0.79 | 0.49 | 1 | 0.07 | 2.52 | sm |
| 2 | nucleusID 15 | 32.614 | 1926 | 2.96 | 2.82 | 1.77 | 3.49 | 1.08 | 4.85 | md |
| 2 | nucleusID 17 | 23.785 | 1098 | 1.58 | 1.5 | 1.32 | 1.75 | 0.56 | 2.45 | md |
| 2 | nucleusID 18 | 38.837 | 2628 | 2.28 | 2.24 | 1.19 | 3.02 | 0.35 | 5.76 | md |
| 2 | nucleusID 19 | 50.609 | 4211 | 2.25 | 2.78 | 1.06 | 4.39 | 0.47 | 6.17 | lg |
| 2 | nucleusID 20 | 17.272 | 621 | 0.81 | 0.83 | 0.51 | 1.09 | 0.06 | 1.75 | sm |
| 2 | nucleusID 23 | 17.223 | 618 | 2.53 | 2.67 | 1.95 | 3.15 | 0.59 | 5.85 | sm |
| 2 | nucleusID 25 | 13.46 | 398 | 0.7 | 0.85 | 0.44 | 1.16 | 0.07 | 1.82 | sm |
| 2 | nucleusID 26 | 10.807 | 269 | 0.52 | 0.54 | 0.37 | 0.65 | 0.12 | 1.27 | sm |
| 2 | nucleusID 27 | 30.684 | 1728 | 1.95 | 2.5 | 1.27 | 3.53 | 0.16 | 6.08 | md |
| 3 | nucleusID 04 | 28.947 | 1557 | 0.47 | 0.64 | 0.35 | 0.74 | 0.01 | 2.5 | md |
| 3 | nucleusID 05 | 59.245 | 5575 | 1.84 | 2.84 | 0.81 | 3.85 | 0.15 | 10.47 | lg |
| 3 | nucleusID 06 | 73.236 | 8132 | 1.89 | 2.11 | 1.33 | 2.9 | 0.47 | 4.38 | lg |
| 3 | nucleusID 07 | 63.345 | 6281 | 2.1 | 2.29 | 1.45 | 3.08 | 0.38 | 5.29 | lg |
| 3 | nucleusID 08 | 28.271 | 1493 | 0.73 | 0.87 | 0.53 | 1.14 | 0.06 | 2.18 | md |
| 3 | nucleusID 11 | 81.051 | 9741 | 1.74 | 1.91 | 1.18 | 2.27 | 0.75 | 4.31 | lg |
| 3 | nucleusID 12 | 59.872 | 5681 | 3.76 | 4.59 | 3.14 | 6.07 | 1.22 | 10.96 | lg |
| 3 | nucleusID 13 | 47.569 | 3771 | 1.19 | 1.55 | 0.85 | 2.04 | 0.4 | 5.33 | lg |
| 4 | nucleusID 01 | 38.403 | 2576 | 2.17 | 2.37 | 1.54 | 2.95 | 0.36 | 5.69 | md |
| 4 | nucleusID 02 | 73.863 | 8256 | 2.68 | 4.1 | 1.54 | 3.72 | 0.46 | 21.02 | lg |
| 4 | nucleusID 03 | 128.042 | 21990 | 4.46 | 5.09 | 2.59 | 7.92 | 0.96 | 11.87 | lg |
| 4 | nucleusID 04 | 38.789 | 2622 | 3.24 | 3.24 | 2.13 | 4.22 | 0.55 | 6.48 | md |
| 4 | nucleusID 05 | 27.741 | 1444 | 2.74 | 2.79 | 2.09 | 3.04 | 1.16 | 5.73 | md |
| 5 | nucleusID 00 | 57.074 | 5216 | 0.93 | 1.06 | 0.54 | 1.32 | 0.23 | 2.87 | lg |
| 5 | nucleusID 01 | 7.381 | 137 | 0.32 | 0.37 | 0.17 | 0.5 | 0.09 | 1.13 | sm |
| 5 | nucleusID 02 | 16.403 | 566 | 1.31 | 1.43 | 0.89 | 1.94 | 0.27 | 3.03 | sm |
| 5 | nucleusID 03 | 137.208 | 24871 | 1.01 | 1.05 | 0.65 | 1.34 | 0.05 | 2.45 | lg |
| 5 | nucleusID 04 | 28.802 | 1543 | 3.37 | 3.74 | 2.28 | 5.29 | 0.4 | 7.17 | md |
| 5 | nucleusID 05 | 54.082 | 4740 | 2.53 | 3.21 | 1.48 | 3.84 | 0.27 | 8.03 | lg |
| 5 | nucleusID 06 | 137.691 | 25027 | 2.35 | 2.56 | 1.55 | 3.34 | 0.24 | 7.04 | lg |
| 5 | nucleusID 07 | 65.131 | 6599 | 5.55 | 5.38 | 2.94 | 7.03 | 2.07 | 10.08 | lg |
| 5 | nucleusID 08 | 95.091 | 12946 | 3.21 | 3.36 | 1.81 | 4.65 | 0.91 | 8.26 | lg |

Table 1.27: Information on nuclei included in the calulations movie 01-05

| movie | nucleus | nuc size<br>$\mu\text{m}^2$ | cell size<br>$\mu\text{m}^2$ | dist<br>med.<br>$\mu\text{m}$ | dist av.<br>$\mu\text{m}$ | dist Q1<br>$\mu\text{m}$ | dist Q3<br>$\mu\text{m}$ | dist min<br>$\mu\text{m}$ | dist max<br>$\mu\text{m}$ | cell class |
| --- | --- | --- | --- | --- | --- | --- | --- | --- | --- | --- |
| 6 | nucleusID 00 | 136.712 | 24711 | 4.54 | 5.26 | 3.88 | 7.33 | 0.89 | 10.25 | lg |
| 6 | nucleusID 01 | 134.408 | 23974 | 4.63 | 4.85 | 4.09 | 5.75 | 0.53 | 8.71 | lg |
| 6 | nucleusID 02 | 54.763 | 4846 | 2.71 | 3.21 | 1.66 | 4.1 | 0.2 | 9.39 | lg |
| 6 | nucleusID 05 | 100.912 | 14391 | 3.9 | 3.82 | 1.37 | 5.1 | 0.48 | 11.79 | lg |
| 7 | nucleusID 02 | 90.385 | 11828 | 1.97 | 2.06 | 1 | 2.79 | 0.42 | 5.31 | lg |
| 7 | nucleusID 06 | 66.672 | 6880 | 2.54 | 2.36 | 1.32 | 3.19 | 0.25 | 5.45 | lg |
| 7 | nucleusID 11 | 47.284 | 3731 | 1.8 | 1.71 | 0.94 | 2.26 | 0.32 | 4.16 | lg |
| 7 | nucleusID 12 | 63.305 | 6273 | 1.95 | 2.91 | 1.35 | 3.12 | 0.76 | 13.04 | lg |
| 7 | nucleusID 13 | 41.329 | 2936 | 5.53 | 5.8 | 2.37 | 6.91 | 1.51 | 15.47 | md |
| 7 | nucleusID 14 | 23.961 | 1112 | 1.57 | 1.77 | 1.05 | 2.29 | 0.21 | 3.82 | md |
| 7 | nucleusID 15 | 34.205 | 2096 | 2.17 | 3.21 | 1.28 | 4.93 | 0.7 | 10.91 | md |
| 7 | nucleusID 18 | 29.951 | 1655 | 1.68 | 2.29 | 1.15 | 3.17 | 0.36 | 6.14 | md |
| 7 | nucleusID 20 | 44.342 | 3328 | 1.78 | 2.34 | 1.12 | 3.06 | 0.16 | 7.21 | lg |
| 7 | nucleusID 21 | 63.66 | 6336 | 2.89 | 3.93 | 2.23 | 5.13 | 0.85 | 10.32 | lg |
| 8 | nucleusID 00 | 38.033 | 2532 | 2.04 | 2.23 | 1.18 | 2.84 | 0.58 | 5.25 | md |
| 8 | nucleusID 01 | 91.236 | 12027 | 1.88 | 1.89 | 0.98 | 2.81 | 0.27 | 5.33 | lg |
| 8 | nucleusID 02 | 66.602 | 6867 | 1.62 | 1.65 | 1.06 | 2.1 | 0.2 | 4.14 | lg |
| 8 | nucleusID 03 | 81.843 | 9911 | 2.3 | 2.8 | 1.58 | 4.05 | 0.37 | 6.39 | lg |
| 8 | nucleusID 04 | 31.511 | 1811 | 1.29 | 1.47 | 0.86 | 1.92 | 0.34 | 3.72 | md |
| 8 | nucleusID 05 | 25.414 | 1235 | 1.8 | 1.72 | 1.21 | 2.16 | 0.72 | 2.78 | md |
| 8 | nucleusID 06 | 48.843 | 3953 | 1.78 | 1.92 | 1.14 | 3.05 | 0.12 | 3.94 | lg |
| 8 | nucleusID 07 | 61.533 | 5964 | 2.23 | 2.76 | 0.94 | 4.15 | 0.06 | 7.86 | lg |
| 8 | nucleusID 08 | 42.463 | 3081 | 0.79 | 0.99 | 0.52 | 1.24 | 0.26 | 3.37 | lg |
| 9 | nucleusID 01 | 32.184 | 1881 | 2.14 | 2.28 | 1.44 | 3.07 | 0.07 | 4.8 | md |
| 9 | nucleusID 02 | 21.515 | 918 | 1.44 | 1.76 | 0.8 | 2.37 | 0.22 | 4.63 | sm |
| 9 | nucleusID 03 | 71.67 | 7825 | 3.1 | 4 | 1.81 | 5.95 | 0.32 | 10.31 | lg |
| 9 | nucleusID 04 | 36.544 | 2358 | 2.47 | 3.4 | 1.9 | 5.05 | 0.19 | 8.7 | md |
| 9 | nucleusID 05 | 37.714 | 2494 | 2.18 | 2.73 | 1.76 | 3.01 | 1.22 | 6.06 | md |
| 9 | nucleusID 07 | 57.847 | 5343 | 4.04 | 4.56 | 2.44 | 5.74 | 1.19 | 11.36 | lg |
| 9 | nucleusID 08 | 81.737 | 9888 | 2.09 | 2.59 | 1.24 | 3.22 | 0.59 | 7.95 | lg |
| 9 | nucleusID 09 | 20.31 | 829 | 0.89 | 1.14 | 0.7 | 1.39 | 0.23 | 3.04 | sm |
| 10 | nucleusID 00 | 19.566 | 775 | 0.78 | 0.83 | 0.6 | 1.03 | 0.21 | 1.84 | sm |
| 10 | nucleusID 02 | 46.717 | 3652 | 4.1 | 4.38 | 3.1 | 5.4 | 1 | 11.03 | lg |
| 10 | nucleusID 03 | 75.605 | 8606 | 2.93 | 2.86 | 1.61 | 3.91 | 0.53 | 6.47 | lg |
| 10 | nucleusID 05 | 82.056 | 9957 | 1.88 | 2.03 | 1.09 | 2.7 | 0.13 | 5.39 | lg |
| 10 | nucleusID 06 | 20.665 | 855 | 0.69 | 0.89 | 0.52 | 1.11 | 0.12 | 2.9 | sm |
| 10 | nucleusID 07 | 42.747 | 3118 | 3.13 | 3.65 | 2.67 | 4.97 | 1.72 | 6.07 | lg |
| 10 | nucleusID 10 | 110.943 | 17036 | 5.6 | 5.53 | 3.22 | 7.21 | 0.69 | 11.68 | lg |

Table 1.28: Information on nuclei included in the calculations movie 06-10

| movie | nucleus | nuc size<br>$\mu\text{m}^2$ | cell size<br>$\mu\text{m}^2$ | dist<br>med.<br>$\mu\text{m}$ | dist av.<br>$\mu\text{m}$ | dist Q1<br>$\mu\text{m}$ | dist Q3<br>$\mu\text{m}$ | dist min<br>$\mu\text{m}$ | dist max<br>$\mu\text{m}$ | cell class |
| --- | --- | --- | --- | --- | --- | --- | --- | --- | --- | --- |
| 11 | nucleusID 02 | 15.348 | 503 | 2.32 | 2.01 | 1.22 | 2.73 | 0.23 | 3.4 | sm |
| 11 | nucleusID 03 | 39.025 | 2651 | 7.4 | 8.25 | 5.44 | 8.6 | 1.27 | 24.34 | md |
| 11 | nucleusID 04 | 33.708 | 2042 | 1.56 | 1.56 | 1.11 | 1.79 | 0.07 | 4.41 | md |
| 11 | nucleusID 05 | 34.417 | 2120 | 3.36 | 3.46 | 1.55 | 5.18 | 0.67 | 7.87 | md |
| 11 | nucleusID 09 | 65.928 | 6744 | 1.12 | 1.39 | 0.68 | 1.89 | 0.09 | 4.1 | lg |
| 11 | nucleusID 10 | 42.499 | 3086 | 2.68 | 2.78 | 1.99 | 3.24 | 1.5 | 5.98 | lg |
| 11 | nucleusID 11 | 45.618 | 3500 | 2.33 | 2.37 | 0.82 | 3.56 | 0.09 | 6.32 | lg |
| 11 | nucleusID 13 | 15.986 | 541 | 1.64 | 1.82 | 1.25 | 2.11 | 0.48 | 4.54 | sm |
| 11 | nucleusID 16 | 35.268 | 2214 | 2.57 | 2.94 | 1.89 | 3.29 | 0.52 | 6.48 | md |
| 11 | nucleusID 18 | 55.365 | 4942 | 1.62 | 2.13 | 0.88 | 2.66 | 0.07 | 9.23 | lg |
| 11 | nucleusID 19 | 35.8 | 2274 | 1.15 | 1.29 | 0.7 | 1.64 | 0.3 | 4.77 | md |
| 11 | nucleusID 20 | 66.814 | 6906 | 2.13 | 2.35 | 1.26 | 2.95 | 0.15 | 5.35 | lg |
| 12 | nucleusID 00 | 39.416 | 2698 | 4.96 | 5.93 | 2.44 | 8.79 | 0.06 | 16.62 | md |
| 12 | nucleusID 01 | 30.925 | 1752 | 1.97 | 1.96 | 1.09 | 2.82 | 0.2 | 4.64 | md |
| 12 | nucleusID 02 | 33.337 | 2003 | 2.4 | 2.88 | 1.56 | 3.5 | 0.57 | 7.78 | md |
| 12 | nucleusID 04 | 48.534 | 3909 | 4.6 | 4.72 | 2.69 | 7.27 | 0.42 | 9.3 | lg |
| 12 | nucleusID 07 | 63.587 | 6323 | 7.63 | 7.79 | 4.88 | 9.77 | 1.4 | 16.06 | lg |
| 12 | nucleusID 08 | 68.604 | 7239 | 1.75 | 2.14 | 1.26 | 2.6 | 0.38 | 5.42 | lg |
| 12 | nucleusID 09 | 56.205 | 5076 | 1.81 | 2.07 | 1.2 | 2.59 | 0.39 | 6.04 | lg |
| 12 | nucleusID 10 | 32.951 | 1961 | 2.59 | 2.64 | 1.7 | 3.48 | 0.58 | 6.03 | md |
| 12 | nucleusID 11 | 24.991 | 1199 | 1.15 | 1.26 | 0.68 | 1.7 | 0.09 | 3.15 | md |
| 12 | nucleusID 12 | 71.402 | 7773 | 1.83 | 2.14 | 1.65 | 2.4 | 0.77 | 5.77 | lg |
| 12 | nucleusID 13 | 37.438 | 2462 | 1.35 | 1.48 | 0.87 | 2.01 | 0.19 | 3.53 | md |
| 12 | nucleusID 14 | 74.104 | 8304 | 2.3 | 2.59 | 1.6 | 3.4 | 0.62 | 5.8 | lg |
| 12 | nucleusID 17 | 50.995 | 4269 | 1.92 | 2.61 | 1.46 | 3.64 | 0.59 | 6.81 | lg |
| 13 | nucleusID 01 | 156.024 | 31266 | 4.28 | 3.82 | 1.93 | 5.54 | 0.59 | 8.04 | lg |
| 13 | nucleusID 02 | 78.157 | 9130 | 3.05 | 3.19 | 2.14 | 4.47 | 0.9 | 5.64 | lg |
| 13 | nucleusID 04 | 147.243 | 28202 | 1.84 | 2.22 | 0.84 | 3.35 | 0.07 | 5.85 | lg |
| 13 | nucleusID 05 | 37.969 | 2525 | 1.65 | 1.65 | 1.19 | 2.48 | 0.26 | 2.89 | md |
| 13 | nucleusID 06 | 52.056 | 4428 | 1.36 | 1.7 | 0.78 | 2.5 | 0.16 | 4.32 | lg |
| 13 | nucleusID 07 | 67.157 | 6969 | 1.94 | 2.4 | 1.3 | 3.52 | 0.51 | 5.46 | lg |
| 14 | nucleusID 00 | 50.119 | 4139 | 0.67 | 0.86 | 0.46 | 1.25 | 0.11 | 2.11 | lg |
| 14 | nucleusID 02 | 69.579 | 7423 | 11.27 | 10.55 | 8.91 | 12.54 | 1.51 | 18.5 | lg |
| 14 | nucleusID 06 | 40.266 | 2803 | 0.9 | 0.95 | 0.73 | 1.16 | 0.17 | 1.88 | md |
| 14 | nucleusID 07 | 79.716 | 9457 | 1.63 | 1.68 | 1.29 | 1.93 | 0.39 | 4.71 | lg |
| 14 | nucleusID 08 | 65.893 | 6737 | 1.63 | 2.03 | 0.97 | 2.85 | 0.14 | 5.1 | lg |
| 14 | nucleusID 09 | 15.312 | 501 | 0.33 | 0.4 | 0.22 | 0.53 | 0.09 | 1.02 | sm |
| 14 | nucleusID 10 | 16.447 | 569 | 0.75 | 0.87 | 0.58 | 1.08 | 0.29 | 1.81 | sm |
| 14 | nucleusID 11 | 27.931 | 1461 | 0.54 | 0.54 | 0.36 | 0.66 | 0.14 | 0.97 | md |
| 14 | nucleusID 13 | 48.241 | 3867 | 0.69 | 0.85 | 0.47 | 1.01 | 0.11 | 2.33 | lg |
| 14 | nucleusID 15 | 62.348 | 6106 | 2.43 | 2.5 | 1.54 | 3.18 | 0.24 | 5.69 | lg |
| 14 | nucleusID 19 | 34.346 | 2112 | 1.31 | 1.66 | 0.62 | 2.39 | 0.14 | 4.83 | md |
| 14 | nucleusID 20 | 100.345 | 14247 | 2.41 | 2.45 | 1.84 | 2.9 | 0.77 | 5.39 | lg |
| 14 | nucleusID 21 | 12.016 | 325 | 2.92 | 2.77 | 2.22 | 3.26 | 1.3 | 4.24 | sm |

Table 1.29: Information on nuclei included in the calculations movie 11-14

##### 1.2.21 Supplemental Information 21 - Actin-Stromule-Contact-Sites

**Supported Figure: 8E** This section provides additional examples supporting the observation that stromule tips are frequently found in close proximity to nucleus-associated actin (main manuscript Figure 8E), based on triple-marker transgenic lines simultaneously labeling actin (eGFP::ABD2::eGFP), nuclei (NLS-SV40:mKOK), and the plastid stroma (SSU:E2-Crimson). Figure 1.30 on the next page shows four additional examples (S1–S4) in which stromule tips are positioned adjacent to actin filaments at or near the nuclear surface, with individual fluorescence channels separated to allow clear assessment of the spatial relationship between actin, the nucleus, and the stromule tip. Together with the examples shown in Supplemental Information 22 and Figure 8E, these observations support actin as a likely anchor point for stromule tips at the nucleus.

Figure 1.30: 4 additional supplementary examples for situations where stromule tips can be found in close proximity of nucleus associated actin. This figure separates the individual channels. A-C = Example S1 (b and c magnified part of panel B and C), D-E = Example S2 (d and e magnified part of panel D and E), F-H = Example S3 (f and g magnified part of panel F and G), I-J = Example S4; Fluorescence color code: A-C: blue = chlorophyll a, red = ssu:E2Crimson (plastid stroma), cyan = NLSSV40:mKOK, scale bar corresponds in A-H to 1 $\mu\text{m}$ and in I-J to 5 $\mu\text{m}$.

##### 1.2.22 Supplemental Information 22 - Actin-Stromule-Contact-Sites (continued)

**Supported Figure: 8E** This section provides six further examples (Figure 1.31) of stromule tips found in close proximity to nucleus-associated actin, imaged in the same triple-marker transgenic lines described in Supplemental Information 21 and Supplemental Methods 3. Across these examples, actin filaments are consistently observed at or near the contact point between the stromule tip and the nuclear surface.

**Figure 1.31:** 6 more supplementary examples for situations where stromule tips can be found in close proximity of nucleus associated actin. A-F example 1 to 6. Colors have been optimized for good contrast for each example, the respective colors correspond to the following cell structures: A) green = actin, purple = nucleus, red = plastid stroma; B) green = actin, blue = nucleus, purple = chlorophyll, red = plastid stroma; C) green = actin, purple = nucleus, red = plastid stroma; D) gray = actin, purple = nucleus, red = plastid stroma; E) green = actin, blue = nucleus, red = plastid stroma; F) upper panel: green = actin, cyan = nucleus, red = plastid stroma; lower panel: purple = nucleus, red = plastid stroma

##### 1.2.23 Supplemental Information 23 - Microtubule-Stromule-Contact-Sites

**Supported Figure: 8F** This section provides four examples illustrating the spatial relationship between microtubules and stromules at the nucleus (main manuscript Figure 8F), based on triple-marker transgenic lines simultaneously labeling microtubules (eGFP::MAP4), the plastid stroma including stromules (SSU:E2-Crimson), and nuclei (NLS-SV40:mKOK). Figure 1.31 shows four examples (S1-S4) in which microtubules (green), plastid stroma and stromules (red), and the nucleus (blue) are shown, with fluorescence channels partially separated to allow clear assessment of the spatial relationship between microtubules, the nucleus, and the stromule tip. In contrast to the actin examples shown in Supplemental Information 25 and 26, microtubules were only rarely found in direct contact (no visual gap between stromule tip and microtubule) with stromule tips near the nuclear surface, supporting the conclusion that microtubules do not serve as the primary anchor for stromules within the SPZ.

##### 1.2.24 Supplemental Information 24 - Impact of Oryzalin on stromules in the SPZ

**Supported Figure: 8X**

Figure 1.32: Four examples illustrating the spatial relationship between microtubules, stromules, and the nucleus. For each example, the upper panel shows all three channels merged, and the lower panel shows only the plastid stroma/stromules and nucleus channels, allowing assessment of stromule tip position relative to the nucleus independent of the microtubule signal; the two panels of each example are separated by a dashed line, with solid lines separating the four examples. A–B = Example S1, C–D = Example S2, E–F = Example S3, G–H = Example S4. Fluorescence color code: green = eGFP::MAP4 (microtubules), red = SSU:E2-Crimson (plastid stroma/stromules), blue = NLS-SV40:mKOK (nucleus). Scale bars are indicated in each panel.

**Experimental design** Stromule frequency was assessed following treatment with oryzalin, a microtubule-depolymerizing drug, compared to a mock-treated control. Leaves were treated with 100 μM oryzalin (from a 50 mM stock in DMSO; final DMSO concentration = 0.2%) for 2 h, with mock-treated controls receiving the corresponding DMSO concentration (0.2%) alone. Following incubation, single time-point z-stacks were captured using a Leica Stellaris 8 confocal microscope. For each treated and mock-treated sample, the total number of plastids (npltotal) and the number of plastids displaying stromules (nplstr) were recorded per nucleus, across four independent images per condition. To account for potential effects of cell size on stromule formation, cells were classified into three size categories based on cross-sectional cell area: small ($\leq 1000 \mu\text{m}^2$), medium ($1000\text{--}3000 \mu\text{m}^2$), and large ($>3000 \mu\text{m}^2$).

**Pooling and stromule frequency calculation** Because stromule frequency (SF) values calculated at the level of individual nuclei would yield only a small number of discrete possible values per nucleus (e.g., $3/4$, $4/5$), plastid counts were first summed across all nuclei within the same image and cell size category. Stromule frequency was then calculated per image as: $\text{SF} = (\text{sum of nplstr}) / (\text{sum of npltotal})$. This produced one SF value per image per cell size category ($n = 4$ images per treatment per size category), which served as the unit of statistical comparison (1.30).

| Cell size category | Treatment | Image ID | n nuclei | npltotal | nplstr | SF |
| --- | --- | --- | --- | --- | --- | --- |
| Small | mock | At114_line-1_P1_S1 | 18 | 47 | 21 | 0.447 |
| Small | mock | At114_line-1_P1_S2 | 14 | 38 | 20 | 0.526 |
| Small | mock | At114_line-3_P5_S2 | 23 | 110 | 75 | 0.682 |
| Small | mock | At114_line-8_P4_S1 | 10 | 26 | 20 | 0.769 |
| Small | ory | At114_line-1_P1_S1 | 27 | 71 | 42 | 0.592 |
| Small | ory | At114_line-1_P1_S2 | 20 | 46 | 38 | 0.826 |
| Small | ory | At114_line-3_P5_S1 | 6 | 19 | 9 | 0.474 |
| Small | ory | At114_line-3_P5_S2 | 5 | 13 | 10 | 0.769 |
| Medium | mock | At114_line-1_P1_S1 | 2 | 8 | 7 | 0.875 |
| Medium | mock | At114_line-1_P1_S2 | 6 | 21 | 19 | 0.905 |
| Medium | mock | At114_line-3_P5_S2 | 8 | 56 | 40 | 0.714 |
| Medium | mock | At114_line-8_P4_S1 | 7 | 20 | 20 | 1.000 |
| Medium | ory | At114_line-1_P1_S1 | 4 | 14 | 12 | 0.857 |
| Medium | ory | At114_line-1_P1_S2 | 4 | 15 | 15 | 1.000 |
| Medium | ory | At114_line-3_P5_S1 | 8 | 24 | 15 | 0.625 |
| Medium | ory | At114_line-3_P5_S2 | 7 | 22 | 20 | 0.909 |
| Large | mock | At114_line-1_P1_S1 | 2 | 8 | 6 | 0.750 |
| Large | mock | At114_line-1_P1_S2 | 3 | 14 | 12 | 0.857 |
| Large | mock | At114_line-3_P5_S2 | 1 | 9 | 8 | 0.889 |
| Large | mock | At114_line-8_P4_S1 | 2 | 5 | 5 | 1.000 |
| Large | ory | At114_line-1_P1_S1 | 1 | 3 | 3 | 1.000 |
| Large | ory | At114_line-1_P1_S2 | 2 | 9 | 8 | 0.889 |
| Large | ory | At114_line-3_P5_S1 | 3 | 14 | 11 | 0.786 |
| Large | ory | At114_line-3_P5_S2 | 3 | 11 | 11 | 1.000 |

Table 1.30: Image-level pooled plastid counts and resulting stromule frequency (SF) for mock and oryzalin-treated samples, by cell size category. Plastid counts were summed across all nuclei within the same image and cell size category prior to calculating SF.

**Statistical test** For each cell size category, stromule frequency was compared between mock and oryzalin-treated samples using a two-sided Wilcoxon rank-sum test (Mann-Whitney U test). This non-parametric test was chosen because of the small sample size per group ($n = 4$) and because stromule frequency is a bounded proportion, for which standard parametric assumptions (e.g., normality) cannot be reliably verified at this sample size. Exact p-values could not be computed in the presence of tied values, so the normal approximation with continuity correction was used in these cases (default behavior of `wilcox.test()` in R).

Because three tests were performed (one per cell size category), raw p-values were adjusted for multiple comparisons using the Benjamini-Hochberg (BH) procedure to control the false discovery rate ( 1.31).

| Cell size category | W statistic | n (mock) | n (ory) | p (raw) | p (BH-adjusted) |
| --- | --- | --- | --- | --- | --- |
| Small | 5.5 | 4 | 4 | 0.561 | 0.842 |
| Medium | 8.5 | 4 | 4 | 1.000 | 1.000 |
| Large | 5.5 | 4 | 4 | 0.552 | 0.842 |

Table 1.31: Wilcoxon rank-sum test results comparing stromule frequency between mock and oryzalin-treated samples, by cell size category. P-values were adjusted for multiple comparisons across the three size categories using the Benjamini-Hochberg procedure.

**Interpretation** Oryzalin treatment did not significantly alter stromule frequency relative to mock-treated controls in any of the three cell size categories tested (all BH-adjusted $p > 0.05$). The medium-sized cell category showed near-complete overlap in stromule frequency between treatment groups ($p = 1.0$), while small and large cell categories showed comparable, non-significant trends.

##### 1.2.25 Supplemental Information 25 - Impact of Cytochalasin D on stromules in the SPZ

**Supported Figure: 8W**

**Experimental design** Stromule frequency was assessed following treatment with cytochalasin D, an actin-depolymerizing drug, compared to a DMSO-treated control. Leaves were treated with 4 $\mu\text{M}$ Cytochalasin D (CTD; from a 50 mM stock in DMSO; final DMSO concentration = 0.1%) for 2 h, with control samples receiving the corresponding DMSO concentration (0.1%) alone. Following incubation, single time-point z-stacks were captured using a Leica Stellaris 8 confocal microscope. For each treated and control sample, the total number of plastids (npl) and the number of plastids displaying stromules (nplstr) were recorded per nucleus, across 23 (cytochalasin D) and 25 (DMSO) independent images. To account for potential effects of cell size on stromule formation, cells were classified into three size categories based on cross-sectional cell area: small ($\leq 1000 \mu\text{m}^2$), medium ($1000\text{--}3000 \mu\text{m}^2$), and large ($> 3000 \mu\text{m}^2$).

**Pooling and stromule frequency calculation** Because stromule frequency (SF) values calculated at the level of individual nuclei would yield only a small number of discrete possible values per nucleus (e.g., $3/4$, $4/5$), plastid counts were first summed across all nuclei within the same image and cell size category. Stromule frequency was then calculated per image as: $\text{SF} = (\text{sum of nplstr}) / (\text{sum of npltotal})$. This produced one SF value per image per cell size category. One image/size-class combination (DMSOP4-1, medium category) had zero total plastids observed in that size class, resulting in an undefined (0/0) ratio; this data point was excluded from the analysis. $\text{SF} = (\text{sum of nplstr}) / (\text{sum of npl})$

| Cell size category | Treatm. | Image ID (File) | n nuclei | npl | nplstr | SF |
| --- | --- | --- | --- | --- | --- | --- |
| Small | ctdp | Ctdp3-1 | 7 | 21 | 0 | 0.000 |
| Small | ctdp | Ctdp3-2 | 12 | 32 | 0 | 0.000 |
| Small | ctdp | Ctdp3-3 | 1 | 4 | 0 | 0.000 |
| Small | ctdp | Ctdp4-1 | 10 | 19 | 0 | 0.000 |
| Small | ctdp | Ctdp4-2 | 17 | 49 | 0 | 0.000 |
| Small | ctdp | Ctdp4-3 | 5 | 13 | 0 | 0.000 |
| Small | ctdp | Ctdp6-2 | 12 | 26 | 0 | 0.000 |
| Small | ctdp | Ctdp9-1 | 4 | 3 | 0 | 0.000 |
| Small | ctdp | Ctdp9-2 | 4 | 10 | 0 | 0.000 |
| Small | ctdp | Ctdp9-3 | 13 | 14 | 0 | 0.000 |
| Small | ctdp | ctdp 1-1 | 7 | 17 | 0 | 0.000 |
| Small | ctdp | ctdp 1-2 | 6 | 13 | 0 | 0.000 |
| Small | ctdp | ctdp1-3 | 8 | 15 | 0 | 0.000 |
| Small | ctdp | ctdp2-1 | 7 | 16 | 1 | 0.062 |
| Small | ctdp | ctdp2-2 | 9 | 20 | 0 | 0.000 |
| Small | ctdp | ctdp2-3 | 7 | 20 | 0 | 0.000 |
| Small | ctdp | ctdp5-1 | 6 | 14 | 0 | 0.000 |
| Small | ctdp | ctdp5-2 | 3 | 11 | 0 | 0.000 |
| Small | ctdp | ctdp5-3 | 7 | 22 | 0 | 0.000 |
| Small | ctdp | ctdp6-1 | 7 | 7 | 0 | 0.000 |
| Small | ctdp | ctdp8-1 | 14 | 28 | 5 | 0.179 |
| Small | ctdp | ctdp8-2 | 7 | 17 | 0 | 0.000 |
| Small | ctdp | ctdp8-3 | 6 | 15 | 0 | 0.000 |
| Small | dms0 | DMSOP1-1 | 10 | 22 | 0 | 0.000 |
| Small | dms0 | DMSOP1-2 | 6 | 12 | 0 | 0.000 |
| Small | dms0 | DMSOP1-3 | 7 | 17 | 0 | 0.000 |
| Small | dms0 | DMSOP2-1 | 3 | 8 | 0 | 0.000 |
| Small | dms0 | DMSOP2-2 | 7 | 13 | 0 | 0.000 |
| Small | dms0 | DMSOP2-3 | 11 | 29 | 0 | 0.000 |
| Small | dms0 | DMSOP3-1 | 3 | 9 | 0 | 0.000 |
| Small | dms0 | DMSOP3-2 | 8 | 25 | 1 | 0.040 |
| Small | dms0 | DMSOP3-3 | 6 | 15 | 0 | 0.000 |
| Small | dms0 | DMSOP4-1 | 10 | 10 | 0 | 0.000 |
| Small | dms0 | DMSOP4-2 | 18 | 28 | 1 | 0.036 |
| Small | dms0 | DMSOP4-3 | 32 | 74 | 0 | 0.000 |
| Small | dms0 | DMSOP4-4 | 19 | 48 | 0 | 0.000 |
| Small | dms0 | DMSOP5-1 | 4 | 6 | 0 | 0.000 |
| Small | dms0 | DMSOP5-2 | 8 | 15 | 0 | 0.000 |
| Small | dms0 | DMSOP5-3 | 6 | 14 | 0 | 0.000 |
| Small | dms0 | DMSOP6-1 | 10 | 15 | 0 | 0.000 |
| Small | dms0 | DMSOP6-2 | 18 | 32 | 0 | 0.000 |
| Small | dms0 | DMSOP6-3 | 11 | 22 | 0 | 0.000 |
| Small | dms0 | DMSOP8-1 | 5 | 13 | 0 | 0.000 |
| Small | dms0 | DMSOP8-2 | 10 | 30 | 1 | 0.033 |
| Small | dms0 | DMSOP8-3 | 9 | 17 | 1 | 0.059 |
| Small | dms0 | DMSOP9-1 | 4 | 8 | 0 | 0.000 |
| Small | dms0 | DMSOP9-2 | 9 | 13 | 0 | 0.000 |
| Small | dms0 | DMSOP9-3 | 4 | 8 | 0 | 0.000 |

Table 1.32: Image-level pooled plastid counts and resulting stromule frequency (SF) for DMSO and cytochalasin D (ctdp) treated samples, for small cells. Plastid counts were summed across all nuclei within the same image (File) and cell size category prior to calculating SF. n nuclei indicates the number of nuclei contributing to each pooled value. One row (DMSOP4-1, medium) with zero total plastids observed (SF undefined) has been excluded.

| Cell size category | Treatm. | Image ID (File) | n nuclei | npl | nplstr | SF |
| --- | --- | --- | --- | --- | --- | --- |
| Medium | ctdp | Ctdp3-1 | 13 | 47 | 0 | 0.000 |
| Medium | ctdp | Ctdp3-2 | 7 | 29 | 0 | 0.000 |
| Medium | ctdp | Ctdp3-3 | 9 | 33 | 0 | 0.000 |
| Medium | ctdp | Ctdp4-1 | 13 | 43 | 1 | 0.023 |
| Medium | ctdp | Ctdp4-2 | 12 | 37 | 0 | 0.000 |
| Medium | ctdp | Ctdp4-3 | 3 | 13 | 1 | 0.077 |
| Medium | ctdp | Ctdp6-2 | 11 | 29 | 0 | 0.000 |
| Medium | ctdp | Ctdp9-1 | 2 | 3 | 0 | 0.000 |
| Medium | ctdp | Ctdp9-2 | 4 | 6 | 0 | 0.000 |
| Medium | ctdp | Ctdp9-3 | 9 | 24 | 0 | 0.000 |
| Medium | ctdp | ctdp 1-1 | 8 | 22 | 0 | 0.000 |
| Medium | ctdp | ctdp 1-2 | 4 | 12 | 0 | 0.000 |
| Medium | ctdp | ctdp1-3 | 7 | 18 | 0 | 0.000 |
| Medium | ctdp | ctdp2-1 | 7 | 21 | 0 | 0.000 |
| Medium | ctdp | ctdp2-2 | 8 | 27 | 0 | 0.000 |
| Medium | ctdp | ctdp2-3 | 9 | 29 | 0 | 0.000 |
| Medium | ctdp | ctdp5-1 | 9 | 30 | 0 | 0.000 |
| Medium | ctdp | ctdp5-2 | 5 | 17 | 0 | 0.000 |
| Medium | ctdp | ctdp5-3 | 8 | 24 | 0 | 0.000 |
| Medium | ctdp | ctdp6-1 | 9 | 21 | 0 | 0.000 |
| Medium | ctdp | ctdp8-1 | 15 | 38 | 6 | 0.158 |
| Medium | ctdp | ctdp8-2 | 4 | 9 | 0 | 0.000 |
| Medium | ctdp | ctdp8-3 | 9 | 33 | 0 | 0.000 |
| Medium | dms0 | DMSOP1-1 | 9 | 23 | 3 | 0.130 |
| Medium | dms0 | DMSOP1-2 | 6 | 21 | 4 | 0.190 |
| Medium | dms0 | DMSOP1-3 | 13 | 35 | 8 | 0.229 |
| Medium | dms0 | DMSOP2-1 | 8 | 23 | 1 | 0.043 |
| Medium | dms0 | DMSOP2-2 | 7 | 18 | 3 | 0.167 |
| Medium | dms0 | DMSOP2-3 | 5 | 15 | 3 | 0.200 |
| Medium | dms0 | DMSOP3-1 | 6 | 20 | 0 | 0.000 |
| Medium | dms0 | DMSOP3-2 | 13 | 39 | 5 | 0.128 |
| Medium | dms0 | DMSOP3-3 | 12 | 44 | 0 | 0.000 |
| Medium | dms0 | DMSOP4-2 | 15 | 59 | 1 | 0.017 |
| Medium | dms0 | DMSOP4-3 | 11 | 27 | 0 | 0.000 |
| Medium | dms0 | DMSOP4-4 | 11 | 32 | 2 | 0.062 |
| Medium | dms0 | DMSOP5-1 | 5 | 17 | 2 | 0.118 |
| Medium | dms0 | DMSOP5-2 | 12 | 41 | 9 | 0.220 |
| Medium | dms0 | DMSOP5-3 | 4 | 13 | 0 | 0.000 |
| Medium | dms0 | DMSOP6-1 | 9 | 19 | 0 | 0.000 |
| Medium | dms0 | DMSOP6-2 | 8 | 21 | 4 | 0.190 |
| Medium | dms0 | DMSOP6-3 | 14 | 37 | 2 | 0.054 |
| Medium | dms0 | DMSOP8-1 | 12 | 30 | 3 | 0.100 |
| Medium | dms0 | DMSOP8-2 | 4 | 8 | 1 | 0.125 |
| Medium | dms0 | DMSOP8-3 | 10 | 25 | 0 | 0.000 |
| Medium | dms0 | DMSOP9-1 | 5 | 16 | 0 | 0.000 |
| Medium | dms0 | DMSOP9-2 | 3 | 9 | 0 | 0.000 |
| Medium | dms0 | DMSOP9-3 | 5 | 12 | 0 | 0.000 |

Table 1.33: Image-level pooled plastid counts and resulting stromule frequency (SF) for DMSO and cytochalasin D (ctdp) treated samples, for medium cells. Plastid counts were summed across all nuclei within the same image (File) and cell size category prior to calculating SF. n nuclei indicates the number of nuclei contributing to each pooled value. One row (DMSOP4-1, medium) with zero total plastids observed (SF undefined) has been excluded.

| Cell size category | Treatm. | Image ID (File) | n nuclei | npl | nplstr | SF |
| --- | --- | --- | --- | --- | --- | --- |
| Large | ctdp | Ctdp3-1 | 12 | 55 | 4 | 0.073 |
| Large | ctdp | Ctdp3-2 | 11 | 54 | 4 | 0.074 |
| Large | ctdp | Ctdp3-3 | 12 | 51 | 3 | 0.059 |
| Large | ctdp | Ctdp4-1 | 7 | 29 | 0 | 0.000 |
| Large | ctdp | Ctdp4-2 | 8 | 40 | 3 | 0.075 |
| Large | ctdp | Ctdp4-3 | 4 | 23 | 2 | 0.087 |
| Large | ctdp | Ctdp6-2 | 11 | 44 | 0 | 0.000 |
| Large | ctdp | Ctdp9-1 | 7 | 8 | 1 | 0.125 |
| Large | ctdp | Ctdp9-2 | 4 | 10 | 0 | 0.000 |
| Large | ctdp | Ctdp9-3 | 6 | 18 | 1 | 0.056 |
| Large | ctdp | ctdp 1-1 | 8 | 32 | 6 | 0.188 |
| Large | ctdp | ctdp 1-2 | 9 | 40 | 8 | 0.200 |
| Large | ctdp | ctdp1-3 | 10 | 41 | 1 | 0.024 |
| Large | ctdp | ctdp2-1 | 9 | 30 | 7 | 0.233 |
| Large | ctdp | ctdp2-2 | 6 | 24 | 3 | 0.125 |
| Large | ctdp | ctdp2-3 | 13 | 63 | 4 | 0.063 |
| Large | ctdp | ctdp5-1 | 7 | 27 | 7 | 0.259 |
| Large | ctdp | ctdp5-2 | 6 | 27 | 4 | 0.148 |
| Large | ctdp | ctdp5-3 | 6 | 32 | 1 | 0.031 |
| Large | ctdp | ctdp6-1 | 13 | 55 | 4 | 0.073 |
| Large | ctdp | ctdp8-1 | 14 | 24 | 3 | 0.125 |
| Large | ctdp | ctdp8-2 | 3 | 9 | 1 | 0.111 |
| Large | ctdp | ctdp8-3 | 11 | 34 | 2 | 0.059 |
| Large | dms0 | DMSOP1-1 | 10 | 45 | 17 | 0.378 |
| Large | dms0 | DMSOP1-2 | 10 | 43 | 10 | 0.233 |
| Large | dms0 | DMSOP1-3 | 7 | 29 | 19 | 0.655 |
| Large | dms0 | DMSOP2-1 | 10 | 36 | 7 | 0.194 |
| Large | dms0 | DMSOP2-2 | 6 | 21 | 5 | 0.238 |
| Large | dms0 | DMSOP2-3 | 4 | 13 | 4 | 0.308 |
| Large | dms0 | DMSOP3-1 | 14 | 52 | 11 | 0.212 |
| Large | dms0 | DMSOP3-2 | 12 | 40 | 20 | 0.500 |
| Large | dms0 | DMSOP3-3 | 12 | 40 | 6 | 0.150 |
| Large | dms0 | DMSOP4-2 | 8 | 34 | 4 | 0.118 |
| Large | dms0 | DMSOP4-3 | 14 | 71 | 14 | 0.197 |
| Large | dms0 | DMSOP4-4 | 15 | 59 | 10 | 0.169 |
| Large | dms0 | DMSOP5-1 | 13 | 50 | 18 | 0.360 |
| Large | dms0 | DMSOP5-2 | 8 | 33 | 14 | 0.424 |
| Large | dms0 | DMSOP5-3 | 3 | 9 | 4 | 0.444 |
| Large | dms0 | DMSOP6-1 | 15 | 40 | 12 | 0.300 |
| Large | dms0 | DMSOP6-2 | 6 | 19 | 5 | 0.263 |
| Large | dms0 | DMSOP6-3 | 10 | 30 | 3 | 0.100 |
| Large | dms0 | DMSOP8-1 | 8 | 20 | 3 | 0.150 |
| Large | dms0 | DMSOP8-2 | 9 | 30 | 4 | 0.133 |
| Large | dms0 | DMSOP8-3 | 3 | 12 | 4 | 0.333 |
| Large | dms0 | DMSOP9-1 | 3 | 9 | 1 | 0.111 |
| Large | dms0 | DMSOP9-2 | 9 | 17 | 2 | 0.118 |
| Large | dms0 | DMSOP9-3 | 9 | 25 | 3 | 0.120 |

Table 1.34: Image-level pooled plastid counts and resulting stromule frequency (SF) for DMSO and cytochalasin D (ctdp) treated samples, for large cells. Plastid counts were summed across all nuclei within the same image (File) and cell size category prior to calculating SF. n nuclei indicates the number of nuclei contributing to each pooled value. One row (DMSOP4-1, medium) with zero total plastids observed (SF undefined) has been excluded.

**Statistical test** For each cell size category, stromule frequency was compared between DMSO and cytochalasin D-treated samples using a two-sided Wilcoxon rank-sum test (Mann-Whitney U test). A non-parametric test was used for consistency with the parallel analysis of the oryzalin dataset, and because stromule frequency is a bounded proportion for which parametric assumptions are not reliably verifiable. Exact p-values could not be computed in the presence of tied values (predominantly arising from the high proportion of $SF = 0$ values in the cytochalasin D group), so the normal approximation with continuity correction was used in these cases (default behavior of `wilcox.test()` in R). Because three tests were performed (one per cell size category), raw p-values were adjusted for multiple comparisons using the Benjamini-Hochberg (BH) procedure to control the false discovery rate.

| Cell size category | W statistic | n (dms) | n (ctdp) | p (raw) | p (BH-adjusted) |
| --- | --- | --- | --- | --- | --- |
| Small | 304.5 | 25 | 23 | 0.553 | 0.553 |
| Medium | 421.0 | 24 | 23 | 0.00044 | 0.00066 |
| Large | 482.5 | 24 | 23 | 0.0000116 | 0.0000347 |

Table 1.35: Wilcoxon rank-sum test results comparing stromule frequency between DMSO (control) and cytochalasin D (ctdp, actin-disrupting drug) treated samples, by cell size category. One image-x-size-class data point (DMSOP4-1, medium category) had zero total plastids observed and was excluded as the stromule frequency ratio was undefined. P-values were adjusted for multiple comparisons across the three size categories using the Benjamini-Hochberg procedure.

**Interpretation** Cytochalasin D treatment significantly reduced stromule frequency relative to DMSO-treated controls in medium-sized (BH-adjusted $p = 0.00066$) and large-sized (BH-adjusted $p = 0.0000347$) cells, but not in small-sized cells (BH-adjusted $p = 0.553$). In the medium and large categories, the cytochalasin D group showed predominantly zero or near-zero stromule frequencies, while the DMSO control group retained substantially higher and more variable stromule frequencies. This pattern is consistent with an actin-cytoskeleton-dependent mechanism of stromule formation or extension that becomes more apparent, or more required, in larger cells/plastid populations, while having comparatively little detectable effect in small cells. The absence of a significant effect in the small cell category may reflect a floor effect, since stromule frequencies in the small category were already low (close to zero) in both treatment groups, leaving little room for treatment-induced differences to be detected.
