## Supplementary material for "“Myosin-Driven Nuclear Movement Generates Stromules in *Arabidopsis thaliana*”": Video Legends

**Video 1 to 5**

Sample time-lapse data of upper epidermis cells illustrating plastid-nucleus-stromule dynamics in wildtype *A. thaliana*, including a detailed breakdown of synchronized versus asynchronous stromule formation relative to nuclear movement (Video 1 and 2), followed by direct comparisons of nuclear movement and stromule dynamics between wildtype and the myosin xi-i, wit1/2, and 4ko mutant lines (Video 3, 4 and 5).

**Video 1:** WT - In this video, we highlight the behavior of nuclei (red), plastids (green), and stromules (green) over a 90-minute time frame, which corresponds to the standard duration of the time-lapse experiments in this manuscript. In general, the nuclei can be seen oscillating in the center of the cell. The larger and more oblong the nucleus, and the larger the cell (as these are linked), the more pronounced this oscillating movement becomes. Plastids in the cell remain mostly stationary throughout the experiment. A subpopulation of plastids forms clusters in close proximity to the nucleus. These clusters often exhibit stromules directed toward the nucleus, with these stromules undergoing deformation (stretching and retracting), typically in synchronization with nuclear movement. This suggests a possible causal link between nuclear movement and stromule deformation. Despite this often-correlated behavior, some nucleus-associated plastids occasionally form stromules asynchronously to nuclear movement, suggesting an independent mechanism of formation. Additionally, stromule formation by plastids that are not part of the plastid-nucleus clusters can occasionally be observed, indicating that in these cases, there is no direct connection to nuclear movement. We zoom in on individual plastid-nucleus clusters to provide a clearer view: Cluster 1: Six plastids reside in the stromule-promoting zone around the oscillating nucleus. Two of these plastids display longer stromules, seemingly connected to the nucleus. Most stromule deformation events appear to be synchronized with nuclear movement. Cluster 2: In this smaller cell, three plastids form the plastid-nucleus complex. All plastids show stromule formation and deformation, seemingly in contact with the nucleus. Here, too, the majority of stromule formation/deformation events are synchronized with nuclear movement. Cluster 3: The nucleus movement in this cluster diverges somewhat from the previous examples, as the nucleus moves in a directed manner to the left, appearing to pull stromules out of the four plastids in the plastid-nucleus complex. Three of the plastids appear to follow the nucleus toward the left, while the rightmost plastid seems to lose its stromule contact with the nucleus. This leads to stromule retraction and the plastid stopping to follow the nucleus, suggesting that stromules attached near the nucleus may act as physical tethers. In summary: Most stromule deformation events (elongation and retraction) within the plastid-nucleus complex seem to be synchronized with nuclear movement. Time lapse experiment of wildtype *A. thaliana* expressing the pn marker. Images are z-projections of z-staks taken over time. Chlorophyll a = blue, FNR-eGFP = green, H2B:mcherry = red.

**Video 2:** Stromule formation coincides in most cases with nuclear movement but not in all cases (WT) - This video illustrates the relationship between nuclear movement and stromule formation that underlies the cor+/cor– categorization presented in Figure 6. The video opens with the full time-lapse sequence of the upper epidermis, showing the nucleus (red) oscillating within the cell while surrounding plastids (grey) remain largely stationary, before zooming in on four individual examples corresponding to Figure 6, panels A–D. Panel A: A stromule forms and elongates as the nucleus moves away from its associated plastids, with the stromule tips appearing to remain in contact with the nucleus surface throughout the movement, a possible direct anchor point. Panel B: A similar synchronized event, but here a visible gap persists between the stromule tip and the nucleus surface throughout the sequence, suggesting the stromule tracks nuclear movement without a stable direct attachment to the nucleus surface. Panel C: A plastid located outside the stromule-promoting zone, distant from the nucleus, forms and retracts a stromule independently of any nuclear movement nearby, an example of stromule formation unrelated to nuclear dynamics (cor–) outside the stromule promoting zone (SPZ). Panel D: Two nucleus-associated plastids within the SPZ form stromules asynchronously of nucleus movement, while the nucleus itself remains largely stationary stromules form, in this case even facing away from the nucleus, illustrating that not all stromule formation events near the nucleus are driven by nuclear displacement (cor–). In summary: while the majority of stromule formation events near the nucleus occur in synchrony with nuclear movement (cor+), a subset of events, both within and outside the SPZ, occur independently of nuclear displacement (cor–), indicating that multiple mechanisms of stromule formation coexist. Time-lapse experiment of wildtype *A. thaliana* expressing the pn marker. Images are z-projections of z-stacks taken over time. FNR-eGFP = grey, H2B:mCherry = red.

**Video 3:** WT vs. xi-i - Direct comparison of time-lapse data from an *A. thaliana* wildtype pn line and an *A. thaliana* *myosin xi-i* pn line reveals clear differences in nuclear movement and shape in *myosin xi-i* plants compared to the wildtype. In *myosin xi-i* plants, the nuclei of all cells, regardless of size, are round and not oblong or spindle-shaped as in the wildtype, which is a known phenotype of *myosin xi-i* plants originally described in root hair cells. Additionally, the amplitude of the normally strong oscillation of epidermal nuclei is severely reduced, and the nuclei appear to only ’wiggle.’ As a consequence of this reduced amplitude, plastids in the plastid-nucleus clusters still show some degree of nucleus-synchronized stromule stretching and retraction. However, the observed changes in stromule length are less pronounced and less frequent. Despite the reduction in nucleus-synchronized stromule formation, asynchronous stromule formation in the plastid-nucleus clusters, independent of nuclear movement, appears unaffected and can still be observed. As in the wildtype, plastids outside the plastid clusters can also form stromules independently of nuclear movement, suggesting that this mechanism also remains functional in the *myosin xi-i* mutant. In summary: Nucleus-movement-synchronized stromule formation is significantly reduced in the *myosin xi-i* mutant. The images shown are z-projections of z-stacks taken over time. Chlorophyll a = blue, FNR-eGFP = green, H2B:mcherry = red.

**Video 4:** WT vs. wit1/2 - Direct comparison of time-lapse data from an *A. thaliana* wildtype pn line and an *A. thaliana* *wit1/2* pn line reveals clear differences in nuclear movement and shape in *wit1/2* plants compared to the wildtype. In *wit1/2* plants, the nuclei of all cells, regardless of size, are round and not oblong or spindle-shaped as in the wildtype, a phenotype also described for this double mutant in root hair cells. Additionally, the amplitude of the normally strong oscillation of epidermal nuclei is severely reduced, and the nuclei appear to only ’wiggle’, as also seen for *myosin xi-i*. As a consequence of this reduced amplitude, plastids in the plastid-nucleus clusters still show some degree of nucleus-synchronized stromule stretching and retraction. However, the observed changes in stromule length are less pronounced and less frequent, as also seen in *myosin xi-i* mutants. Despite the reduction in nucleus-synchronized stromule formation, asynchronous stromule formation in the plastid-nucleus clusters, independent of nuclear movement, appears unaffected and can still be observed. As in the wildtype, plastids outside the plastid clusters can also form stromules independently of nuclear movement, suggesting that this mechanism remains functional in the *wit1/2* mutant. In summary: Nucleus-movement-synchronized stromule formation is significantly reduced in *wit1/2* mutants. The images shown are z-projections of z-stacks taken over time. Chlorophyll a = blue, FNR-eGFP = green, H2B:mcherry = red.

**Video 5:** WT vs. *4ko* - Direct comparison of time-lapse data from an *A. thaliana* wildtype pn line and a *4ko* pn line reveals clear differences in nuclear movement and shape in *4ko* plants compared to the wildtype. In *4ko* plants, the nuclei of all cells, regardless of size, are round and not oblong or spindle-shaped as in the wildtype. Unlike the *myosin* *xi-i* single and the *wit1/2* double mutant, the quadruple mutant shows almost no oscillating nuclear movement. The nuclei are stationary, with only minimal movement detectable. Furthermore, no stromule formation events synchronized with nuclear movement are observed. Only in rare cases, when the nucleus slowly drifts away from a plastid in the plastid-nucleus cluster, is a seeming attachment of the plastid to the nucleus periphery visible, along with slight plastid membrane deformation. This suggests that some plastid-nucleus attachment points, the potential source of stromule formation in the wildtype, may still exist. Nucleus-asynchronous stromule formation in the plastid-nucleus complex is also severely reduced. Occasional stromule formation outside the stromule-promoting zone indicates that this specific nucleus distant mechanism remains likely unaffected. In summary: Nuclear movement in the *4ko* mutant is severely reduced, and stromule formation in connection with the nucleus is effectively abolished. The images shown are z-projections of z-stacks taken over time. Chlorophyll a = blue, FNR-eGFP = green, H2B:mcherry = red.

**Video 6 to 8**

To directly test whether microtubules or actin filaments serve as the anchor for stromule tips at the nuclear periphery, three-dimensional reconstructions of the nucleus-cytoskeleton-plastid arrangement were generated. These reconstructions, utilizing a single marker expressing plant line, revealed that microtubules contact the nuclear envelope almost exclusively on the cell wall-facing side, with no microtubules observed at plastid-nucleus contact sites, a spatial arrangement inconsistent with a direct role in stromule anchoring at the nucleus. To demonstrate that this spatial relationship is not anecdotal but instead represents the majority of observed cases, a series of three-dimensional reconstructions documenting the relative positioning of microtubules and the nucleus has been compiled.

**Video 6:** Spatial organization of microtubules relative to the nucleus - Three-dimensional reconstruction showing the spatial organization of microtubules relative to the nucleus in an upper epidermis cell of a GFP:MAP4 *A. thaliana* (ecotype Columbia). Microtubules (MTs, visualized by eGFP:MAP4, green) are predominantly localized to the cell cortex, with MT-nucleus contact points occurring exclusively where the nuclear envelope meets the cortical MT network. No microtubules were observed at plastid-nucleus contact sites. Nuclear DNA was stained with DAPI (cyan/blue) and chlorophyll autofluorescence is shown in magenta.

**Video 7:** Spatial organization of microtubules relative to the nucleus - Three-dimensional reconstruction showing the spatial organization of microtubules relative to the nucleus in an upper epidermis cell of a GFP:MAP4 *A. thaliana* (ecotype Columbia). Microtubules (MTs, visualized by eGFP:MAP4, green) are predominantly localized to the cell cortex, with MT-nucleus contact points occurring exclusively where the nuclear envelope meets the cortical MT network. No microtubules were observed at plastid-nucleus contact sites. Nuclear DNA was stained with DAPI (cyan/blue), which also revealed mitochondrial DNA as small particles distributed near the cell periphery between the cortical MT network. Chlorophyll autofluorescence is shown in magenta.

**Video 8:** A rare instance of a microtubule traversing the nuclear surface - Three-dimensional reconstruction showing the spatial organization of microtubules relative to the nucleus in an upper epidermis cell of a GFP:MAP4 transgenic *A. thaliana* (ecotype Columbia). While microtubules (MTs, visualized by eGFP:MAP4, green) are predominantly localized to the cell cortex, this cell displays a rare instance of a single microtubule traversing the nuclear surface between the nucleus and vacuole. As in other cells, cortical MT-nucleus contact points occur exclusively where the nuclear envelope meets the cortical MT network, and no microtubules were observed at plastid-nucleus contact sites. Nuclear DNA was stained with DAPI (cyan/blue) and chlorophyll autofluorescence is shown in magenta.

**Video 9 to 11**

In contrast to microtubules, these three-dimensional reconstructions in a single actin marker expressing line revealed a fine actin mesh encasing the nucleus, with distinct filaments frequently observed sandwiched directly between plastids and the nuclear surface, a spatial arrangement consistent with actin serving as the anchor for stromule tips at the nucleus. To demonstrate that this spatial relationship is not anecdotal but instead represents the majority of observed cases, a series of three-dimensional reconstructions documenting the relative positioning of actin filaments and the nucleus has been compiled.

**Video 9:** Actin filaments at plastid-nuclear contact surfaces - Three-dimensional reconstruction of the actin cytoskeleton relative to the nucleus and plastids in an upper epidermis cell of a GFP:ABD2:GFP transgenic *A. thaliana* (ecotype Columbia). The actin network (visualized by GFP:ABD2:GFP, green) forms a fine mesh that encases the nucleus and extends to plastid-nuclear contact surfaces. Notably, distinct actin filaments are visible between plastids and the nucleus (white circle), where they appear sandwiched between these organelles. Nuclear DNA was stained with DAPI (blue) and chlorophyll autofluorescence is shown in magenta.

**Video 10:** Nucleus suspended on thick actin bundles in large cells - Three-dimensional reconstruction of the actin cytoskeleton relative to the nucleus and plastids in an upper epidermis cell of a GFP:ABD2:GFP transgenic *A. thaliana* (ecotype Columbia). The actin network (visualized by GFP:ABD2:GFP, green) forms a fine mesh that encases the nucleus. In large cells, the nucleus is frequently suspended on thick actin bundles that lift it away from the cell periphery. In these cases thicker actin bundles are also present along the nucleus surface (white circle). Nuclear DNA was stained with DAPI (blue) and chlorophyll autofluorescence is shown in magenta.

**Video 11:** Actin filaments at plastid-nuclear contact surfaces - Three-dimensional reconstruction of the actin cytoskeleton relative to the nucleus and plastids in an upper epidermis cell of a GFP:ABD2:GFP transgenic *A. thaliana* (ecotype Columbia). The actin network (visualized by GFP:ABD2:GFP, green) forms a fine mesh that encases the nucleus and extends to plastid-nuclear contact surfaces. Notably, distinct actin filaments are visible between plastids and the nucleus (white circle), where they appear sandwiched between these organelles. Nuclear DNA was stained with DAPI (blue) and chlorophyll autofluorescence is shown in magenta.

**Video 12 to 16**

The preceding reconstructions of the actin cytoskeleton (Videos 9–11) utilized revealed a fine actin mesh encasing the nucleus, with distinct filaments frequently observed at plastid-nuclear contact surfaces, consistent with actin serving as an anchor for stromule tips at the nucleus. However, these observations relied on single-marker lines and could not directly visualize stromule morphology simultaneously with actin and nuclear structure. To obtain direct evidence for stromule tip contact with nuclear-associated actin, triple-marker transgenic A. thaliana wild-type lines were generated to simultaneously visualize plastid morphology (SSU::E2-Crimson), the actin cytoskeleton (GFP::ABD2::GFP), and the nucleus (NLSS40::mKOK::GUS), enabling three-dimensional reconstruction of stromule-actin-nucleus arrangements directly. To demonstrate that the resulting observations are not anecdotal but instead represent the majority of observed cases, five three-dimensional reconstructions documenting stromule tip proximity to nuclear-associated actin have been compiled.

**Video 12:** Stromule nucleus-actin anchoring in a larger cell - Three-dimensional of the plastids outer shape (SSU::E2-Crimson), the actin cytoskeleton (GFP::ABD2::GFP) and the nucleus (NLSS40::mKOK::GUS) in an upper epidermis cell of transgenic *A. thaliana* (ecotype Columbia). The actin network (green) forms a fine mesh that encases the nucleus. Notably, a plastid stromule extends towards the nucleus, with its tip establishing contact with the actin network on the nuclear surface (circle). The nucleus is highlighted in blue, plastid morphology in red, and the actin network in green.

**Video 13:** Stromule actin anchoring in close nucleus proximity - Three-dimensional reconstruction of the plastids outer shape (SSU::E2-Crimson), the actin cytoskeleton (GFP::ABD2::GFP) and the nucleus (NLSS40::mKOK::GUS) in an upper epidermis cell of transgenic *A. thaliana* (ecotype Columbia). The actin network (green) forms a fine mesh that encases the nucleus. Notably, a plastid stromule extends towards the nucleus, with its tip establishing contact with the actin network, which is in this case not directly on the surface of the nucleus, but within the distance of the stromule promoting zone (arrow). The nucleus is highlighted in blue, plastid morphology in red, and the actin network in green.

**Video 14:** Stromule nucleus-actin anchoring in a larger cell - Three-dimensional reconstruction of the plastids outer shape (SSU::E2-Crimson), the actin cytoskeleton (GFP::ABD2::GFP) and the nucleus (NLSS40::mKOK::GUS) in an upper epidermis cell of transgenic *A. thaliana* (eco type Columbia). Similar to Video12, but here two plastids extend stromules towards the nucleus, with their tips establishing contact with the actin network (arrows). The nucleus is suspended by characteristic thick actin bundles. The nucleus is highlighted in blue, plastid morphology in red, the actin network in green and chlorophyll a autofluorescence of mesophyll plastids is shown in purple.

**Video 15:** Stromule initiation at the nucleus - Three-dimensional reconstruction of the plastids outer shape (SSU::E2-Crimson), the actin cytoskeleton (GFP::ABD2::GFP) and the nucleus (NLSS40::mKOK::GUS) in an upper epidermis cell of transgenic *A. thaliana* (eco type Columbia). One plastid at a greater distance is connected to the nucleus by a long stromule, whose tip is in contact with actin at the nucleus surface (upper arrow). Another plastid shows stromule initiation, likely in response to nuclear movement to the left of the field of view (lower arrow). The nucleus is highlighted in blue, plastid morphology in red, and the actin network in green.

**Video 16:** Stromule nucleus-actin anchoring in a larger cell - Three-dimensional reconstruction of the plastids outer shape (SSU::E2-Crimson), the actin cytoskeleton (GFP::ABD2::GFP) and the nucleus (NLSS40::mKOK::GUS) in an upper epidermis cell of transgenic *A. thaliana* (eco type Columbia). Similar to 12, one plastid at a greater distance is connected to the nucleus by a long stromule, whose tip is in contact with actin at the nucleus surface (upper arrow). The nucleus is highlighted in blue, plastid morphology in red, and the actin network in green.

**Video 17 to 18**

The preceding reconstructions established that stromule tips frequently make contact with nuclear-associated actin, identifying actin as the most likely anchor for stromules at the nucleus. However, these observations alone could not exclude the possibility that microtubules contribute similarly to stromule-nucleus contacts. To obtain comparable evidence for microtubules, triple-marker transgenic *A. thaliana* lines were generated to simultaneously visualize plastid morphology (SSU::E2-Crimson), microtubules (GFP::MAP4), and the nucleus (NLSS40::mKOK::GUS), enabling three-dimensional reconstruction of stromule-microtubule-nucleus arrangements directly. To demonstrate that the resulting observations are not anecdotal but instead represent the majority of observed cases, two three-dimensional reconstructions documenting stromule tip proximity to nuclear-associated microtubules have been compiled.

**Video 17:** Stromule tips show no microtubule contact, with a single rare exception - Three-dimensional reconstruction of the plastids’ outer shape (SSU::E2-Crimson), microtubules (GFP::MAP4), and the nucleus (NLSS40::mKOK::GUS) in an upper epidermis cell of transgenic *A. thaliana* (ecotype Columbia). Four plastids with stromules directed toward the nucleus, which sits atop the cortical microtubule array, are visible; however, none of their stromule tips show close proximity to a microtubule that would suggest physical contact (arrows, top and left). One plastid displays a thicker protrusion facing away from the nucleus, which is likewise not in contact with a microtubule (arrow, right). However, the opposite end of this same plastid shows an apparent association with a microtubule, the only such instance documented in this study (circle). The nucleus is highlighted in blue, plastid morphology in red, and microtubules in green.

**Video 18:** Apparent stromule-microtubule proximity resolved as no contact upon 3D rotation - Three-dimensional reconstruction of the plastids’ outer shape (SSU::E2-Crimson), microtubules (GFP::MAP4), and the nucleus (NLSS40::mKOK::GUS) in an upper epidermis cell of transgenic *A. thaliana* (ecotype Columbia). Two stromules are visible: one clearly lacks contact with a nearby microtubule (bottom arrow), while the other initially appears to contact a microtubule fragment from the starting viewing angle, but rotation of the three-dimensional reconstruction reveals a clear gap between the stromule tip and the nearby microtubule (middle-left persistent arrow), demonstrating that apparent contact in a single static view can be misleading. Additionally, two broader plastid protrusions are visible, one spreading across the nuclear surface at its tip (top-left arrow) and one extending away from the nucleus (right arrow), both of which are likewise clearly independent of microtubules. The nucleus is highlighted in blue, plastid morphology in red, and microtubules in green.

**Video 19 to 22**

To test directly whether actin or microtubules are required to maintain stromule attachment at the nucleus, plants were treated with the actin-depolymerizing drug cytochalasin D or the microtubule-depolymerizing drug oryzalin, alongside matched DMSO solvent controls, and stromule and nucleus dynamics were followed continuously by time-lapse imaging.

**Video 19:** Stromule and nucleus dynamics under DMSO control (CTD experiment) - Maximum intensity projection of a time-lapse experiment using a triple-marker line (plastids = SSU::E2-Crimson (red), nucleus = NLS::mKOK (purple), actin = eGFP::ABD2::eGFP (green)), captured immediately after DMSO treatment in the upper epidermis (approx. 1 h 45 min). A) Nuclei and stromules exhibit dynamic movement and reshaping, while actin filaments remain intact throughout the observation. Mesophyll plastids display robust light-avoidance movement, with actin filaments forming in association with chloroplast bodies. B) Close-up of selected nuclei, with arrows highlighting example stromules that continuously reshape in synchrony with nuclear movement and remain visible at the end of the observation. C) Channels showing only plastids and nuclei reveal active nuclear movement and stromule formation; arrows indicate stromules reshaping with nuclear movement, while some stromules also form independently, providing an example for the additional mechanism involved stromule formation. D) Overview of the imaged area in the same two channels, showing consistent nucleus movement and stromule formation events across all visible cells.

**Video 20:** Impact of cytochalasin D treatment on nucleus and stromule dynamics - Maximum intensity projection of a time-lapse experiment using a triple-marker line (plastids = SSU::E2-Crimson (red), nucleus = NLS::mKOK (purple), actin = eGFP::ABD2::eGFP (green)), captured immediately after CTD treatment in the upper epidermis (approx. 1 h 45 min). A) Initial frames show dynamic, green-labeled actin filaments and bundles; CTD treatment rapidly induces filament breakdown, progressing throughout the movie, and by the end most actin is depolymerized, though some filaments remain near nuclei. Mesophyll chloroplasts show no strong light-avoidance movement. B) Close-up of selected nuclei, with arrows indicating stromules retracting from nuclei over time. C) Channels displaying plastids and nuclei alone reveal stromule tip retraction and halted nuclear movement following actin disruption (arrows). D) Overview of the imaged area in the same two channels, showing consistent stromule retraction and halted nucleus movement across cells.

**Video 21:** Stromule and nucleus dynamics under DMSO control (oryzalin experiment) - Maximum intensity projection of a time-lapse experiment using a triple-marker line (plastids = SSU::E2-Crimson (red), nucleus = NLS::mKOK (purple), microtubules = eGFP::MAP4 (green)), captured immediately after DMSO treatment in the upper epidermis (approx. 1h 45min). A) Nuclei and stromules exhibit dynamic movement and reshaping, while microtubules remain intact and cortical throughout the observation. This is accompanied by typical growing and shrinking of microtubule filaments. B) Close-up of selected nuclei, with arrows highlighting example stromules that continuously reshape in synchrony with nuclear movement and remain visible at the end of the observation. C) Channels showing only plastids and nuclei reveal active nuclear movement and stromule formation, with stromules reshaping in step with nuclear displacement. D) Overview of the imaged area in the same two channels, showing consistent nucleus movement, stromule formation in all cells.

**Video 22:** Impact of oryzalin treatment on nucleus and stromule dynamics - Maximum intensity projection of a time-lapse experiment using a triple-marker line (plastids = SSU::E2-Crimson (red), nucleus = NLS::mKOK (purple), microtubules = eGFP::MAP4 (green)), captured immediately after oryzalin treatment in the upper epidermis, movie has been captured at the end of a 1.5 h incubation time and runs for 30 min. A) Despite the prolonged pre-incubation a strongly fragmented microtubule network is visible in the first frames; oryzalin continued to break down microtubules into smaller fragments and punctate structures, however small fragments remained. B) Close-up of selected nuclei, with arrows indicating stromules that continue to extend and retract in synchrony with nuclear movement despite microtubule disruption; nuclei can be seen gliding over the stationary microtubule fragments. C) Channels displaying plastids and nuclei alone reveal that nuclear movement and stromule dynamics, both nucleus-dependent (cor+) and nucleus-independent (cor-) events, persist unchanged despite the dramatic reduction of cortical microtubules. D) Overview of the imaged area, showing that nucleus movement and stromule frequency remain comparable to control conditions across cells.
